# Welcome to Hotel Hymenoptera: monitoring cavity-nesting bee and wasp distribution and their trophic interactions using community science and metabarcoding

**DOI:** 10.1101/2024.07.10.602935

**Authors:** Sage Handler, Katerina Coveny, Thomas Braukmann, Nigel E. Raine, Dirk Steinke

## Abstract

Essential ecosystem services are provided by many interactions, including plant-pollinator, predator-prey, and host-parasitoid. These services support food and natural systems through pollination and pest control, however they are challenging to qualify, and previous observational studies may underestimate their complexity. The cavity nesting Hymenoptera are a good example showing all these three interactions and they can be monitored using trap nests. For this study, trap nests were installed at schools across Canada by community scientists to investigate cavity-nesting bee and wasp distributions and interactions. DNA metabarcoding was used to identify the occupants and their food sources. New bee and wasp distributions were found that might be the result of previous under-sampling or recent range expansions. Detailed bipartite and tripartite networks describing species interactions suggest some novel bee, wasp, and parasite associations. These results encourage further investigation into these interactions using molecular methods as detailed range maps and networks provide information to natural historians and conservationists alike.

## Introduction

Ecosystems that are influenced by human development are often highly disrupted and host reduced levels of biodiversity (Elmqvist et al. 2003, Sánchez-Bayo and Wyckhuys 2019). Although many factors contribute to species declines and extinctions, anthropogenically driven landscape changes are seen as the largest drivers (Vanbergen and Initiative 2013, Sánchez-Bayo and Wyckhuys 2019). Both agricultural intensification and urbanization are landcover changes that alter natural habitat, both reducing and fragmenting the amount of accessible living space for many species. Unfortunately, the extent of these disturbed ecosystems is increasing and has been globally for decades. These disruptions have negative consequences on many of the ecosystem services we humans rely on, by not only disturbing the species but also their interactions with each other.

An additional consequence of increased urbanization is that it may benefit the establishment and persistence of non-native species over native species (Hernandez et al. 2009, Fitch et al. 2019, Kammerer et al. 2021, Gruver and Caradonna 2021). Non-native bees for instance are not only resilient against increasing levels of urban development, but they might benefit from it to a point of outcompeting native species (Fitch et al. 2019). Non-native species are capable of outcompeting native specialist bees since they are generalists, able to feed on a wide variety of food sources (Russo 2016, Fitch et al. 2019, Geslin et al. 2020). Globally, 16 non-native bee species have been purposefully introduced for agricultural purposes, including the European honeybee *Apis mellifera*, multiple bumblebee (*Bombus*) species, and some solitary bees including *Megachile rotundata* (Pitts-Singer and Cane 2011, Graystock et al. 2016, Russo et al. 2021). Additionally, over 65 non-native species were introduced unknowingly, either mistaken for a similar looking species or transported by accident (LeCroy et al. 2020). The majority (70%) of introduced bee species are cavity-nesting (Russo 2016), increasing the potential for competition with native cavity-nesting bees. Whether they were introduced purposefully or by accident, non-native bee species have the potential to harm native species due to their increased fitness in urban areas.

Solitary cavity-nesting Hymenoptera represent a unique community where multiple ecological guilds occur and interact in the form of bees (pollinators), wasps (predators and pollinators) and their prey, as well as parasitoids which target all the former (Farzan et al. 2017). Reproductive solitary cavity-nesting bee and wasp females occupy their own nests, within hollow, above-ground nesting sites such as pithy stems and wood-feeding beetle burrows (Krombein 1967), and construct linear brood cells from the rear of the tubular nest hole to the front (Farzan et al. 2017, MacIvor 2017). In non-parasitic species, each brood cell is separated by species-specific material such as leaves, soil, or tree sap and supplied with food. Cavity-nesting bees provision offspring with a combination of pollen and nectar called bee bread, while wasps provision their nests with arthropod prey, such as spiders, caterpillars, or aphids (Krombein 1967, Farzan et al. 2017, Hoffmann et al. 2018). Both cavity-nesting bees and wasps are hosts for parasitoid wasps which can lay their eggs within the previously constructed nests, either once the nest is completed or while the female nest provisioner is out foraging (Farzan et al. 2017, Osorio-Canadas et al. 2018). Many parasitoid wasps can alternatively inject their egg directly into the host larvae.

While bees are well known for their pollinating services (Klein et al. 2007, Potts et al. 2016), their close relatives, wasps, also provide an additional beneficial ecosystem service. Many wasps act as natural predators that can be used in integrative pest management (IPM) strategies to control insect pests on plants without the use of chemical insecticides (Harris 1994, Hoffmann et al. 2018, da Rocha-Filho et al. 2020, Brock et al. 2021). Predatory wasps consume nectar and pollen as adults but require arthropods, such as caterpillars or spiders, to provision their offspring (Hoffmann et al. 2018, Wang et al. 2019). In addition to biocontrol agents, wasps are considered important biodiversity indicators as they require multiple ecosystem resources and are sensitive to their loss (Turčinavičiene et al. 2014, Hoffmann et al. 2018). As higher trophic level organisms, wasps are more vulnerable to habitat change, making conservation strategies increasingly important (da Rocha-Filho et al. 2020).

Due to their nesting habits, cavity-nesting bees and wasps can be sampled in human-made nests. Human-made trap nests, also called bee hotels or nest boxes, mimic natural cavity-nesting sites of hollow plant stems and other holes (Staab et al. 2018). Trap nests may be made from bundled plant stems, holes drilled in wood, paper-based tubes, or a variety of other methods (MacIvor 2017). The first recorded use of trap nests to study cavity-nesting Hymenoptera was by Jean-Henri Fabre (Fabre 1914), and since then, many have used this strategy to study the natural history or ecosystem service potential of cavity-nesting species (Tasei et al. 1976, Bosch 1994, Aguiar and Garófalo 2004). Although trap nests are limited to detecting only solitary cavity-nesting species, they collect additional information on the ecology of these species. Unlike many other passive insect sampling strategies, such as pan traps and blue vane traps, which may detect species in the area, trap nests provide an indisputable way of knowing a species’ nesting location (Westphal et al. 2008, Staab et al. 2018). Trap nests are an extremely valuable method of monitoring interactions between species, such as parasitic relationships and plant-pollinator interactions (Staab et al. 2018). The pollen or prey stored as food for larvae in trap nests can be directly linked to the species of bee or wasp that collected it (Staab et al. 2018). This makes the trap nest strategy ideal for studying the interactions of solitary cavity-nesting bees and wasps, their food and prey, and their parasitoids. Although cavity-nesting bees represent only a small fraction of all solitary bee species, their abundance is highly correlated with the overall abundance of nest-provisioning bees (Staab et al. 2018, Mayr et al. 2020). Therefore, they are representative of local bee communities and can allow some extrapolation on how local conditions are affecting bee communities.

Many methods exist to study species interactions and movement, one more recently introduced is DNA barcoding. DNA barcoding is a molecular method of species identification that has been used for over two decades (Hebert et al. 2003). DNA metabarcoding uses a similar premise but identifies multiple species in a community sample (Cristescu 2014). Many studies have used DNA barcoding techniques to identify both insect and pollen specimens (for example Magnacca and Brown 2012, Bell et al. 2016, Bell et al. 2017, Gresty et al. 2018). In many insect taxa, taxonomic identification to species level is impossible due to cryptic diversity or incomplete specimens (Packer et al. 2009). The identification of pollen through DNA barcoding is popular since pollen is traditionally very difficult to identify using comparative morphological approaches for taxonomic identifications (Bell et al. 2016, Peel et al. 2019). Recently, DNA metabarcoding has been used to form plant-pollinator networks by collecting pollen from bees’ legs (Bell et al. 2017, Gresty et al. 2018). As DNA barcoding databases, such as the Barcode of Life Database (BOLD) and GenBank, continue to grow, more species level identifications become possible using this molecular technique.

To date, only one other study has used DNA barcoding of cavity-nesting bees and wasps and their pollen and prey, respectively, together (Dürrbaum et al. 2023). Our study uses a similar approach to investigate the interactions between solitary cavity nesting Hymenoptera across Canada. Combining a community science approach with DNA metabarcoding, we took an in-depth look at solitary cavity-nesting bee and wasp distributions and interactions. Specifically, we wanted to determine: 1) What are the distributions of solitary cavity-nesting bee species across Canada and what flowers are they using as sources of nectar and pollen to provision their offspring? (2) What are the distributions of solitary cavity-nesting wasp species across Canada and how are they interacting with their diverse prey and parasitoids? (3) How does local landcover (within 500 m of the trap nest) affect the distribution of solitary cavity-nesting bees and wasps?

## Methods

### Sample collection

Trap nests were constructed following a design that has been used successfully in past studies (MacIvor and Salehi 2014, MacIvor et al. 2014, MacIvor 2016). A 30 cm section of 10cm-diameter PVC pipe was used as housing, with one end cut on a 60° angle to provide weather protection. The other end of the pipe was covered with a PVC pipe cap. A 10cm diameter circular piece of Styrofoam was cut to fit into the PVC pipe. Ten holes each of three diameters (3.4 mm, 5.6 mm, 7.6 mm diameter) were punctured through the Styrofoam using screwdrivers. Bee paper tubes (Custom Paper Company, US) matching those diameters were then inserted into the cardboard after one end had been plugged with paper clay.

Sites Canada-wide were managed by participants in the Bees@Schools community science program (beesatschools.ca) at the University of Guelph. During the summers of 2019 and 2020, trap nests were distributed to elementary and high schools across Canada. In 2019, 88 trap nests were sent out and 42 were returned, and in 2020, 202 nests were distributed and 170 returned to Guelph (Figure 1). Trap nests were installed by teachers from early May to late August to allow phenological coverage for various nesting strategies and provide sufficient observation time for students. Teachers were instructed to install nests facing south at a height of 1-1.5 metres above the ground. They reported the orientation, height, latitude, and longitude of the trap nest, as well as provided photos of the installation to ensure correct use. School locations were mapped using ArcGIS Online with the 2015 Land Cover of Canada dataset (Natural Resources Canada 2015). The landcover categories include urban, cropland, forest cover, and wetland. Buffers of 500m, 1km, and 5km for each school location were made in ArcMap and exported to the web map. ArcMap’s zonal histogram tool analyzed the dominant landscape surrounding each school location by counting the number of pixels for each land cover category found within the buffer zones around the schools. The output histogram table was sorted by schools with the greatest number of cropland/forest/urban pixels within the 500m, 1km or 5km buffers and sites were grouped into categories based on dominant land cover type.

**Figure 1:**
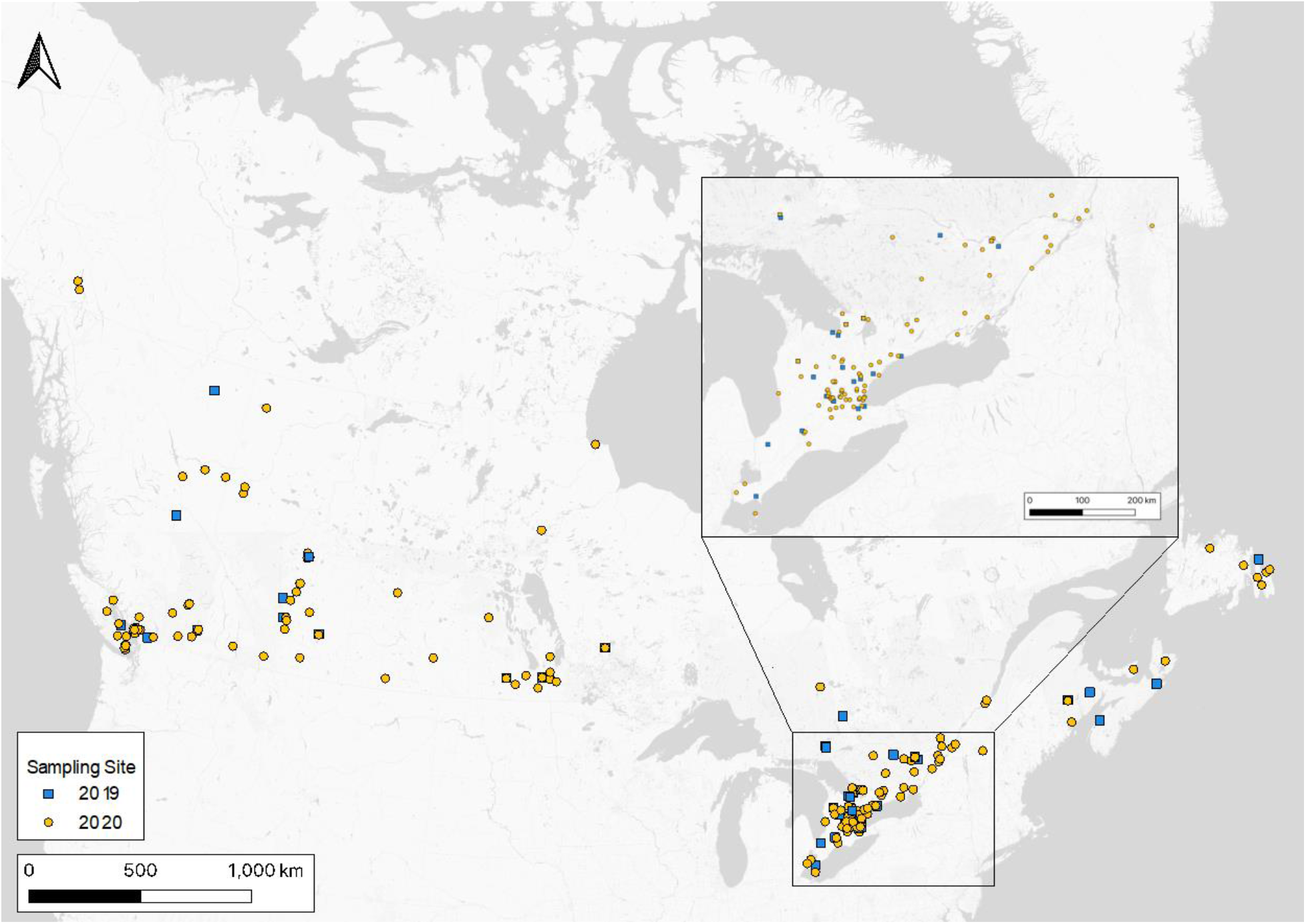
Map showing sampling sites across Canada in 2019 (42 sites, blue) and 2020 (170 sites, yellow).

### Sample processing

#### Sample extraction

Once nests were returned in late August-early September, cardboard nesting tubes were removed from the Styrofoam and separated by location on the nest face as quadrants (top or bottom and left or right). Cardboard tubes were stored at −20°C until further processing. Each cardboard tube represented a sample unit, resulting in multiple barcode samples available for a single site. Tubes were opened using a sterilized knife and forceps, and insect larvae, pollen, and prey were separated from leafy nesting material into two sterile tubes. Larvae samples were split into subsamples when necessary to ensure approximately equal tissue mass per DNA extraction. Leaf material was stored for further projects but not used for this study. The diameter of cardboard tube, type of nesting material (leaf, soil, tree sap, etc.), and number of larvae were recorded.

#### Molecular identification

All molecular lab work was conducted in a clean-room laboratory at the University of Guelph compliant with DNA processing standards. Work surfaces were cleaned with 1% bleach and 70% ethanol, followed by UVC-light radiation for 15 minutes. Pollen and larvae samples were mechanically masticated using a disposable pestle (VWR International, LLC., PA, USA) to break open the larval cuticle and release pollen. 40-100 mesh sand (ACRO Organics, Thermo Fisher Scientific Inc., MA, USA) and two 5mm stainless steel beads (IKA Works Inc., NC, USA) were added to further mechanically break-down tissue along with 900μL insect lysis buffer (700mM GuSCN, 30mM EDTA, 30mM Tris-HCL, 0.5% Triton X-100, 5% Tween-20, pH 8) and 100μL Proteinase K (QIAGEN, 20 mg/mL). Samples were shaken on a QIAGEN TissueLyser II at 28Hz for two minutes to further pulverize pollen. Samples were incubated on a rocking platform (OHAUS Corporation, NJ, USA) for 2.5 hours at 56°C, followed by one hour at 65°C. The resulting lysate was combined with binding buffer (6M GuSCN, 20mM EDTA, 10mM Tris-HCl, 4% Triton X-100) in a 2:1 ratio of buffer to lysate (1,200μL to 600μL), mixed briefly by vortexing, and centrifuged at 1,000 x g for 20 seconds. Lysate was transferred to a glass fibre column (Qiagen, Blood and tissue kit) in three aliquots of 600μL, centrifuged at 10,000 x g for two minutes between aliquots. 300μL of binding mix (150μL binding buffer, 150μL 96% ethanol) was added to the column followed by centrifuging at 10,000 x g for two minutes. The column was washed twice with 600μL of wash buffer (60% EtOH, 50mM NaCl, 10mM Tris-HCl, 0.5mM EDTA) and centrifuged at 10,000 x g for two minutes between washes. The column was then centrifuged at 10,000 x g for two minutes to dry. The glass fibre column was transferred to a clean 1.5mL microcentrifuge tube and 25μL of elution buffer (0.1M Tris-HCL, pH 8) was added. The sample was incubated at 56°C for one minute then centrifuged at 10,000 x g for two minutes. This was done twice for a total elution volume of 50μL. Each group of extractions contained one extraction negative control.

Gene fragments were amplified using a PCR reaction mix of 12.5μL of QIAGEN multiplex plus master mix, 1.25μL each of 10X PCR forward and reverse primer, 5μL of DNA template, and 5μL of nfH20. For bee, wasp, and prey identification, the BF3 + BR2 primer set was used (Supplementary Table 1; Elbrecht et al. 2019) to isolate the mitochondrial cytochrome c oxidase I (COI) gene. For plant identification, the rbcl1 + rbcLB primer set was used to isolate the ribulose bisphosphate carboxylase large chain (rbcL) gene (Supplementary Table 1; Palmieri et al. 2009, Little 2014). The following conditions were used for PCR: denaturation for 5 minutes at 95°C; 25 cycles of 30 s at 95°C, 45 s at 50°C, and 50 s at 72°C; final elongation for 5 min at 72°C; then held at 4°C. Samples were then indexed using a second PCR under the same conditions and the same reaction mix, though forward and reverse primers had custom tags and were sample specific (Supplementary Table 2; Elbrecht and Steinke 2019). Gel visualizations of tagged PCR products were done using 1% agarose gels on GelRed® Nucleic Acid Gel Stain (Biotium, CA, USA). Sample size ranges were optimized using SequalPrep™ Normalization Plate Kit (Invitrogen, Thermo Fisher Scientific Inc., MA, USA) according to the manufacturer’s instructions. Samples were pooled and SPRISelect (Beckman Coulter, CA, USA) was used to remove primer dimers with a ratio of 0.7X. Libraries were quantified using a Qubit Fluorometer with the Qubit dsDNA HS Assay Kit (Invitrogen, Thermo Fisher Scientific Inc., MA, USA) according to manufacturer’s instructions. Libraries from 2019 were sequenced using Illumina MiSeq at the University of Guelph Advanced Analytics Centre’s Genomics Facility. Libraries from 2020 were sequenced using one lane on a SP Chip on Illumina Novaseq at Sick Kids Hospital’s DNA Sequencing and Synthesis Facility.

### Data analysis

#### Data processing

Raw sequence data quality control was initially done using FastQC v0.11.8. Resulting libraries were analyzed using JAMP v0.69 (github.com/VascoElbrecht/JAMP). Sequences first underwent demultiplexing, then paired-end merging using Usearch v11.0.667 with fastq_pctid=75 (Edgar 2010). Primer sequences were trimmed using Cutadapt v1.18 with default settings (Martin 2011) and only sequences where primers were trimmed at each end were kept. Additionally, Cutadapt removed sequences outside lengths of 421 +/- 10 bp for COI sequences or 184 +/- 10 bp for rbcL sequences. Poor quality sequences were removed in Usearch with an expected error value of 1 (Edgar and Flyvbjerg 2015). Filtered reads for each sample were dereplicated and singletons removed. Each amplicon sequence variant (ASV) was compared to a custom database, with one each for Hymenoptera specifically, insects in general, and plants. Both the Hymenoptera and insect databases were based on the BOLD reference database (www.boldsystems.org; Ratnasingham and Hebert 2007). The custom plant database used data from Genbank (https://www.ncbi.nlm.nih.gov/genbank; Sayers et al. 2018). Insect detections (using COI) from 2019 and 2020 with less than 50 and 90 reads were removed for all analyses, representing 85% of all detections. The read threshold was increased in 2020 due to the increased detection sensitivity of Illumina Novaseq compared with Illumina Miseq. Hymenoptera detections with less than 1,000 reads (94% of all detections) were removed from interaction analyses to ensure their presence as true nesting occupants and not accidental visitors.

#### Statistical analyses

Unless otherwise mentioned, all statistical analyses were done using R statistical software v4.1.0 (R Core Team 2021). Spatial analyses were done using QGIS v3.18.3 (QGIS Development Team 2021).

Bipartite plant-pollinator and predator-prey networks were created using the R *bipartite* package (Dormann et al. 2008). Gephi was used to create a tripartite network (Bastian et al. 2009). Prior to creating networks, individual species were investigated to determine various life history traits. Insects were categorized based on trophic function as prey, pollinator, predator, or parasitoid (Krombein 1967, Krombein and Hurd 1979, Cane et al. 2007, Sheffield et al. 2011). Historical ranges were used to categorize each species as native or non-native. For taxa that could not be identified to species, a genus or family level identification was used. For these taxa, native range could not be identified, and in some cases neither could trophic function. These were subsequently left out of these analyses. Plants were analyzed at a minimum genus level since species-level identifications using rbcL have variable reliability (Braukmann et al. 2017). Network metrics including nestedness, links per bee species, and connectance were calculated using *bipartite*. Nestedness measures whether locations with low species richness are unique from locations with high species richness, and connectance is the proportion of possible links among species that are seen. Species range maps were made using historical data taken from GBIF using the R *rgbif* package (Chamberlain et al. 2022).

Local landscape features were determined by mapping site locations alongside the 2015 Land Cover of Canada dataset (Natural Resources Canada 2015). A 500-metre buffer was created for each site, then QGIS’s zonal histogram tool determined the number of pixels per buffer of each landcover type (nine types of forest, wetland, cropland, barren land, urban, water, snow/ice). These were converted into percent cover, and finally prominent landcover type was determined by which landcover type had the greatest area. The nine types of forest cover were amalgamated into one forest designation. Once landcovers were determined, redundancy discriminant analyses (RDAs) were run to investigate the relationship between community composition and land cover designation using the vegan package in R (Oksanen et al. 2008). The vegan package was also used to calculate bee and wasp community richness and abundance (Oksanen et al. 2008). Analyses of variance (ANOVAs) were run to determine whether communities were significantly different based on landcover types.

## Results

### General results

#### Metabarcoding success

High quality DNA was extracted from 887 samples corresponding to 638 cardboard tubes (Table 1). However, only 52% of 2019 tubes were successful compared with 91% of 2020 tubes. Reads generated by Illumina sequencing can be found in Table 2.

**Table 1:** Trap nest installation, occupation, and extraction in 2019 and 2020.

| Year | Nests installed | Nests occupied | Tubes occupied | Min / max tubes per site | Average tube occupancy per nest | Nests sequenced | Tubes sequenced |
| --- | --- | --- | --- | --- | --- | --- | --- |
| 2019 | 44 | 22 | 172 | 1 / 24 | 7.8 +/- 1.4 | 20 | 90 |
| 2020 | 170 | 93 | 604 | 1 / 27 | 6.5 +/- 0.6 | 93 | 548 |

**Table 2:** Trap nest sequencing reads generated by Illumina sequencing prior to and post filtering for COI and rbcL in both sampling years (2019 and 2020)

| Year | Marker | Unfiltered Reads | Average Reads Post-Filtering per Sample | Total Reads Post-Filtering |
| --- | --- | --- | --- | --- |
| 2019 | COI | 22,048,123 | 49,290 | 7,097,815 |
|  | rbcL | 17,179,971 | 56,982 | 8,091,722 |
| 2020 | COI | 574,568,330 | 75,612 | 57,842,823 |
|  | rbcL | 514,959,887 | 67,969 | 51,996,248 |

#### Species occurrences

Sequences revealed 2,313 hymenopteran detections from 78 taxa. These included 29 species of bee (24 cavity-nesting, 5 cleptoparasitic; Supplementary Table 3) and 49 taxa of wasp (27 predatory, 22 parasitic/parasitoid). Species richness and abundance of bees and wasps were significantly higher in 2020 than in 2019 (ANOVAs: bee abundance p=0.01, bee richness p<0.01, wasp abundance p<0.01, wasp richness p<0.01). In 2020, 25 species of bee and 41 species of wasp were found, compared to 7 and 16 in 2019. There was considerable overlap between years, with 85% of species found in 2019 also found in 2020. A further break-down of bee and wasp diversity as well as species richness variation between nests can be seen in Supplementary Figures 1 and 2.

In addition to hymenopteran occupants, 328 detections from 11 other insect orders were found. Twelve taxa of parasitic/parasitoid insects were detected, including 10 Diptera and one each of Coleoptera and Strepsiptera. As well, 72 taxa of potential prey were detected, including 10 Araneae, 10 Coleoptera, five Diptera, one Ephemeroptera, eight Hemiptera, 27 Lepidoptera, seven Orthoptera, and four Thysanoptera species. Three additional taxa (*Forficula auricularia* (Dermaptera), *Trogoderma inclusum* (Coleoptera), *Elipsocus moebiusi* (Psocoptera)) are thought to be incidental detections. Literature does not classify them as parasites, predators, pollinators, or prey, and all are opportunistic in determining their nest. For one taxon (*Megaselia* sp. (Diptera)) we were unable to determine whether we obtained a parasitic species, a scavenger or prey. All options were reported from other *Megaselia* species including a known bee parasitoid (*M. rufipes*). The sequences obtained did not match any of 379 species on BOLD which could point to a yet unknown species with unknown life history.

Analysis of plant tissue revealed 384 plant genera from 91 families and 36 orders (Figure 2). A full list of all identified genera can be found in the Supplementary Table 4.

**Figure 2:**
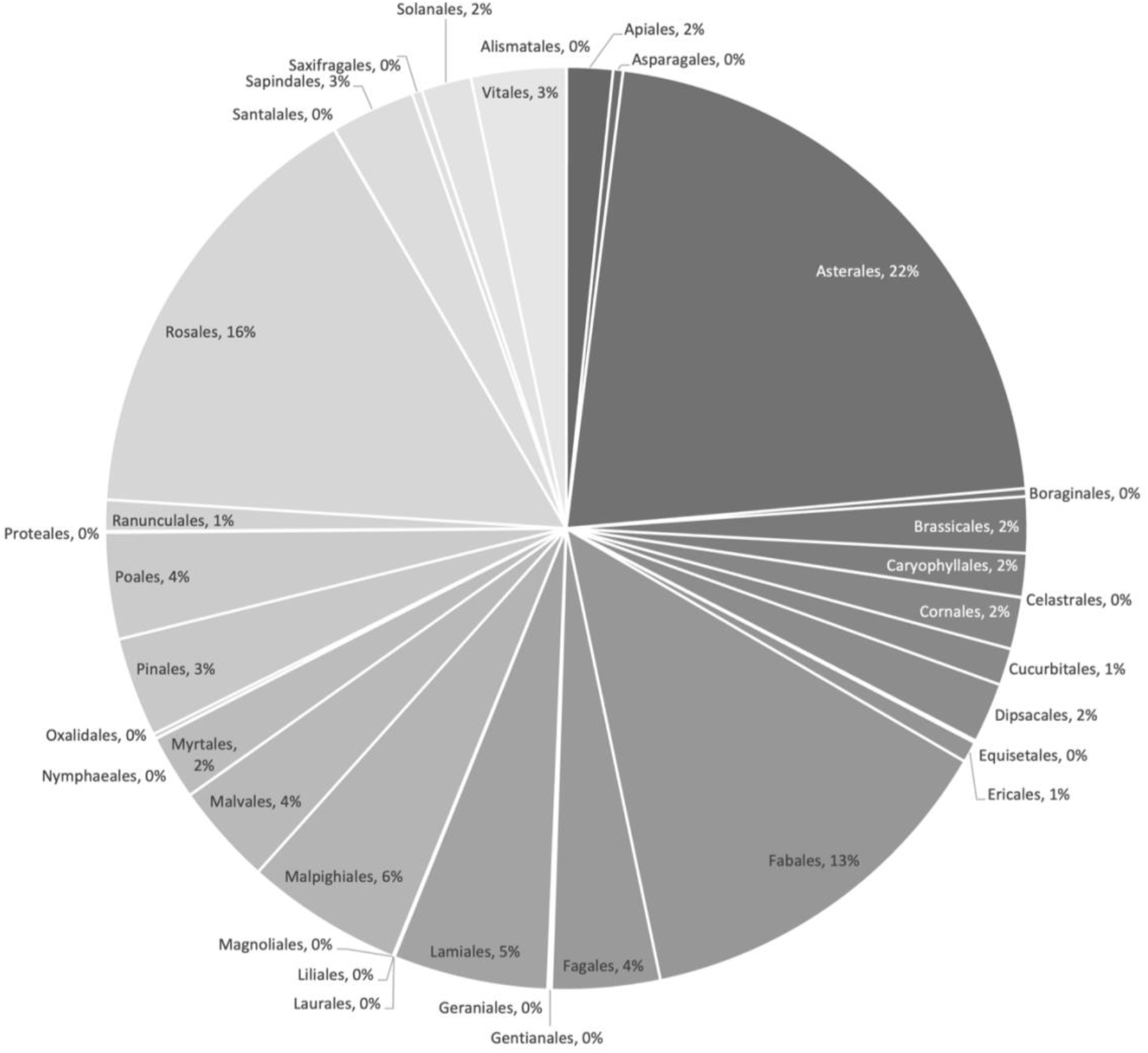
Percentage of all plant order pollen detections in cardboard tubes in trap nests in 2019 and 2020 found through rbcL metabarcoding.

A food network including all natural enemies, predatory wasps, pollinators, plants, and prey demonstrates the entire complexity of species interactions detected (Figure 3). Various categories of interactions are described in the the sections below.

**Figure 3:**
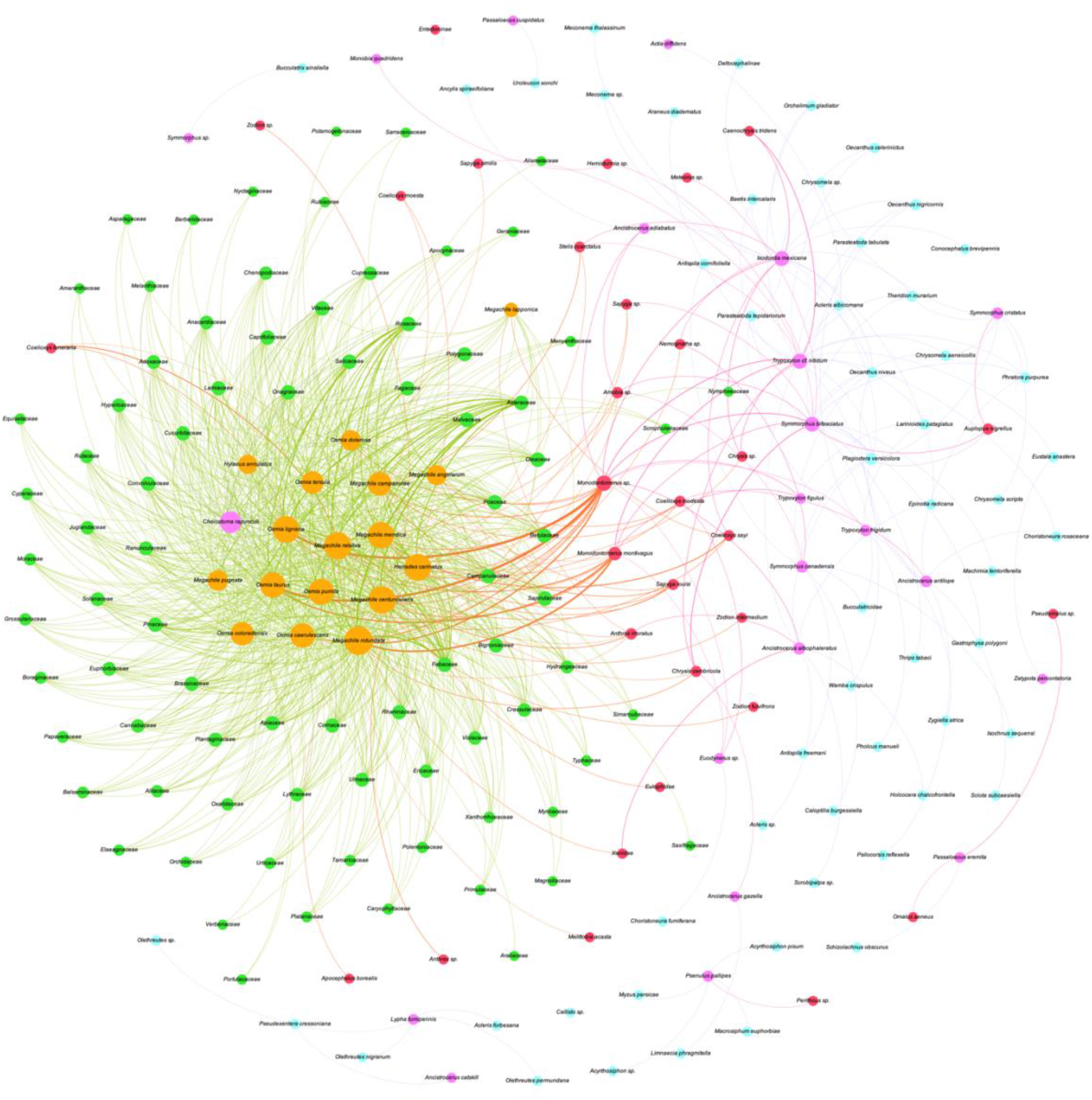
Tripartite network showing interactions between parasites (red), predatory wasps (pink), pollinators (orange), insect prey (blue), and plants (green). The diameter of the circles represents species abundance, and the thickness of a connecting line shows interaction strength.

### Cavity-nesting bees and wasps

#### Interspecific nest sharing

Nest occupancy was investigated using a read count threshold of 1,000 to distinguish true visits from accidental visits by insects. Overall, 45% of tubes (284) were found with multiple (two to four) Hymenoptera occupants. Often these were hymenopteran parasites/parasitoids along with their hosts, but once those instances were removed, 155 cases remained where multiple predators and/or pollinators nested in the same tube (Table 3, Supplementary Table 5).

**Table 3:** Nest sharing occupant combinations in nest boxes in 2019 and 2020.

| Occupant combination | Number of tubes |
| --- | --- |
| Two pollinators | 59 |
| Two predators | 18 |
| One pollinator/one predator | 43 |
| Two pollinators/one predator | 13 |
| Three pollinators | 7 |
| Three predators | 2 |
| Two pollinators/two predators | 2 |
| Three pollinators/one predators | 1 |
| Four pollinators | 1 |

#### Nesting materials

A variety of nesting material was recorded from occupied nest tubes, classified as: cellophane, grass, leaves, masticated leaves, mud, rocks/sand, straw, tree sap, or none (Supplementary Table 6). Nesting material was identified for 746 predatory wasps and pollinating bees. Mud was the most common nesting material (36%), followed by leaves (24%), and tree sap (14%). Nineteen nests had mixed nesting materials and 10 of these nests were occupied by two (or more) species.

#### Cavity-nesting bees

A total of 22 species of cavity-nesting bees were included in a bipartite analysis with 362 plant genera from 88 plant families. All bipartite networks created based on site, landcover designation, or bee species showed patterns consistent with polylectic pollen consumption: i.e. each bee species harvested pollen from a variety of plant genera and families. The number of unique plant families and genera visited by each species at crop, forest and urban sites is shown in Supplementary Table 7. Polylectic foraging agrees with historical foraging records (Supplementary Table 8) for all but three bee species (*Chelostoma rapunculi, Megachile lapponica, Megachile pugnata*) that were considered oligolectic (Gathmann and Tscharntke 2002, Sheffield et al. 2011). Three out of four *C. rapunculi*, two out of four *M. lapponica*, and one out of seven *M. pugnata* were detected in nests with their known host plant, in addition to many other plant genera.

Although 88 plant families were detected and used in bipartite networks (Figure 4), most interactions with bees were from three families: Asteraceae (20%), Fabaceae (13%), and Rosaceae (13%). Nestedness of the overall bipartite plant-pollinator network was low (19.7) at the plant family level, showing that interactions at sites with low species richness were not generally a subset of interactions at sites with high species richness. At the plant family level, the overall network showed an average of 8.7 links per bee species and a connectance of 0.59. The urban network demonstrated the highest number of pollinators (18) and plants (77), links per species (8.3), connectance (0.57), nestedness (19), and cluster coefficient (0.6) of the landscape-based networks. Full details of all plant-pollinator network metrics can be found in Supplementary Table 9.

**Figure 4:**
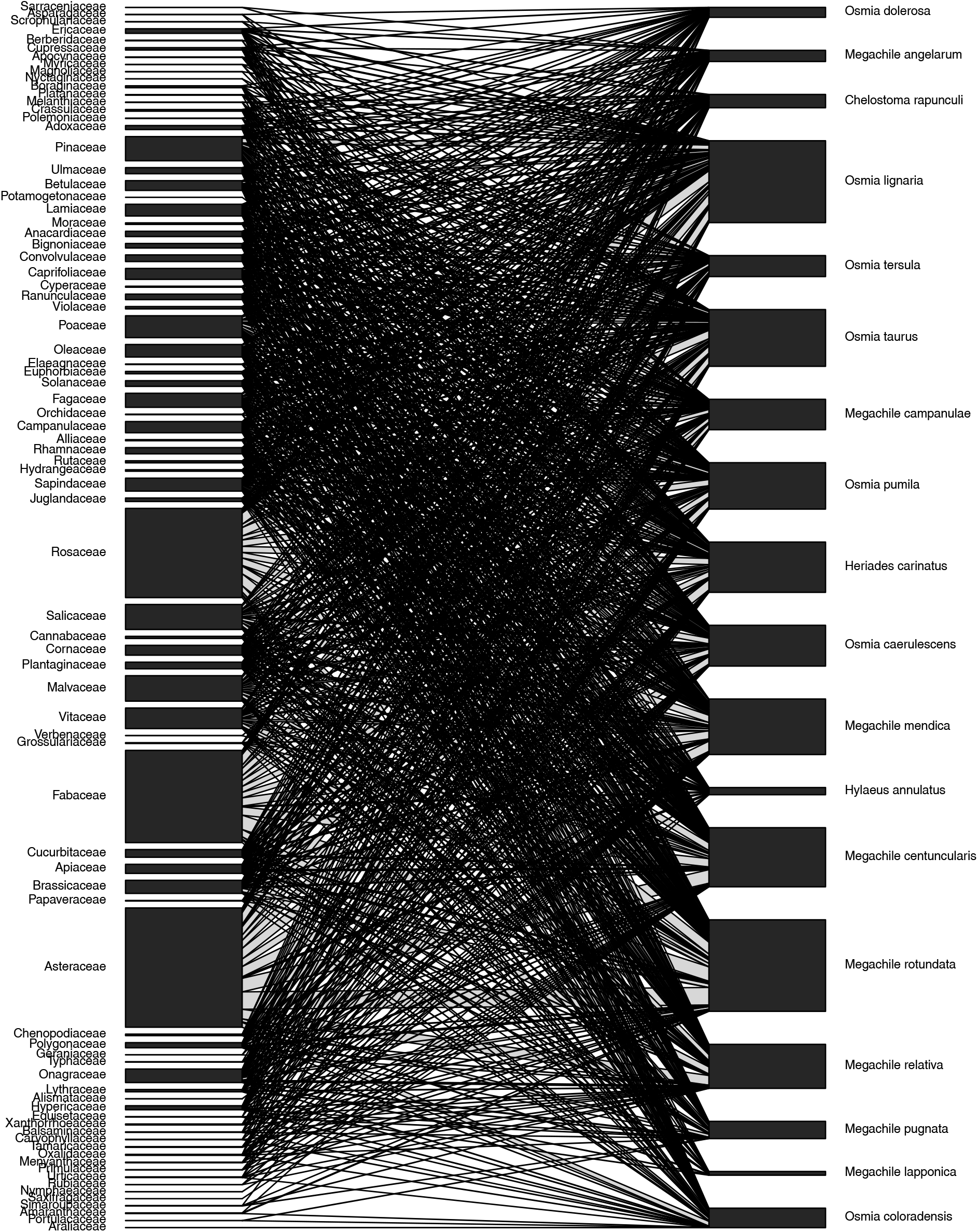
Bipartite network of all pollen-collecting bees found with pollen/nectar in trap nests in 2019 and 2020.

When compared to previous distributional records (accessed from GBIF and discoverlife.org), 10 bee species appear to have expanded ranges, including three non-native species (*Chelostoma rapunculi, Hylaeus pictipes, Osmia taurus)* and seven native species (*Coelioxys modesta, Megachile campanulae, Megachile angelarum, Megachile snowi, Megachile mendica, Osmia dolerosa, Osmia caerulescens*). Distribution maps for these species are shown in Supplementary Figure 2, while maps of the 17 species with ranges comparable to existing records can be found in Supplementary Figure 3.

#### Cavity-nesting wasps

Distributional ranges of wasps were mapped and compared to previous distributional records (GBIF, discoverlife.org). Four species were found to have potentially expanded ranges (*Passaloecus eremita, Passaloecus gracilis*, *Psenulus pallipes, Trypoxylon* cf. *nitidum –* Supplementary Figure 4). Predatory wasps were analyzed in bipartite networks to evaluate their interactions with prey (Figure 5). Eleven of these species were identified in cavities with prey that is not prominent in historical records. All predatory interactions can be seen in Supplementary Table 10.

**Figure 5:**
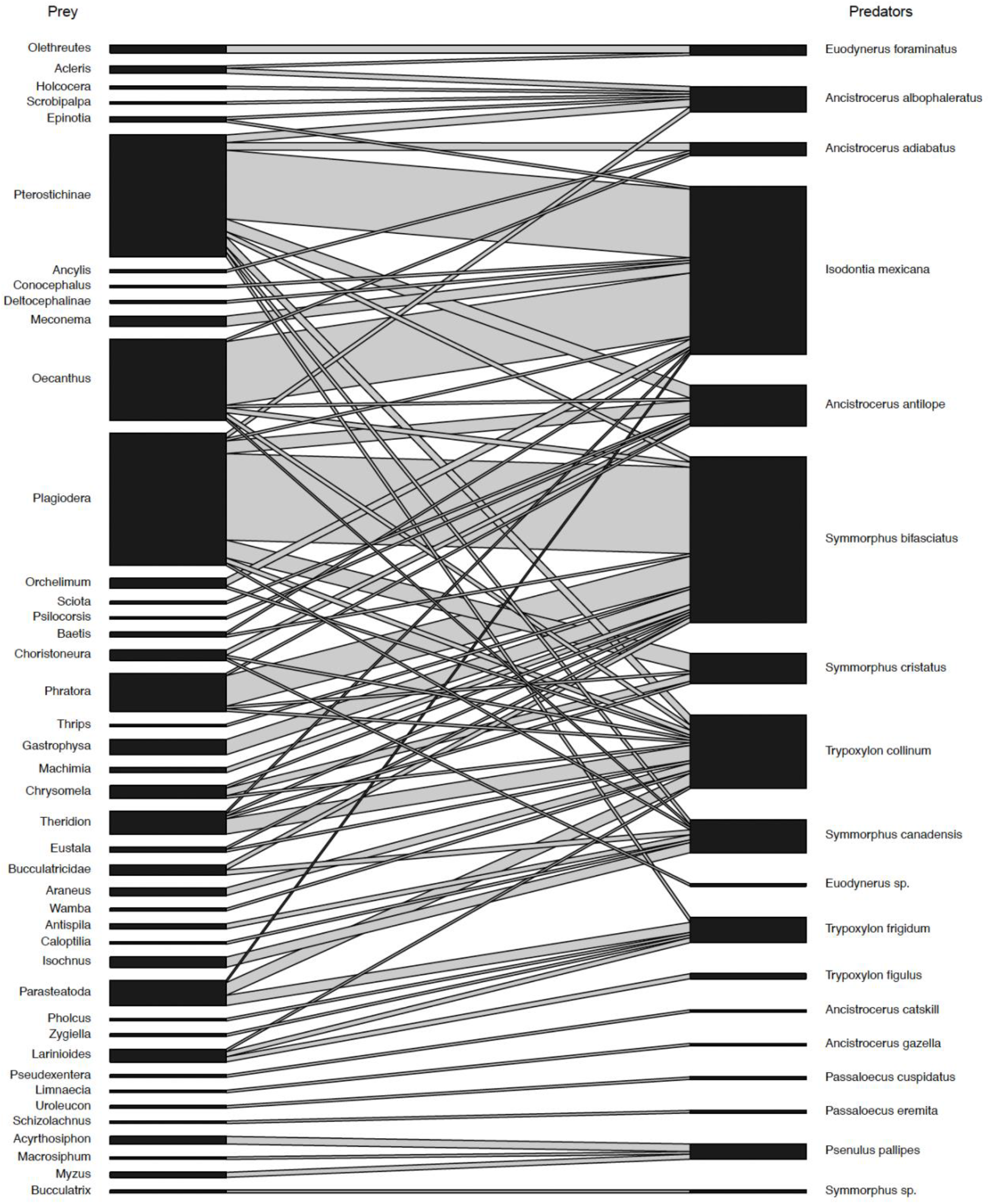
Bipartite network depicting prey and predators in trap nests in 2019 and 2020.

#### Natural enemies

Trap nests from 59 sites (49%) and 152 cardboard tubes (27%) were found to contain natural enemies (parasites/parasitoids) and their host(s) when investigated with read counts greater than 1,000. The percentage of tubes parasitized per site ranged from nine to 100%. Of all tubes, 30 had multiple natural enemy detections. Parasitic and parasitoid wasps, flies, and beetles were analyzed in bipartite networks to evaluate their interactions with hosts (Figure 6). Ten natural enemies were identified in cavities with hosts not prominent in historical records. All parasitic interactions are shown in Supplementary Table 11.

**Figure 6:**
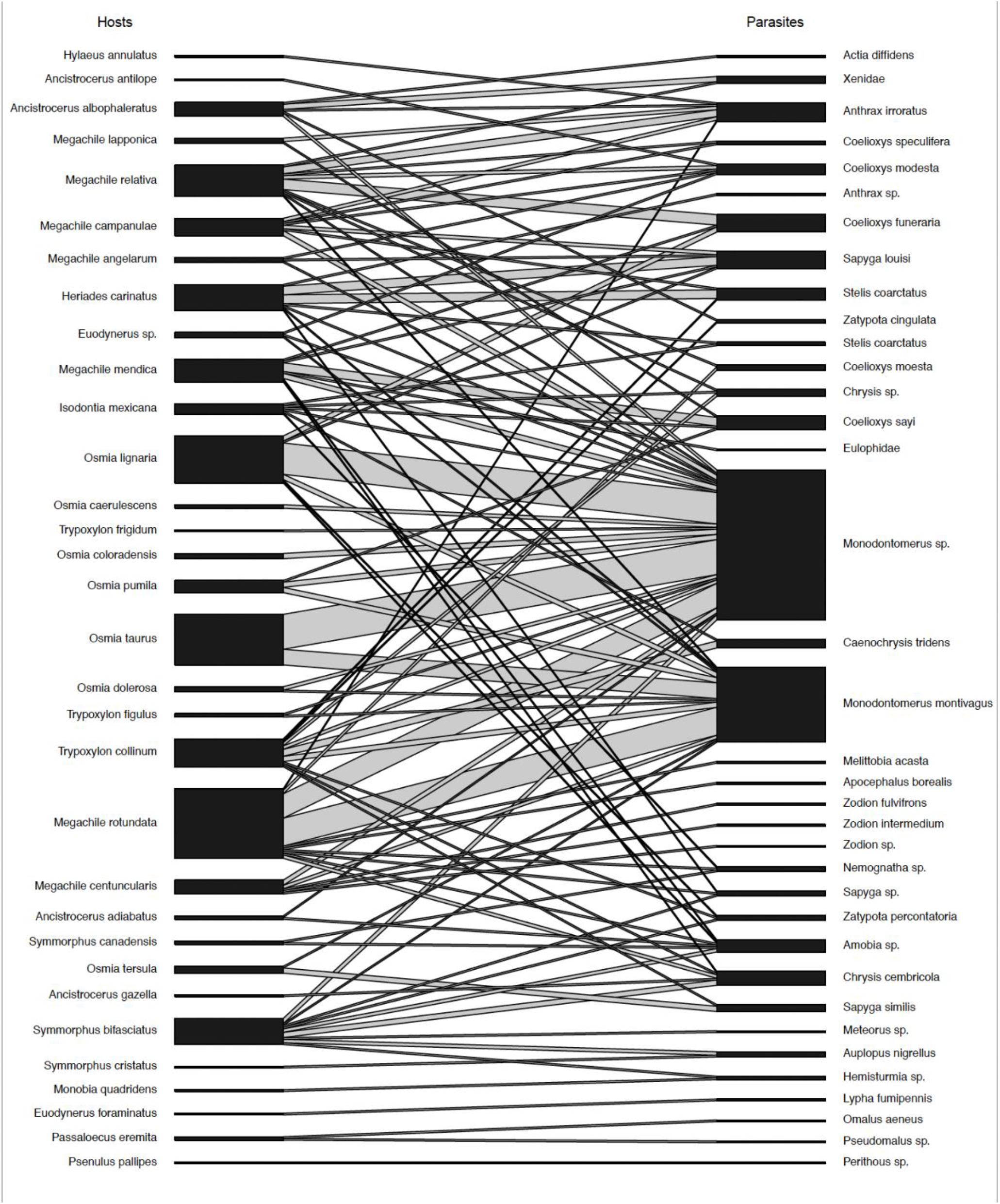
Bipartite network depicting parasites and hosts in trap nests in 2019 and 2020.

#### Landcover

Landcover was examined around each site using QGIS v3.18.3 (QGIS Development Team 2021) and the 2015 Land Cover of Canada dataset (Natural Resources Canada 2015). Sites were assigned a landcover designation based on which landcover type (crop, forest, or urban) covered the largest percentage of land within a 500m radius of a site (Table 4). Comparisons of bee and wasp species richness and abundance among landcover types (Table 4) did not reveal any significant patterns (p>0.05) except for wasp abundance in 2019 (ANOVA, p = 0.04). However, post-hoc tests showed no significant differences (Tukey’s Honest Significant Difference p>0.05).

**Table 4:**
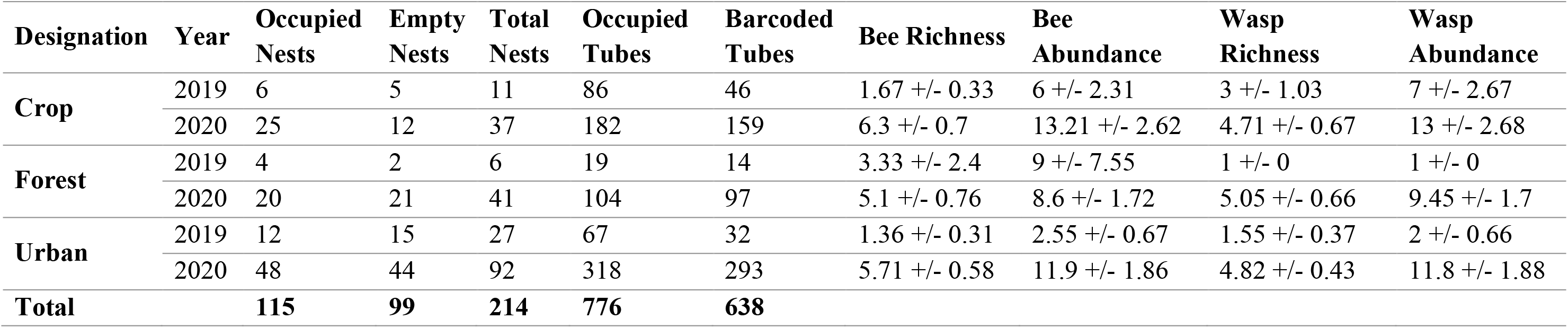
Mean trap-nest occupancy, number of occupied and barcoded tubes, bee and wasp species richness and abundances (+/- SE) during 2019 and 2020, sorted by landcover designation. Landcover designation was determined in QGIS using the 2015 Land Cover of Canada dataset within a 500-metre radius circle around each site.

While there were no differences in abundance or species richness, bee community composition differed among landcover types with *Megachile rotundata* most abundant in cropping landscapes, *Megachile relativa* most abundant in forested landscapes, and *Osmia lignaria* and *Megachile rotundata* most common in urban areas.

## Discussion

Solitary cavity-nesting bees and wasps, and their natural enemies, provide important ecosystem services through their interactions (Ollerton et al. 2011, Hoffmann et al. 2018, Brock et al. 2021). This study further investigated solitary cavity-nesting bee and wasp distributions and interactions across Canada and evaluated how local landcover affected their nesting. DNA barcoding revealed complex interactions previously not detected through observation.

Overall, 83% of occupied tubes were successfully barcoded representing 113 sites, however when evaluated by year, 52% of 2019 tubes were successful compared with 91% of 2020 tubes. The main difference in sample processing between years was that 2019 samples were sequenced on an Illumina MiSeq while 2020 samples were sequenced on Illumina’s NovaSeq platform. It is likely that the increased read depth of the latter resulted in this increased success. In addition, lab protocols were optimized for the second year.

### Cavity-nesting bees and wasps

#### Interspecific nest sharing

A quarter of all nesting tubes contained two or more solitary cavity-nesting predatory wasp and/or pollinating bee species (Supplementary Table 5). This pattern might occur, even in the presence of sufficient empty nests, if the first species abandons their nest early and another takes up occupancy or if the nest is usurped (Černá et al. 2013). Usurpation occurs when a female takes over the nest of another female that is in the process of being built (Černá et al. 2013). Multiple occupancies have been reported in other studies including interspecific usurpation in *Megachile inermis* and *Megachile relativa* nests where each species was both usurped and usurper (Strickler et al. 1996). Over six years of sampling, usurpation rates ranged from 0-20% (Strickler et al. 1996). From their trap-nesting study, O’Neill and O’Neill (2018) found 3% of nests in which the first species of bee or wasp was superseded by another. They report instances of either two bees or wasps or a combination of the two co-habiting. Similarly, Mesquita and Augusto (2011) found 4.6% and 13.6% of trap nests contained two species at two sampling sites. It is possible that the high proportion of multiple occupancies found in this study was due to increased detection sensitivity through DNA barcoding compared to previous studies in which occupants were allowed to hatch out into contained spaces. In those cases, any pupae which failed to hatch would have been very difficult to identify. Additionally, hosts or prey would have been consumed, confounding determination of who the parasite or predator was. For these reasons, metabarcoding represents a superior approach to investigating interspecific nest sharing.

Although undetectable with our methodology, intraspecific nest usurpation or multiple occupancy in solitary bees and wasps has been studied more frequently than interspecific usurpation (Field 1992, McCorquodale and Owen 1994, Kim 1997, Vinson and Frankie 2000, Bosch and Vicens 2006, Černá et al. 2013). Černá et al. (2013) found that 10-45% of nests changed ownership during the active season, though the percentage of those that were true usurpations is lower. Nest usurpation can be a positive ecological strategy for multiple reasons. Since nest building is a high-cost activity, usurping or using a partially constructed nest can save energy and shorten the amount of time it takes to build a nest (Field 1992, Vinson and Frankie 2000). As well, laying eggs in multiple nests spreads the risk of parasitism and increases the probability of successful nests (Field 1992). However, if a nest is usurped by a later season species, the original founder will have their larvae killed as they will be blocked inside. Therefore, founding bees will build thick caps in their entrances to prevent usurpation, sometimes even doing so multiple times between foraging trips.

Our results together with prior studies provide rationale for the large number of nests found with two species; however, they do not address the cases (N=35) where three or four species were found together. Whilst it is possible that these represent multiple abandonments or usurpations, this remains an interesting area for future studies to investigate.

#### Nesting materials

In many cases, bees and wasp species were detected in tubes whose nesting materials did not match their well-known preferences (Cane et al. 2007). For example, leaf-cutting bees were found in tubes with soil as the nesting material and mason bees were found in tubes with leaf disks. It is possible that some bees were detected as only visitors to a nest, not occupants, because DNA barcoding is a very sensitive detection technique. Solitary bees depend heavily on olfactory, visual, and spatial cues to locate their nests (Guédot et al. 2006), and if something affects their ability to identify their nest (e.g., pesticide exposure (Artz and Pitts-Singer 2015)), they might enter another one incorrectly. However, the read count threshold was raised to a level where we assumed detections represent true occupants (minimum 1,000 reads). Some of these mismatched nesting materials might be due to tubes being used by multiple occupants. However, this does not explain instances of mismatched nesting materials in tubes with single occupancy. Once we accounted for nests which shared by two or more species, there were still many instances of singly occupying species using nesting materials that are not considered typical. For example, *Heriades carinatus* is known to use tree resin as a nesting material and we found 19 instances confirming this. However, we also found *Heriades carinatus* in tubes lined with mud, grass, masticated leaves, and cellophane even when it was nesting alone. Further cases similar to this can be seen for an additional 27 species in Supplementary Table 6.

Typically, solitary bees have adaptations to assist with collection of specific types of nesting materials (Cane et al. 2007). For example, leaf-collecting bees have sharp mandibles for cutting while species which masticate their leaves have mandibles better adapted to chewing (Cane et al. 2007). This generally leaves bees poorly suited to collect nesting material outside of the norm. However, some species are found to collect multiple nesting substrates. For example, *Megachile pugnata* has been observed to use a combination of leaves, masticated leaves, and soil to form their nests (Medler 1964). In our study, species observed more than 10 times tended to have a primary nesting substrate (Supplementary Table 6). Although some bee species were quite rigid in their nest material choices, this indicates the possibility that some cavity-nesting bee species are fluid in their choice of nesting substrate if their preferred choice is unavailable.

#### Cavity-nesting bee foraging

Asteraceae (20%), Fabaceae (13%), and Rosaceae (13%) were the most abundant plant families found in tube nests occupied by cavity nesting bees. Asteraceae and Fabaceae are two of the largest plant families (Panero and Crozier 2016), so their high level of detection is unsurprising. Plant-pollinator bipartite networks revealed pollen consumption patterns consistent with generalist foragers. For the majority of bee species detected, this agrees with historical records classifying them as polylectic. However, three species classified as oligolectic appeared to forage on plant genera and families outside their known hosts.

*Chelostoma rapunculi* is oligolectic on *Campanula*, though Buck et al. (2005) noted that their specimens were collected on non-*Campanula* flowers. Three out of four of our *C. rapunculi* detections were found with *Campanula* pollen in addition to 107 other genera of plants. However, three out of four of these were also found in nests occupied by multiple species. Similarly, while *Megachile lapponica* is reported to be oligolectic on *Epilobium* (Gathmann and Tscharntke 2002), we recorded four *M. lapponica* with 37 genera of pollen not including *Epilobium*. Two of the four *M. lapponica* were in multiply occupied nests. *Megachile pugnata* is known as a sunflower (*Helianthus*) specialist (Sheffield et al. 2011), however of seven detections at separate sites, only one was found to provision her offspring with *Helianthus* pollen in addition to pollen from 122 other plant genera. Five out of seven *M. pugnata* were in multiple occupied nests. These nests likely explain some of the additional plant genera detected, however they cannot explain the missing known specialist pollen sources. Comparing plant-pollinator networks made through observation with those made using pollen metabarcoding, some species were considered oligolectic when networks were made through observation, but their classification changed to polylectic when pollen metabarcoding data was used instead (Arstingstall et al. 2021). Some of these additional observations might be explained by instances when a bee inadvertently collected pollen from multiple plants when visiting one flower, if that pollen landed there through the actions of wind or other flower visitors, but that pollen could still represent a valuable food source. As plant-centred visual surveys tend to skew the true number of specialist bee species (Bosch et al. 2009), it is therefore possible that some of the bee species generally considered oligolectic will use alternative pollen sources when needed. However, since specialists are defined by the pollen they collect, it could also be that detections of nectar within bee bread are contributing to increased plant diversity of foraging sources. Determination of the exact reason for these interactions is outside the scope of this study, but it is likely to be a combination of multiple nesting, nectar detection, and sensitive detection through DNA barcoding.

The overall plant-pollinator network showed a community with low nestedness. This indicates high beta diversity and dissimilar species assemblages among sites (Wright and Reeves 1992, Dormann et al. 2009). Although low nestedness is often associated with low network stability (Gresty et al. 2018), in this case it is understandable that interactions among sites differ from one another since this network represents sites Canada-wide. Overall network connectance was 0.59 indicates that 59% of the potential interactions were realized (Dormann et al. 2009, Rivera-Hutinel et al. 2012). This connectance was driven by having many species in the network and many single links, a number that is likely even higher. A higher level of connectance suggest high network stability since there is some redundancy (Gresty et al. 2018). Many studies of plant-pollinator networks collected their data through observation as opposed to molecular methods. Because of this, our study shows a far higher proportion of plants to pollinators that is reflected in the negative web asymmetry.

#### Cavity-nesting bee ranges

Installation of trap-nests across Canada paired with DNA barcoding detected bee species outside their known ranges. Knowledge of species’ ranges is essential to make informed management decisions and track changes over time (Jamieson et al. 2019). Although our sampling was not site- intensive, three non-native and seven native bee species were found in previously unrecorded provinces or areas. Whether these detections represent true range expansions or are the result of previous sampling in these areas is difficult to determine. While some may be due to low sampling, native western Canada species like *Megachile angelarum*, *Megachile snowi*, and *Osmia dolerosa* were all found in southern Ontario that is intensively sampled by multiple working groups. Specifically, MacIvor (2019) sampled cavity-nesting bees in Toronto for three seasons and did not report detecting any of these three species. This sampling occurred from 2011 to 2013 so it is possible these species have shifted to Ontario since then. Likewise, *Coelioxys modesta* and *Megachile campanulae* are native eastern Canada species that were detected in southern British Columbia. Sheffield and Heron (2019) provide a comprehensive checklist of bee species in British Columbia but do not include either *C. modesta* or *M. campanulae*. The recency of this species list publication makes it unlikely that these species shifted that quickly, instead it might be that they were difficult to detect using methods other than trap-nesting. This is often true of kleptoparasite species that do not collect pollen for their offspring, such as *Coelioxys*, which are not as attracted to pan traps as pollen-foraging species (Wilson et al. 2008). While native species *Megachile mendica* and *Osmia caerulescens* were found within provinces where they had previously been recorded, they were found further north than earlier records indicate. It is therefore likely that these represent range expansions northward, although how recently these ranges expanded is difficult to say without more detailed survey information.

We found *Chelostoma rapunculi*, native to the Palearctic range but previously only reported in Ontario, at one site in British Columbia. *Hylaeus pictipes* is native to Europe but was previously only reported from Ontario, and we also found it in Manitoba. Finally, and most surprisingly, we found numerous detections of *Osmia taurus* in British Columbia, Alberta, Manitoba, Ontario, and Prince Edward Island while it has previously only been seen in Ontario. The frequency and wide distribution of *O. taurus* records is particularly interesting because its recent progress throughout the United States and into Canada has been closely watched, with a rapid increase of *O. taurus* observations in the Mid-Atlantic United States since its first detection in 2002 (Le Croy et al. 2020). This species is thought to have been accidentally imported in place of *Osmia cornifrons* that is used agriculturally and looks very similar (Gibbs et al. 2017, LeCroy et al. 2020). This finding is perhaps the most contentious because detecting it so far north represents a massive range expansion that has not been detected previously. Reference specimens which were 100% barcode matches were sent to a taxonomic expert (Dr. Tom Onuferko, Canadian Museum of Nature), for revision to confirm this finding. The identity as *Osmia taurus* was indeed confirmed, meaning they have expanded far further than previously thought.

Solitary cavity-nesting bees are uniquely positioned to be distributed to new areas quickly due to their nesting habits. While they might nest in shipping containers and be accidentally transported, there are also companies who intentionally ship cavity-nesting bee cocoons. For example, Crown Bees (Woodinville, WA) ships mason and leaf-cutting bees to Canada and the United States. They market both *Osmia lignaria* and *Osmia cornifrons*, although their method of buying back mason bee cocoons from customers in the fall means they cannot be sure that the species that are overwintering are what they advertise. There are dozens of similar companies which might also contribute to the rapid spread of non-native species or parasites.

#### Cavity-nesting wasp prey

Solitary wasps in the families Pompilidae, Sphecidae, and Vespidae are considered prey specialists, while the Crabronidae are more generalists (Brock et al. 2021). Of 23 species of cavity-nesting wasp that were analyzed in bipartite networks, 10 were found with prey outside their recorded choices. As a member of the genus *Trypoxylon*, *Trypoxylon* cf. *nitidum* is historically known as a spider-collecting wasp. In addition to spiders in the genera Araneidae and Theridiidae, *T.* cf. *nitidum* collected beetle and moth larvae as well as grasshoppers. *Isodontia mexicana* is likewise well-known to collect grasshoppers but was also noted with beetle and moth larvae and spiders. Multiple species of *Ancistrocerus* were identified with beetle larvae or grasshopper prey, in addition to their recorded preference for moth larvae. Some of these detections are likely explained by nest-sharing, however they might also represent a less preferred but still acceptable prey. Many of these historical predator-prey records are from observational studies from the 1900s, so further investigation using modern methods should be conducted.

#### Cavity-nesting wasp ranges

As with cavity-nesting bees, tracking the ranges of cavity-nesting wasps is important for informing management decisions (Jamieson et al. 2019). We found four species outside their historically reported ranges in Canada. Three of these are non-native species that had previously been found in Ontario or eastern Canada, but we additionally found them in British Columbia. These species had few detections meaning they might have only recently arrived on the west coast. *Trypoxylon nitidum* is native to South America (Scher and das Graças Pompolo 2003), though a few records are available from southern Ontario. In addition to Ontario, we found *Trypoxylon* cf. *nitidum* in British Columbia, Alberta, and Manitoba. At this point, further investigation into this is required since it might represent a closely related but undocumented species or a significant range expansion.

#### Natural enemies

Half of all occupied sites and a quarter of occupied nesting tubes were parasitized. Previous studies report parasitism rates of 13% of nests (Loyola and Martins 2006) or 5% (Klein et al. 2006) and 7% (Steckel et al. 2014) of brood cells. Once again, those studies relied on hatching out occupants to observe parasitism that could contribute to the higher rates seen in this study. It has been suggested that trap nests could increase parasitism rates (Moenen 2012, MacIvor and Packer 2015), attributing this mainly to the high nesting densities that provide excellent host sources for parasites (Harris et al. 2021). Trap nests encourage different species to nest in proximity through differently sized tubes, which may also allow parasites to cross between species.

In addition to the high parasitism rates, ten natural enemies were recorded with hosts outside previous records. Finding recent host records in Canada was difficult, if not impossible, for many of these natural enemies that may explain some of the unexpected parasite-host interactions.

*Caenochrysis tridens* parasitized *Trypoxylon* cf. *nitidum*, although its host is reported to be *Trypoxylon politum* in Pennsylvania (Conrow et al. 2016). In this case, *T. politum* is not found as north or west as *C. tridens* and so it used a similar but alternative host that might be common but not yet recorded. *Symmorphus canadensis* has been reported as a host for *Chrysis cembricola* (Krombein and Hurd 1979) and since then no further hosts have been named. We found *C. cembricola* to parasitize five species of bees and wasps, including *Symmorphus bifasciatus*. Four *Coelioxys* species were found with unrecorded hosts. As *Coelioxys* are known as parasites of *Megachile* bees (Scott et al. 2000), new *Megachile* sp. records are not surprising. *Coelioxys* were also found parasitizing three species of wasps. *Stelis coarctatus* is known to parasitize *Heriades carinatus*, though Gibbs et al. (2017) notes that it has a broad host range, meaning the additional bee and wasp hosts we found help to expand that list. *Sapyga louisi* and *Sapyga similis* were both found parasitizing new hosts, and since recent host checklists could not be found this study starts to fill that gap. *Amobia* sp., a dipteran parasite, was found parasitizing Megachilidae for the first time. Previously they have been noted to parasitize Vepid and Sphecid wasps (Spofford et al. 1989), and *Hylaeus* to be *Amobia* hosts (Marinho and Vivallo 2020).

#### Landcover

There are many reports of anthropogenic modification of landscapes affecting species and their interactions, including urbanisation (Hooper et al. 2005, Chace and Walsh 2006, McKinney 2006, Winfree et al. 2009). Urban landscapes have been observed to support reduced pollinator biodiversity (Elmqvist et al. 2003, Potts et al. 2010, Sánchez-Bayo and Wyckhuys 2019), with impacts largely based on the degree of urbanization and the type of bee (Wenzel et al. 2020). Some studies have found that urban areas positively influence solitary bee populations and areas of medium urbanization generally support pollinator populations, especially when compared to intensive agricultural areas (Fortel et al. 2014). Compared to ground-nesting bees, cavity-nesting bees tend to fare better in the face of urbanization since they do not depend on exposed soil for nesting (Wenzel et al. 2020). With these previous findings in mind, the lack of significant results when comparing bee and wasp species richness and abundance between cropping, forest, and urban landcovers is not surprising. Sampling sites were all located either on school property or on the teachers’ property during COVID-19 lockdown. While these sites were distinguishable as being predominantly agricultural, forest, or urban, it is unlikely that many of them would be considered intensely agricultural or intensely urban landcovers. It is thought that this may have reduced the signal of abundance or species richness differences among landcover types.

It is likely that the significant differences in wasp species richness and abundance in 2019 forests is due to lower sample sizes. Likewise, the significantly higher species richness and abundance found in 2020 compared to 2019 is likely due to increased sampling in 2020. These trends can be examined further once additional data are collected from further years of sampling.

#### Implications, considerations, and future direction

This study shows the immense amount of information that can be gathered using community science and DNA metabarcoding. Community science allowed this study to gather samples from a far larger area than could have been accomplished otherwise. Additionally, this provided an incredible educational opportunity, and Bees@Schools reached hundreds of students, teachers, and families. Multiple teachers reported that participating in the Bees@Schools program encouraged them to begin a native pollinator garden initiative with the whole school. The benefits to using community science are many, as outlined by MacPhail and Colla (2020). Although challenges such as study design, sample consistency, and time management presented some difficulties, the benefits were greater. Pairing the augmented data collection power of community science with DNA barcoding can result in large and detailed datasets (Henter et al. 2016, Steinke et al. 2017). The study of interactions proves challenging, and metabarcoding is one method of increasing the number of observable interactions (Pornon et al. 2017). Thanks to the combination of community science and DNA metabarcoding, a number of novel interactions and distributions were discovered.

It is important to note the limitations of this study. The quality of reference libraries is essential for accurate identifications. Both the BOLD and GenBank reference libraries are public data repositories which depend on correct taxonomic information (Meiklejohn et al. 2019, Pentinsaari et al. 2020). For barcoding to work, errors must be avoided in both sequence preparation and sequence analysis. Extreme care was taken during sequence preparation and analysis, and any potentially erroneous identifications were investigated further. Due to the destructive nature of the DNA metabarcoding protocol used, there are no reference specimens. The protocol was carefully designed, and raw data were cautiously processed and examined to greatly reduce the chance for error. All species found with potential new ranges were compared to BOLD to ensure the existence of a reference specimen in a collection. In the cases of *Osmia taurus* and *Trypoxylon* cf. *nitidum*, these 100% barcode match reference specimens were sent to or photographed for taxonomic experts to corroborate their identities, and at this time, *Osmia taurus* has been confirmed. Although the use of metabarcoding brings additional considerations, the benefits and quality of data it produced are excellent. Although extreme care was taken to separate leafy nesting material from larval samples, it may be that some nesting materials were detected. In the future, barcoding nesting materials separately could produce a dataset to compare pollen records to. However, that was beyond the scope of this study.

Ensuring a large read depth during sequencing increased the number of successfully identified occupants and reduced cost per sample, which is essential for the continued success of a community science program. Data collection through the Bees@Schools program continued in 2021, 2022 and 2023, so further analyses will be done with a larger dataset.

These results greatly expand our knowledge of parasite-host and predator-prey interactions in the solitary cavity-nesting Hymenoptera. Barcoding trap nest inhabitants has provided great insight into interactions not previously observed by the great studies of the 20^th^ century (Krombein 1967, Krombein and Hurd 1979). Natural history observations for many of these wasp species have not been pursued using updated technology, and great benefits can continue to be had from doing so. While bees are more extensively studied, there is little recent research on interspecific nest sharing, and new insights were learned about nesting material choices and pollen foraging. Solitary cavity-nesting bees and wasps provide essential pollination and predation services. These benefits are fueled by their interactions that are notoriously difficult to study. Further knowledge of these interactions can help advise conservation policies meant to support these beneficial insects and the ecosystem services they provide.

## Acknowledgements

The authors are extremely grateful to all the Canadian students and teachers for their participation in the Bees@School project. We also thank the members of the Raine and Steinke labs for their support. Financial support for this work was provided by the Food from Thought: Agricultural Systems for a Healthy Planet Initiative, through the Canada First Research Excellence Fund (Project 000054), Natural Sciences and Engineering Research Council of Canada (NSERC) Discovery Grant (2015-06783), Ontario Ministry of Agriculture, Food and Rural Affairs (UofG2018-3307). NER is supported as the Rebanks Family Chair in Pollinator Conservation by the Weston Family Foundation.

