## Supplementary data for "Welcome to Hotel Hymenoptera: monitoring cavity-nesting bee and wasp distribution and their trophic interactions using community science and metabarcoding"

*In order of occurrence in manuscript main text*

**Supplementary Table 1: Primers used for first round PCR**

| Primer name | Primer sequence (5‘ - 3‘) | Primer conc. in PCR (μM) | Target gene | Fragment length (bp) | Reference |
|---|---|---|---|---|---|
| **BF3** | CCHGAYATRGCHTTYCCHCG | 0.5 | COI | 418 | Elbrecht et al. 2019 |
| **BR2** | TCDGGRTGNCCRAARAAYCA | 0.5 | | | |
| **rbcL1** | TTGGCAGCATTYCGAGTAACTCC | 0.5 | rbcL | 184 | Palmieri et al. 2009; Little 2014 |
| **rbcLB** | AACCYTCTTCAAAAAGGTC | 0.5 | | | |

**Supplementary Table 2: COI and rbcL fusion primers for PCR2**

| COI ID | Illumina Adaptor | Tag | Primer |
|---|---|---|---|
| P5-BF3_a | AATGATACGGCGACCACCGAGATCTACACTCTTTCCCTACACGACGCTCTTCCGATCT | TACCAC | CCHGAYATRGCHTTYCCHCG |
| P5-BF3_b | AATGATACGGCGACCACCGAGATCTACACTCTTTCCCTACACGACGCTCTTCCGATCT | ACCATAA | CCHGAYATRGCHTTYCCHCG |
| P5-BF3_c | AATGATACGGCGACCACCGAGATCTACACTCTTTCCCTACACGACGCTCTTCCGATCT | CTGTATCG | CCHGAYATRGCHTTYCCHCG |
| P5-BF3_d | AATGATACGGCGACCACCGAGATCTACACTCTTTCCCTACACGACGCTCTTCCGATCT | TTATGCAAG | CCHGAYATRGCHTTYCCHCG |
| P5-BF3_e | AATGATACGGCGACCACCGAGATCTACACTCTTTCCCTACACGACGCTCTTCCGATCT | CACCATGGAA | CCHGAYATRGCHTTYCCHCG |
| P5-BF3_f | AATGATACGGCGACCACCGAGATCTACACTCTTTCCCTACACGACGCTCTTCCGATCT | GGTAAC | CCHGAYATRGCHTTYCCHCG |
| P5-BF3_g | AATGATACGGCGACCACCGAGATCTACACTCTTTCCCTACACGACGCTCTTCCGATCT | AATTCTA | CCHGAYATRGCHTTYCCHCG |
| P5-BF3_h | AATGATACGGCGACCACCGAGATCTACACTCTTTCCCTACACGACGCTCTTCCGATCT | CTCGGAGA | CCHGAYATRGCHTTYCCHCG |
| P5-BF3_i | AATGATACGGCGACCACCGAGATCTACACTCTTTCCCTACACGACGCTCTTCCGATCT | TAGTTGGGT | CCHGAYATRGCHTTYCCHCG |
| P5-BF3_j | AATGATACGGCGACCACCGAGATCTACACTCTTTCCCTACACGACGCTCTTCCGATCT | GGTGAGCAGA | CCHGAYATRGCHTTYCCHCG |
| P5-BF3_k | AATGATACGGCGACCACCGAGATCTACACTCTTTCCCTACACGACGCTCTTCCGATCT | GAAATG | CCHGAYATRGCHTTYCCHCG |
| P5-BF3_l | AATGATACGGCGACCACCGAGATCTACACTCTTTCCCTACACGACGCTCTTCCGATCT | TGGACCA | CCHGAYATRGCHTTYCCHCG |
| P5-BF3_m | AATGATACGGCGACCACCGAGATCTACACTCTTTCCCTACACGACGCTCTTCCGATCT | GCACAGAG | CCHGAYATRGCHTTYCCHCG |
| P5-BF3_n | AATGATACGGCGACCACCGAGATCTACACTCTTTCCCTACACGACGCTCTTCCGATCT | ATCGGCTGA | CCHGAYATRGCHTTYCCHCG |
| P5-BF3_o | AATGATACGGCGACCACCGAGATCTACACTCTTTCCCTACACGACGCTCTTCCGATCT | TGTAGAGAAA | CCHGAYATRGCHTTYCCHCG |
| P5-BF3_p | AATGATACGGCGACCACCGAGATCTACACTCTTTCCCTACACGACGCTCTTCCGATCT | CCCTTG | CCHGAYATRGCHTTYCCHCG |
| P5-BF3_q | AATGATACGGCGACCACCGAGATCTACACTCTTTCCCTACACGACGCTCTTCCGATCT | ACAGCTG | CCHGAYATRGCHTTYCCHCG |
| P5-BF3_r | AATGATACGGCGACCACCGAGATCTACACTCTTTCCCTACACGACGCTCTTCCGATCT | AACGTGGG | CCHGAYATRGCHTTYCCHCG |
| P5-BF3_s | AATGATACGGCGACCACCGAGATCTACACTCTTTCCCTACACGACGCTCTTCCGATCT | CCGCATATT | CCHGAYATRGCHTTYCCHCG |
| P5-BF3_t | AATGATACGGCGACCACCGAGATCTACACTCTTTCCCTACACGACGCTCTTCCGATCT | CGTTGCCAAT | CCHGAYATRGCHTTYCCHCG |
| P5-BF3_u | AATGATACGGCGACCACCGAGATCTACACTCTTTCCCTACACGACGCTCTTCCGATCT | ATCTCC | CCHGAYATRGCHTTYCCHCG |
| P5-BF3_v | AATGATACGGCGACCACCGAGATCTACACTCTTTCCCTACACGACGCTCTTCCGATCT | CGTCGTA | CCHGAYATRGCHTTYCCHCG |
| P5-BF3_w | AATGATACGGCGACCACCGAGATCTACACTCTTTCCCTACACGACGCTCTTCCGATCT | CATAACAT | CCHGAYATRGCHTTYCCHCG |
| P5-BF3_x | AATGATACGGCGACCACCGAGATCTACACTCTTTCCCTACACGACGCTCTTCCGATCT | CAGTAATTA | CCHGAYATRGCHTTYCCHCG |
| P5-BR2_a | AATGATACGGCGACCACCGAGATCTACACTCTTTCCCTACACGACGCTCTTCCGATCT | GTATCCCGGT | TCDGGRTGNCCRAARAAYCA |
| P5-BR2_b | AATGATACGGCGACCACCGAGATCTACACTCTTTCCCTACACGACGCTCTTCCGATCT | TGAGTT | TCDGGRTGNCCRAARAAYCA |
| P5-BR2_c | AATGATACGGCGACCACCGAGATCTACACTCTTTCCCTACACGACGCTCTTCCGATCT | CTGATGT | TCDGGRTGNCCRAARAAYCA |
| P5-BR2_d | AATGATACGGCGACCACCGAGATCTACACTCTTTCCCTACACGACGCTCTTCCGATCT | GCACGTAC | TCDGGRTGNCCRAARAAYCA |
| P5-BR2_e | AATGATACGGCGACCACCGAGATCTACACTCTTTCCCTACACGACGCTCTTCCGATCT | TTAGATCGG | TCDGGRTGNCCRAARAAYCA |
| P5-BR2_f | AATGATACGGCGACCACCGAGATCTACACTCTTTCCCTACACGACGCTCTTCCGATCT | GAAACGAAGA | TCDGGRTGNCCRAARAAYCA |
| P5-BR2_g | AATGATACGGCGACCACCGAGATCTACACTCTTTCCCTACACGACGCTCTTCCGATCT | ACCGGA | TCDGGRTGNCCRAARAAYCA |
| P5-BR2_h | AATGATACGGCGACCACCGAGATCTACACTCTTTCCCTACACGACGCTCTTCCGATCT | ACACCCT | TCDGGRTGNCCRAARAAYCA |
| P5-BR2_i | AATGATACGGCGACCACCGAGATCTACACTCTTTCCCTACACGACGCTCTTCCGATCT | TCGCTTTA | TCDGGRTGNCCRAARAAYCA |
| P5-BR2_j | AATGATACGGCGACCACCGAGATCTACACTCTTTCCCTACACGACGCTCTTCCGATCT | TCTATACGA | TCDGGRTGNCCRAARAAYCA |
| P5-BR2_k | AATGATACGGCGACCACCGAGATCTACACTCTTTCCCTACACGACGCTCTTCCGATCT | TTCTCAGGGT | TCDGGRTGNCCRAARAAYCA |
| P5-BR2_l | AATGATACGGCGACCACCGAGATCTACACTCTTTCCCTACACGACGCTCTTCCGATCT | AGAACA | TCDGGRTGNCCRAARAAYCA |

| COI ID | Illumina Adaptor | Tag | Primer |
|---|---|---|---|
| P5-BR2_m | AATGATACGGCGACCACCGAGATCTACACTCTTTCCCTACACGACGCTCTTCCGATCT | ATTCAAT | TCDGGRTGNCCRAARAAYCA |
| P5-BR2_n | AATGATACGGCGACCACCGAGATCTACACTCTTTCCCTACACGACGCTCTTCCGATCT | TCTTTCTT | TCDGGRTGNCCRAARAAYCA |
| P5-BR2_o | AATGATACGGCGACCACCGAGATCTACACTCTTTCCCTACACGACGCTCTTCCGATCT | TAGGAGGAA | TCDGGRTGNCCRAARAAYCA |
| P5-BR2_p | AATGATACGGCGACCACCGAGATCTACACTCTTTCCCTACACGACGCTCTTCCGATCT | GTGCCAGAAT | TCDGGRTGNCCRAARAAYCA |
| P5-BR2_q | AATGATACGGCGACCACCGAGATCTACACTCTTTCCCTACACGACGCTCTTCCGATCT | GAGGCC | TCDGGRTGNCCRAARAAYCA |
| P5-BR2_r | AATGATACGGCGACCACCGAGATCTACACTCTTTCCCTACACGACGCTCTTCCGATCT | AAGACTC | TCDGGRTGNCCRAARAAYCA |
| P5-BR2_s | AATGATACGGCGACCACCGAGATCTACACTCTTTCCCTACACGACGCTCTTCCGATCT | CGCCGGAA | TCDGGRTGNCCRAARAAYCA |
| P5-BR2_t | AATGATACGGCGACCACCGAGATCTACACTCTTTCCCTACACGACGCTCTTCCGATCT | CGTTTTGGA | TCDGGRTGNCCRAARAAYCA |
| P5-BR2_u | AATGATACGGCGACCACCGAGATCTACACTCTTTCCCTACACGACGCTCTTCCGATCT | GTCATCGGGA | TCDGGRTGNCCRAARAAYCA |
| P5-BR2_v | AATGATACGGCGACCACCGAGATCTACACTCTTTCCCTACACGACGCTCTTCCGATCT | ACTTAG | TCDGGRTGNCCRAARAAYCA |
| P5-BR2_w | AATGATACGGCGACCACCGAGATCTACACTCTTTCCCTACACGACGCTCTTCCGATCT | GGGGAAA | TCDGGRTGNCCRAARAAYCA |
| P5-BR2_x | AATGATACGGCGACCACCGAGATCTACACTCTTTCCCTACACGACGCTCTTCCGATCT | GGAGGAGT | TCDGGRTGNCCRAARAAYCA |
| P7-BF3_a | CAAGCAGAAGACGGCATACGAGATGTGACTGGAGTTCAGACGTGTGCTCTTCCGATCT | TACCAC | CCHGAYATRGCHTTYCCHCG |
| P7-BF3_b | CAAGCAGAAGACGGCATACGAGATGTGACTGGAGTTCAGACGTGTGCTCTTCCGATCT | ACCATAA | CCHGAYATRGCHTTYCCHCG |
| P7-BF3_c | CAAGCAGAAGACGGCATACGAGATGTGACTGGAGTTCAGACGTGTGCTCTTCCGATCT | CTGTATCG | CCHGAYATRGCHTTYCCHCG |
| P7-BF3_d | CAAGCAGAAGACGGCATACGAGATGTGACTGGAGTTCAGACGTGTGCTCTTCCGATCT | TTATGCAAG | CCHGAYATRGCHTTYCCHCG |
| P7-BF3_e | CAAGCAGAAGACGGCATACGAGATGTGACTGGAGTTCAGACGTGTGCTCTTCCGATCT | CACCATGGAA | CCHGAYATRGCHTTYCCHCG |
| P7-BF3_f | CAAGCAGAAGACGGCATACGAGATGTGACTGGAGTTCAGACGTGTGCTCTTCCGATCT | GGTAAC | CCHGAYATRGCHTTYCCHCG |
| P7-BF3_g | CAAGCAGAAGACGGCATACGAGATGTGACTGGAGTTCAGACGTGTGCTCTTCCGATCT | AATTCTA | CCHGAYATRGCHTTYCCHCG |
| P7-BF3_h | CAAGCAGAAGACGGCATACGAGATGTGACTGGAGTTCAGACGTGTGCTCTTCCGATCT | CTCGGAGA | CCHGAYATRGCHTTYCCHCG |
| P7-BF3_i | CAAGCAGAAGACGGCATACGAGATGTGACTGGAGTTCAGACGTGTGCTCTTCCGATCT | TAGTTGGGT | CCHGAYATRGCHTTYCCHCG |
| P7-BF3_j | CAAGCAGAAGACGGCATACGAGATGTGACTGGAGTTCAGACGTGTGCTCTTCCGATCT | GGTGAGCAGA | CCHGAYATRGCHTTYCCHCG |
| P7-BF3_k | CAAGCAGAAGACGGCATACGAGATGTGACTGGAGTTCAGACGTGTGCTCTTCCGATCT | GAAATG | CCHGAYATRGCHTTYCCHCG |
| P7-BF3_l | CAAGCAGAAGACGGCATACGAGATGTGACTGGAGTTCAGACGTGTGCTCTTCCGATCT | TGGACCA | CCHGAYATRGCHTTYCCHCG |
| P7-BF3_m | CAAGCAGAAGACGGCATACGAGATGTGACTGGAGTTCAGACGTGTGCTCTTCCGATCT | GCACAGAG | CCHGAYATRGCHTTYCCHCG |
| P7-BF3_n | CAAGCAGAAGACGGCATACGAGATGTGACTGGAGTTCAGACGTGTGCTCTTCCGATCT | ATCGGCTGA | CCHGAYATRGCHTTYCCHCG |
| P7-BF3_o | CAAGCAGAAGACGGCATACGAGATGTGACTGGAGTTCAGACGTGTGCTCTTCCGATCT | TGTAGAGAAA | CCHGAYATRGCHTTYCCHCG |
| P7-BF3_p | CAAGCAGAAGACGGCATACGAGATGTGACTGGAGTTCAGACGTGTGCTCTTCCGATCT | CCCTTG | CCHGAYATRGCHTTYCCHCG |
| P7-BF3_q | CAAGCAGAAGACGGCATACGAGATGTGACTGGAGTTCAGACGTGTGCTCTTCCGATCT | ACAGCTG | CCHGAYATRGCHTTYCCHCG |
| P7-BF3_r | CAAGCAGAAGACGGCATACGAGATGTGACTGGAGTTCAGACGTGTGCTCTTCCGATCT | AACGTGGG | CCHGAYATRGCHTTYCCHCG |
| P7-BF3_s | CAAGCAGAAGACGGCATACGAGATGTGACTGGAGTTCAGACGTGTGCTCTTCCGATCT | CCGCATATT | CCHGAYATRGCHTTYCCHCG |
| P7-BF3_t | CAAGCAGAAGACGGCATACGAGATGTGACTGGAGTTCAGACGTGTGCTCTTCCGATCT | CGTTGCCAAT | CCHGAYATRGCHTTYCCHCG |
| P7-BF3_u | CAAGCAGAAGACGGCATACGAGATGTGACTGGAGTTCAGACGTGTGCTCTTCCGATCT | ATCTCC | CCHGAYATRGCHTTYCCHCG |
| P7-BF3_v | CAAGCAGAAGACGGCATACGAGATGTGACTGGAGTTCAGACGTGTGCTCTTCCGATCT | CGTCGTA | CCHGAYATRGCHTTYCCHCG |
| P7-BF3_w | CAAGCAGAAGACGGCATACGAGATGTGACTGGAGTTCAGACGTGTGCTCTTCCGATCT | CATAACAT | CCHGAYATRGCHTTYCCHCG |
| P7-BF3_x | CAAGCAGAAGACGGCATACGAGATGTGACTGGAGTTCAGACGTGTGCTCTTCCGATCT | CAGTAATTA | CCHGAYATRGCHTTYCCHCG |
| P7-BR2_a | CAAGCAGAAGACGGCATACGAGATGTGACTGGAGTTCAGACGTGTGCTCTTCCGATCT | GTATCCCGGT | TCDGGRTGNCCRAARAAYCA |
| P7-BR2_b | CAAGCAGAAGACGGCATACGAGATGTGACTGGAGTTCAGACGTGTGCTCTTCCGATCT | TGAGTT | TCDGGRTGNCCRAARAAYCA |
| P7-BR2_c | CAAGCAGAAGACGGCATACGAGATGTGACTGGAGTTCAGACGTGTGCTCTTCCGATCT | CTGATGT | TCDGGRTGNCCRAARAAYCA |

| COI ID | Illumina Adaptor | Tag | Primer |
|---|---|---|---|
| P7-BR2_d | CAAGCAGAAGACGGCATACGAGATGTGACTGGAGTTCAGACGTGTGCTCTTCCGATCT | GCACGTAC | TCDGGRTGNCCRAARAAYCA |
| P7-BR2_e | CAAGCAGAAGACGGCATACGAGATGTGACTGGAGTTCAGACGTGTGCTCTTCCGATCT | TTAGATCGG | TCDGGRTGNCCRAARAAYCA |
| P7-BR2_f | CAAGCAGAAGACGGCATACGAGATGTGACTGGAGTTCAGACGTGTGCTCTTCCGATCT | GAAACGAAGA | TCDGGRTGNCCRAARAAYCA |
| P7-BR2_g | CAAGCAGAAGACGGCATACGAGATGTGACTGGAGTTCAGACGTGTGCTCTTCCGATCT | ACCGGA | TCDGGRTGNCCRAARAAYCA |
| P7-BR2_h | CAAGCAGAAGACGGCATACGAGATGTGACTGGAGTTCAGACGTGTGCTCTTCCGATCT | ACACCCT | TCDGGRTGNCCRAARAAYCA |
| P7-BR2_i | CAAGCAGAAGACGGCATACGAGATGTGACTGGAGTTCAGACGTGTGCTCTTCCGATCT | TCGCTTTA | TCDGGRTGNCCRAARAAYCA |
| P7-BR2_j | CAAGCAGAAGACGGCATACGAGATGTGACTGGAGTTCAGACGTGTGCTCTTCCGATCT | TCTATACGA | TCDGGRTGNCCRAARAAYCA |
| P7-BR2_k | CAAGCAGAAGACGGCATACGAGATGTGACTGGAGTTCAGACGTGTGCTCTTCCGATCT | TTCTCAGGGT | TCDGGRTGNCCRAARAAYCA |
| P7-BR2_l | CAAGCAGAAGACGGCATACGAGATGTGACTGGAGTTCAGACGTGTGCTCTTCCGATCT | AGAACA | TCDGGRTGNCCRAARAAYCA |
| P7-BR2_m | CAAGCAGAAGACGGCATACGAGATGTGACTGGAGTTCAGACGTGTGCTCTTCCGATCT | ATTCAAT | TCDGGRTGNCCRAARAAYCA |
| P7-BR2_n | CAAGCAGAAGACGGCATACGAGATGTGACTGGAGTTCAGACGTGTGCTCTTCCGATCT | TCTTTCTT | TCDGGRTGNCCRAARAAYCA |
| P7-BR2_o | CAAGCAGAAGACGGCATACGAGATGTGACTGGAGTTCAGACGTGTGCTCTTCCGATCT | TAGGAGGAA | TCDGGRTGNCCRAARAAYCA |
| P7-BR2_p | CAAGCAGAAGACGGCATACGAGATGTGACTGGAGTTCAGACGTGTGCTCTTCCGATCT | GTGCCAGAAT | TCDGGRTGNCCRAARAAYCA |
| P7-BR2_q | CAAGCAGAAGACGGCATACGAGATGTGACTGGAGTTCAGACGTGTGCTCTTCCGATCT | GAGGCC | TCDGGRTGNCCRAARAAYCA |
| P7-BR2_r | CAAGCAGAAGACGGCATACGAGATGTGACTGGAGTTCAGACGTGTGCTCTTCCGATCT | AAGACTC | TCDGGRTGNCCRAARAAYCA |
| P7-BR2_s | CAAGCAGAAGACGGCATACGAGATGTGACTGGAGTTCAGACGTGTGCTCTTCCGATCT | CGCCGGAA | TCDGGRTGNCCRAARAAYCA |
| P7-BR2_t | CAAGCAGAAGACGGCATACGAGATGTGACTGGAGTTCAGACGTGTGCTCTTCCGATCT | CGTTTTGGA | TCDGGRTGNCCRAARAAYCA |
| P7-BR2_u | CAAGCAGAAGACGGCATACGAGATGTGACTGGAGTTCAGACGTGTGCTCTTCCGATCT | GTCATCGGGA | TCDGGRTGNCCRAARAAYCA |
| P7-BR2_v | CAAGCAGAAGACGGCATACGAGATGTGACTGGAGTTCAGACGTGTGCTCTTCCGATCT | ACTTAG | TCDGGRTGNCCRAARAAYCA |
| P7-BR2_w | CAAGCAGAAGACGGCATACGAGATGTGACTGGAGTTCAGACGTGTGCTCTTCCGATCT | GGGGAAA | TCDGGRTGNCCRAARAAYCA |
| P7-BR2_x | CAAGCAGAAGACGGCATACGAGATGTGACTGGAGTTCAGACGTGTGCTCTTCCGATCT | GGAGGAGT | TCDGGRTGNCCRAARAAYCA |

| rbcL ID | Illumina Adaptor | Tag | Primer |
|---|---|---|---|
| P5-rbcL1_a | AATGATACGGCGACCACCGAGATCTACACTCTTTCCCTACACGACGCTCTTCCGATCT | CCC | TTGGCAGCATTYCGAGTAACTCC |
| P5-rbcL1_b | AATGATACGGCGACCACCGAGATCTACACTCTTTCCCTACACGACGCTCTTCCGATCT | ACTT | TTGGCAGCATTYCGAGTAACTCC |
| P5-rbcL1_c | AATGATACGGCGACCACCGAGATCTACACTCTTTCCCTACACGACGCTCTTCCGATCT | CTAAC | TTGGCAGCATTYCGAGTAACTCC |
| P5-rbcL1_d | AATGATACGGCGACCACCGAGATCTACACTCTTTCCCTACACGACGCTCTTCCGATCT | GATAGG | TTGGCAGCATTYCGAGTAACTCC |
| P5-rbcL1_e | AATGATACGGCGACCACCGAGATCTACACTCTTTCCCTACACGACGCTCTTCCGATCT | TGTACGC | TTGGCAGCATTYCGAGTAACTCC |
| P5-rbcL1_f | AATGATACGGCGACCACCGAGATCTACACTCTTTCCCTACACGACGCTCTTCCGATCT | AAG | TTGGCAGCATTYCGAGTAACTCC |
| P5-rbcL1_g | AATGATACGGCGACCACCGAGATCTACACTCTTTCCCTACACGACGCTCTTCCGATCT | TACC | TTGGCAGCATTYCGAGTAACTCC |
| P5-rbcL1_h | AATGATACGGCGACCACCGAGATCTACACTCTTTCCCTACACGACGCTCTTCCGATCT | ACGTA | TTGGCAGCATTYCGAGTAACTCC |
| P5-rbcL1_i | AATGATACGGCGACCACCGAGATCTACACTCTTTCCCTACACGACGCTCTTCCGATCT | GGTTGC | TTGGCAGCATTYCGAGTAACTCC |
| P5-rbcL1_j | AATGATACGGCGACCACCGAGATCTACACTCTTTCCCTACACGACGCTCTTCCGATCT | CCGGTGA | TTGGCAGCATTYCGAGTAACTCC |
| P5-rbcL1_k | AATGATACGGCGACCACCGAGATCTACACTCTTTCCCTACACGACGCTCTTCCGATCT | GTTC | TTGGCAGCATTYCGAGTAACTCC |
| P5-rbcL1_l | AATGATACGGCGACCACCGAGATCTACACTCTTTCCCTACACGACGCTCTTCCGATCT | GTGCC | TTGGCAGCATTYCGAGTAACTCC |
| P5-rbcL1_m | AATGATACGGCGACCACCGAGATCTACACTCTTTCCCTACACGACGCTCTTCCGATCT | CCGGCG | TTGGCAGCATTYCGAGTAACTCC |

| rbcL ID | Illumina Adaptor | Tag | Primer |
|---|---|---|---|
| P5-rbcL1_n | AATGATACGGCGACCACCGAGATCTACACTCTTTCCCTACACGACGCTCTTCCGATCT | TAATGTG | TTGGCAGCATTYCGAGTAACTCC |
| P5-rbcL1_o | AATGATACGGCGACCACCGAGATCTACACTCTTTCCCTACACGACGCTCTTCCGATCT | ACA | TTGGCAGCATTYCGAGTAACTCC |
| P5-rbcL1_p | AATGATACGGCGACCACCGAGATCTACACTCTTTCCCTACACGACGCTCTTCCGATCT | CGGA | TTGGCAGCATTYCGAGTAACTCC |
| P5-rbcL1_q | AATGATACGGCGACCACCGAGATCTACACTCTTTCCCTACACGACGCTCTTCCGATCT | TCAGA | TTGGCAGCATTYCGAGTAACTCC |
| P5-rbcL1_r | AATGATACGGCGACCACCGAGATCTACACTCTTTCCCTACACGACGCTCTTCCGATCT | TGTGTC | TTGGCAGCATTYCGAGTAACTCC |
| P5-rbcL1_s | AATGATACGGCGACCACCGAGATCTACACTCTTTCCCTACACGACGCTCTTCCGATCT | GGCACCG | TTGGCAGCATTYCGAGTAACTCC |
| P5-rbcL1_t | AATGATACGGCGACCACCGAGATCTACACTCTTTCCCTACACGACGCTCTTCCGATCT | CTT | TTGGCAGCATTYCGAGTAACTCC |
| P5-rbcL1_u | AATGATACGGCGACCACCGAGATCTACACTCTTTCCCTACACGACGCTCTTCCGATCT | TATT | TTGGCAGCATTYCGAGTAACTCC |
| P5-rbcL1_v | AATGATACGGCGACCACCGAGATCTACACTCTTTCCCTACACGACGCTCTTCCGATCT | GCGAT | TTGGCAGCATTYCGAGTAACTCC |
| P5-rbcL1_w | AATGATACGGCGACCACCGAGATCTACACTCTTTCCCTACACGACGCTCTTCCGATCT | ACTTAG | TTGGCAGCATTYCGAGTAACTCC |
| P5-rbcL1_x | AATGATACGGCGACCACCGAGATCTACACTCTTTCCCTACACGACGCTCTTCCGATCT | AAGACTC | TTGGCAGCATTYCGAGTAACTCC |
| P5-rbcLB_a | AATGATACGGCGACCACCGAGATCTACACTCTTTCCCTACACGACGCTCTTCCGATCT | GCC | AACCYTCTTCAAAAAGGTC |
| P5-rbcLB_b | AATGATACGGCGACCACCGAGATCTACACTCTTTCCCTACACGACGCTCTTCCGATCT | TTCC | AACCYTCTTCAAAAAGGTC |
| P5-rbcLB_c | AATGATACGGCGACCACCGAGATCTACACTCTTTCCCTACACGACGCTCTTCCGATCT | TAGCG | AACCYTCTTCAAAAAGGTC |
| P5-rbcLB_d | AATGATACGGCGACCACCGAGATCTACACTCTTTCCCTACACGACGCTCTTCCGATCT | TAAGCC | AACCYTCTTCAAAAAGGTC |
| P5-rbcLB_e | AATGATACGGCGACCACCGAGATCTACACTCTTTCCCTACACGACGCTCTTCCGATCT | GTAACCG | AACCYTCTTCAAAAAGGTC |
| P5-rbcLB_f | AATGATACGGCGACCACCGAGATCTACACTCTTTCCCTACACGACGCTCTTCCGATCT | AAA | AACCYTCTTCAAAAAGGTC |
| P5-rbcLB_g | AATGATACGGCGACCACCGAGATCTACACTCTTTCCCTACACGACGCTCTTCCGATCT | ACGC | AACCYTCTTCAAAAAGGTC |
| P5-rbcLB_h | AATGATACGGCGACCACCGAGATCTACACTCTTTCCCTACACGACGCTCTTCCGATCT | CTATG | AACCYTCTTCAAAAAGGTC |
| P5-rbcLB_i | AATGATACGGCGACCACCGAGATCTACACTCTTTCCCTACACGACGCTCTTCCGATCT | CGCGCG | AACCYTCTTCAAAAAGGTC |
| P5-rbcLB_j | AATGATACGGCGACCACCGAGATCTACACTCTTTCCCTACACGACGCTCTTCCGATCT | TGCGGCT | AACCYTCTTCAAAAAGGTC |
| P5-rbcLB_k | AATGATACGGCGACCACCGAGATCTACACTCTTTCCCTACACGACGCTCTTCCGATCT | CGAG | AACCYTCTTCAAAAAGGTC |
| P5-rbcLB_l | AATGATACGGCGACCACCGAGATCTACACTCTTTCCCTACACGACGCTCTTCCGATCT | AGTGG | AACCYTCTTCAAAAAGGTC |
| P5-rbcLB_m | AATGATACGGCGACCACCGAGATCTACACTCTTTCCCTACACGACGCTCTTCCGATCT | AGGCAT | AACCYTCTTCAAAAAGGTC |
| P5-rbcLB_n | AATGATACGGCGACCACCGAGATCTACACTCTTTCCCTACACGACGCTCTTCCGATCT | AGACTCC | AACCYTCTTCAAAAAGGTC |
| P5-rbcLB_o | AATGATACGGCGACCACCGAGATCTACACTCTTTCCCTACACGACGCTCTTCCGATCT | GAGGCC | AACCYTCTTCAAAAAGGTC |
| P5-rbcLB_p | AATGATACGGCGACCACCGAGATCTACACTCTTTCCCTACACGACGCTCTTCCGATCT | CCCT | AACCYTCTTCAAAAAGGTC |
| P5-rbcLB_q | AATGATACGGCGACCACCGAGATCTACACTCTTTCCCTACACGACGCTCTTCCGATCT | ATCAT | AACCYTCTTCAAAAAGGTC |
| P5-rbcLB_r | AATGATACGGCGACCACCGAGATCTACACTCTTTCCCTACACGACGCTCTTCCGATCT | GGTATT | AACCYTCTTCAAAAAGGTC |
| P5-rbcLB_s | AATGATACGGCGACCACCGAGATCTACACTCTTTCCCTACACGACGCTCTTCCGATCT | TTACTGC | AACCYTCTTCAAAAAGGTC |
| P5-rbcLB_t | AATGATACGGCGACCACCGAGATCTACACTCTTTCCCTACACGACGCTCTTCCGATCT | GTTT | AACCYTCTTCAAAAAGGTC |
| P5-rbcLB_u | AATGATACGGCGACCACCGAGATCTACACTCTTTCCCTACACGACGCTCTTCCGATCT | GACTT | AACCYTCTTCAAAAAGGTC |
| P5-rbcLB_v | AATGATACGGCGACCACCGAGATCTACACTCTTTCCCTACACGACGCTCTTCCGATCT | TACCAC | AACCYTCTTCAAAAAGGTC |
| P5-rbcLB_w | AATGATACGGCGACCACCGAGATCTACACTCTTTCCCTACACGACGCTCTTCCGATCT | ACCATAA | AACCYTCTTCAAAAAGGTC |
| P5-rbcLB_x | AATGATACGGCGACCACCGAGATCTACACTCTTTCCCTACACGACGCTCTTCCGATCT | GAA | AACCYTCTTCAAAAAGGTC |
| P7-rbcL1_a | CAAGCAGAAGACGGCATACGAGATGTGACTGGAGTTCAGACGTGTGCTCTTCCGATCT | CCC | TTGGCAGCATTYCGAGTAACTCC |
| P7-rbcL1_b | CAAGCAGAAGACGGCATACGAGATGTGACTGGAGTTCAGACGTGTGCTCTTCCGATCT | ACTT | TTGGCAGCATTYCGAGTAACTCC |
| P7-rbcL1_c | CAAGCAGAAGACGGCATACGAGATGTGACTGGAGTTCAGACGTGTGCTCTTCCGATCT | CTAAC | TTGGCAGCATTYCGAGTAACTCC |
| P7-rbcL1_d | CAAGCAGAAGACGGCATACGAGATGTGACTGGAGTTCAGACGTGTGCTCTTCCGATCT | GATAGG | TTGGCAGCATTYCGAGTAACTCC |

| rbcL ID | Illumina Adaptor | Tag | Primer |
|---|---|---|---|
| P7-rbcL1_e | CAAGCAGAAGACGGCATACGAGATGTGACTGGAGTTCAGACGTGTGCTCTTCCGATCT | TGTACGC | TTGGCAGCATTYCGAGTAACTCC |
| P7-rbcL1_f | CAAGCAGAAGACGGCATACGAGATGTGACTGGAGTTCAGACGTGTGCTCTTCCGATCT | AAG | TTGGCAGCATTYCGAGTAACTCC |
| P7-rbcL1_g | CAAGCAGAAGACGGCATACGAGATGTGACTGGAGTTCAGACGTGTGCTCTTCCGATCT | TACC | TTGGCAGCATTYCGAGTAACTCC |
| P7-rbcL1_h | CAAGCAGAAGACGGCATACGAGATGTGACTGGAGTTCAGACGTGTGCTCTTCCGATCT | ACGTA | TTGGCAGCATTYCGAGTAACTCC |
| P7-rbcL1_i | CAAGCAGAAGACGGCATACGAGATGTGACTGGAGTTCAGACGTGTGCTCTTCCGATCT | GGTTGC | TTGGCAGCATTYCGAGTAACTCC |
| P7-rbcL1_j | CAAGCAGAAGACGGCATACGAGATGTGACTGGAGTTCAGACGTGTGCTCTTCCGATCT | CCGGTGA | TTGGCAGCATTYCGAGTAACTCC |
| P7-rbcL1_k | CAAGCAGAAGACGGCATACGAGATGTGACTGGAGTTCAGACGTGTGCTCTTCCGATCT | GTTC | TTGGCAGCATTYCGAGTAACTCC |
| P7-rbcL1_l | CAAGCAGAAGACGGCATACGAGATGTGACTGGAGTTCAGACGTGTGCTCTTCCGATCT | GTGCC | TTGGCAGCATTYCGAGTAACTCC |
| P7-rbcL1_m | CAAGCAGAAGACGGCATACGAGATGTGACTGGAGTTCAGACGTGTGCTCTTCCGATCT | CCGGCG | TTGGCAGCATTYCGAGTAACTCC |
| P7-rbcL1_n | CAAGCAGAAGACGGCATACGAGATGTGACTGGAGTTCAGACGTGTGCTCTTCCGATCT | TAATGTG | TTGGCAGCATTYCGAGTAACTCC |
| P7-rbcL1_o | CAAGCAGAAGACGGCATACGAGATGTGACTGGAGTTCAGACGTGTGCTCTTCCGATCT | ACA | TTGGCAGCATTYCGAGTAACTCC |
| P7-rbcL1_p | CAAGCAGAAGACGGCATACGAGATGTGACTGGAGTTCAGACGTGTGCTCTTCCGATCT | CGGA | TTGGCAGCATTYCGAGTAACTCC |
| P7-rbcL1_q | CAAGCAGAAGACGGCATACGAGATGTGACTGGAGTTCAGACGTGTGCTCTTCCGATCT | TCAGA | TTGGCAGCATTYCGAGTAACTCC |
| P7-rbcL1_r | CAAGCAGAAGACGGCATACGAGATGTGACTGGAGTTCAGACGTGTGCTCTTCCGATCT | TGTGTC | TTGGCAGCATTYCGAGTAACTCC |
| P7-rbcL1_s | CAAGCAGAAGACGGCATACGAGATGTGACTGGAGTTCAGACGTGTGCTCTTCCGATCT | GGCACCG | TTGGCAGCATTYCGAGTAACTCC |
| P7-rbcL1_t | CAAGCAGAAGACGGCATACGAGATGTGACTGGAGTTCAGACGTGTGCTCTTCCGATCT | CTT | TTGGCAGCATTYCGAGTAACTCC |
| P7-rbcL1_u | CAAGCAGAAGACGGCATACGAGATGTGACTGGAGTTCAGACGTGTGCTCTTCCGATCT | TATT | TTGGCAGCATTYCGAGTAACTCC |
| P7-rbcL1_v | CAAGCAGAAGACGGCATACGAGATGTGACTGGAGTTCAGACGTGTGCTCTTCCGATCT | GCGAT | TTGGCAGCATTYCGAGTAACTCC |
| P7-rbcL1_w | CAAGCAGAAGACGGCATACGAGATGTGACTGGAGTTCAGACGTGTGCTCTTCCGATCT | ACTTAG | TTGGCAGCATTYCGAGTAACTCC |
| P7-rbcL1_x | CAAGCAGAAGACGGCATACGAGATGTGACTGGAGTTCAGACGTGTGCTCTTCCGATCT | AAGACTC | TTGGCAGCATTYCGAGTAACTCC |
| P7-rbcLB_a | CAAGCAGAAGACGGCATACGAGATGTGACTGGAGTTCAGACGTGTGCTCTTCCGATCT | GCC | AACCYTCTTCAAAAAGGTC |
| P7-rbcLB_b | CAAGCAGAAGACGGCATACGAGATGTGACTGGAGTTCAGACGTGTGCTCTTCCGATCT | TTCC | AACCYTCTTCAAAAAGGTC |
| P7-rbcLB_c | CAAGCAGAAGACGGCATACGAGATGTGACTGGAGTTCAGACGTGTGCTCTTCCGATCT | TAGCG | AACCYTCTTCAAAAAGGTC |
| P7-rbcLB_d | CAAGCAGAAGACGGCATACGAGATGTGACTGGAGTTCAGACGTGTGCTCTTCCGATCT | TAAGCC | AACCYTCTTCAAAAAGGTC |
| P7-rbcLB_e | CAAGCAGAAGACGGCATACGAGATGTGACTGGAGTTCAGACGTGTGCTCTTCCGATCT | GTAACCG | AACCYTCTTCAAAAAGGTC |
| P7-rbcLB_f | CAAGCAGAAGACGGCATACGAGATGTGACTGGAGTTCAGACGTGTGCTCTTCCGATCT | AAA | AACCYTCTTCAAAAAGGTC |
| P7-rbcLB_g | CAAGCAGAAGACGGCATACGAGATGTGACTGGAGTTCAGACGTGTGCTCTTCCGATCT | ACGC | AACCYTCTTCAAAAAGGTC |
| P7-rbcLB_h | CAAGCAGAAGACGGCATACGAGATGTGACTGGAGTTCAGACGTGTGCTCTTCCGATCT | CTATG | AACCYTCTTCAAAAAGGTC |
| P7-rbcLB_i | CAAGCAGAAGACGGCATACGAGATGTGACTGGAGTTCAGACGTGTGCTCTTCCGATCT | CGCGCG | AACCYTCTTCAAAAAGGTC |
| P7-rbcLB_j | CAAGCAGAAGACGGCATACGAGATGTGACTGGAGTTCAGACGTGTGCTCTTCCGATCT | TGCGGCT | AACCYTCTTCAAAAAGGTC |
| P7-rbcLB_k | CAAGCAGAAGACGGCATACGAGATGTGACTGGAGTTCAGACGTGTGCTCTTCCGATCT | CGAG | AACCYTCTTCAAAAAGGTC |
| P7-rbcLB_l | CAAGCAGAAGACGGCATACGAGATGTGACTGGAGTTCAGACGTGTGCTCTTCCGATCT | AGTGG | AACCYTCTTCAAAAAGGTC |
| P7-rbcLB_m | CAAGCAGAAGACGGCATACGAGATGTGACTGGAGTTCAGACGTGTGCTCTTCCGATCT | AGGCAT | AACCYTCTTCAAAAAGGTC |
| P7-rbcLB_n | CAAGCAGAAGACGGCATACGAGATGTGACTGGAGTTCAGACGTGTGCTCTTCCGATCT | AGACTCC | AACCYTCTTCAAAAAGGTC |
| P7-rbcLB_o | CAAGCAGAAGACGGCATACGAGATGTGACTGGAGTTCAGACGTGTGCTCTTCCGATCT | GAGGCC | AACCYTCTTCAAAAAGGTC |
| P7-rbcLB_p | CAAGCAGAAGACGGCATACGAGATGTGACTGGAGTTCAGACGTGTGCTCTTCCGATCT | CCCT | AACCYTCTTCAAAAAGGTC |
| P7-rbcLB_q | CAAGCAGAAGACGGCATACGAGATGTGACTGGAGTTCAGACGTGTGCTCTTCCGATCT | ATCAT | AACCYTCTTCAAAAAGGTC |
| P7-rbcLB_r | CAAGCAGAAGACGGCATACGAGATGTGACTGGAGTTCAGACGTGTGCTCTTCCGATCT | GGTATT | AACCYTCTTCAAAAAGGTC |
| P7-rbcLB_s | CAAGCAGAAGACGGCATACGAGATGTGACTGGAGTTCAGACGTGTGCTCTTCCGATCT | TTACTGC | AACCYTCTTCAAAAAGGTC |

| rbcL ID | Illumina Adaptor | Tag | Primer |
|---|---|---|---|
| P7-rbcLB_t | CAAGCAGAAGACGGCATACGAGATGTGACTGGAGTTCAGACGTGTGCTCTTCCGATCT | GTTT | AACCYTCTTCAAAAAGGTC |
| P7-rbcLB_u | CAAGCAGAAGACGGCATACGAGATGTGACTGGAGTTCAGACGTGTGCTCTTCCGATCT | GACTT | AACCYTCTTCAAAAAGGTC |
| P7-rbcLB_v | CAAGCAGAAGACGGCATACGAGATGTGACTGGAGTTCAGACGTGTGCTCTTCCGATCT | TACCAC | AACCYTCTTCAAAAAGGTC |
| P7-rbcLB_w | CAAGCAGAAGACGGCATACGAGATGTGACTGGAGTTCAGACGTGTGCTCTTCCGATCT | ACCATAA | AACCYTCTTCAAAAAGGTC |
| P7-rbcLB_x | CAAGCAGAAGACGGCATACGAGATGTGACTGGAGTTCAGACGTGTGCTCTTCCGATCT | GAA | AACCYTCTTCAAAAAGGTC |
| P7-rbcLB_n | CAAGCAGAAGACGGCATACGAGATGTGACTGGAGTTCAGACGTGTGCTCTTCCGATCT | AGACTCC | AACCYTCTTCAAAAAGGTC |
| P7-rbcLB_o | CAAGCAGAAGACGGCATACGAGATGTGACTGGAGTTCAGACGTGTGCTCTTCCGATCT | GAGGCC | AACCYTCTTCAAAAAGGTC |
| P7-rbcLB_p | CAAGCAGAAGACGGCATACGAGATGTGACTGGAGTTCAGACGTGTGCTCTTCCGATCT | CCCT | AACCYTCTTCAAAAAGGTC |
| P7-rbcLB_q | CAAGCAGAAGACGGCATACGAGATGTGACTGGAGTTCAGACGTGTGCTCTTCCGATCT | ATCAT | AACCYTCTTCAAAAAGGTC |
| P7-rbcLB_r | CAAGCAGAAGACGGCATACGAGATGTGACTGGAGTTCAGACGTGTGCTCTTCCGATCT | GGTATT | AACCYTCTTCAAAAAGGTC |
| P7-rbcLB_s | CAAGCAGAAGACGGCATACGAGATGTGACTGGAGTTCAGACGTGTGCTCTTCCGATCT | TTACTGC | AACCYTCTTCAAAAAGGTC |
| P7-rbcLB_t | CAAGCAGAAGACGGCATACGAGATGTGACTGGAGTTCAGACGTGTGCTCTTCCGATCT | GTTT | AACCYTCTTCAAAAAGGTC |
| P7-rbcLB_u | CAAGCAGAAGACGGCATACGAGATGTGACTGGAGTTCAGACGTGTGCTCTTCCGATCT | GACTT | AACCYTCTTCAAAAAGGTC |
| P7-rbcLB_v | CAAGCAGAAGACGGCATACGAGATGTGACTGGAGTTCAGACGTGTGCTCTTCCGATCT | TACCAC | AACCYTCTTCAAAAAGGTC |
| P7-rbcLB_w | CAAGCAGAAGACGGCATACGAGATGTGACTGGAGTTCAGACGTGTGCTCTTCCGATCT | ACCATAA | AACCYTCTTCAAAAAGGTC |
| P7-rbcLB_x | CAAGCAGAAGACGGCATACGAGATGTGACTGGAGTTCAGACGTGTGCTCTTCCGATCT | GAA | AACCYTCTTCAAAAAGGTC |

**Supplementary Table 3:** Bee species detected in trap nests in 2019 and 2020 across Canada using the COI plant marker, number of detections, and number of sites per landcover type with that species.

| Bee Family | Bee Species | Total Detections | Crop Sites | Forest Sites | Urban Sites |
|---|---|---|---|---|---|
| **Colletidae** | *Hylaeus annulatus* | 6 | | 3 | 3 |
| | *Hylaeus pictipes* | 1 | | | 1 |
| **Megachilidae** | *Chelostoma rapunculi* | 5 | | 1 | 3 |
| | *Coelioxys funeraria* | 11 | 6 | | 5 |
| | *Coelioxys modesta* | 7 | 2 | 1 | 3 |
| | *Coelioxys moesta* | 4 | | 1 | |
| | *Coelioxys sayi* | 5 | | 1 | 4 |
| | *Heriades carinatus* | 95 | 16 | 12 | 27 |
| | *Hoplitis albifrons* | 2 | | 1 | 1 |
| | *Megachile angelarum* | 14 | 3 | 2 | 5 |
| | *Megachile campanulae* | 54 | 10 | 10 | 15 |
| | *Megachile centuncularis* | 168 | 20 | 13 | 34 |
| | *Megachile lapponica* | 6 | | 1 | 1 |
| | *Megachile mendica* | 266 | 22 | 18 | 39 |
| | *Megachile pugnata* | 9 | 3 | | 6 |
| | *Megachile relativa* | 43 | 9 | 5 | 12 |
| | *Megachile rotundata* | 156 | 18 | 13 | 33 |
| | *Megachile snowi* | 15 | 3 | 2 | 7 |
| | *Osmia albiventris* | 2 | | | 2 |
| | *Osmia bicornis bicornis* | 1 | | | 1 |
| | *Osmia caerulescens* | 23 | 5 | 3 | 8 |
| | *Osmia coloradensis* | 20 | 3 | 2 | 11 |
| | *Osmia dolerosa* | 7 | 1 | 1 | 3 |
| | *Osmia lignaria* | 92 | 11 | 7 | 25 |
| | *Osmia pumila* | 26 | 6 | 2 | 13 |
| | *Osmia taurus* | 81 | 16 | 8 | 19 |
| | *Osmia tersula* | 32 | 4 | 4 | 12 |
| | *Stelis coarctatus* | 5 | 3 | | 2 |

**Supplementary Figure 1:** Bee (A) and wasp (B) abundance by species and family in trap nests installed across Canada in 2019 and 2020 detected through COI metabarcoding.

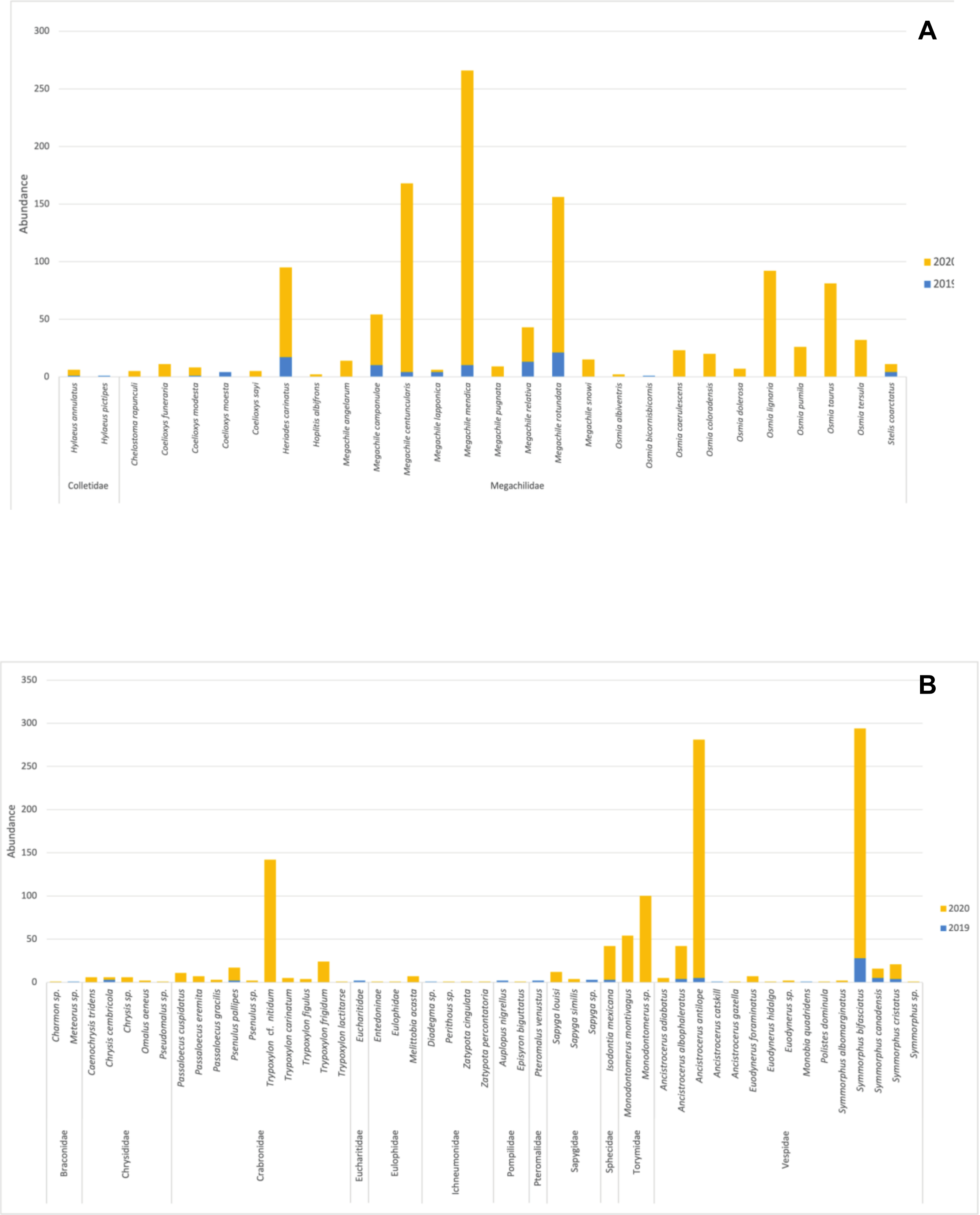

**Supplementary Figure 2:** Species richness variation for bee and wasp species found in each trap nest in 2019 and 2020 detected through COI metabarcoding. In each box, the thick horizontal bar is the median value, whilst the lower and upper edges represent the 25% and 75% quartiles respectively. Whiskers indicate the maximum and minimum values that are not outliers.

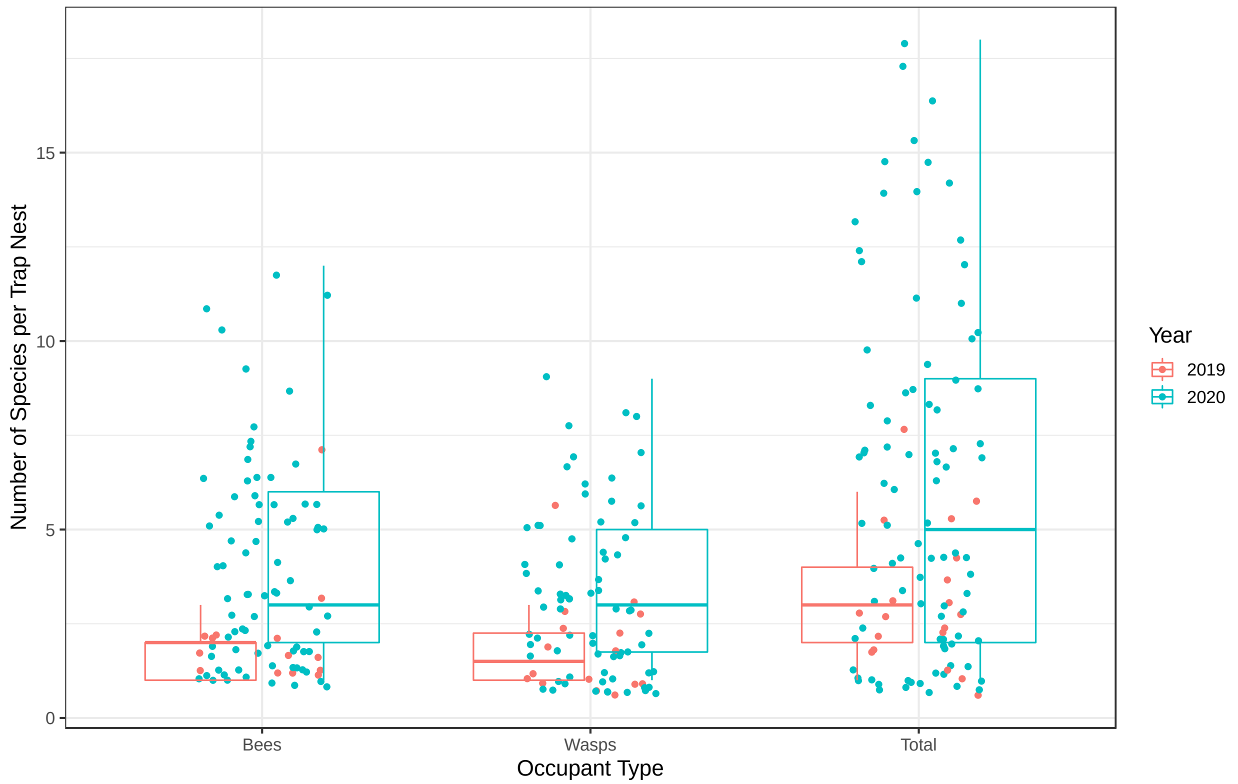

**Supplementary Table 4:** Plant genera detected in trap nests in 2019 and 2020 across Canada using the rbcL plant marker, number of detections, and number of sites per landcover type with that plant.

| Plant Order | Plant Family | Plant Genus | Detections | Crop Sites | Forest Sites | Urban Sites |
|---|---|---|---|---|---|---|
| **Alismatales** | Alismataceae | *Sagittaria* | 1 | 1 | | |
| | Potamogetonaceae | *Potamogeton* | 1 | 1 | | |
| **Apiales** | Apiaceae | *Aegopodium* | 6 | 3 | | 1 |
| | | *Berula* | 5 | 1 | 2 | 1 |
| | | *Cicuta* | 16 | 6 | 2 | 4 |
| | | *Conium* | 33 | 5 | 5 | 14 |
| | | *Daucus* | 89 | 14 | 7 | 20 |
| | | *Foeniculum* | 24 | 2 | 3 | 12 |
| | | *Osmorhiza* | 4 | 1 | | 2 |
| | | *Oxypolis* | 15 | 2 | 3 | 8 |
| | | *Perideridia* | 3 | | 1 | |
| | | *Podistera* | 1 | | | 1 |
| | | *Zizia* | 5 | 1 | 1 | 1 |
| | Araliaceae | *Aralia* | 3 | | 2 | 1 |
| **Asparagales** | Alliaceae | *Allium* | 16 | 1 | 3 | 11 |
| | Asparagaceae | *Camassia* | 1 | 1 | | |
| | | *Ornithogalum* | 1 | | 1 | |
| | | *Polygonatum* | 1 | | | 1 |
| | Iridaceae | *Sisyrinchium* | 1 | | | 1 |
| | Orchidaceae | *Cypripedium* | 1 | | 1 | |
| | | *Epipactis* | 2 | | | 2 |
| | | *Malaxis* | 1 | | | 1 |
| | Ruscaceae | *Maianthemum* | 1 | | 1 | |
| | Xanthorrhoeaceae | *Hemerocallis* | 19 | 3 | | 5 |
| **Asterales** | Asteraceae | *Acanthospermum* | 1 | | 1 | |
| | | *Achillea* | 109 | 9 | 11 | 24 |
| | | *Ageratina* | 2 | 1 | 1 | |
| | | *Ambrosia* | 115 | 15 | 9 | 25 |
| | | *Antennaria* | 8 | 2 | 2 | 2 |
| **Asterales** | Asteraceae | *Anthemis* | 12 | 2 | | 3 |
| | | *Arctium* | 121 | 13 | 10 | 35 |

| Plant Order | Plant Family | Plant Genus | Detections | Crop Sites | Forest Sites | Urban Sites |
|---|---|---|---|---|---|---|
| | | *Artemisia* | 5 | 1 | | 2 |
| | | *Baccharis* | 2 | | 1 | 1 |
| | | *Bellis* | 3 | | 1 | 1 |
| | | *Bidens* | 19 | 1 | 1 | 9 |
| | | *Calendula* | 21 | | 2 | 7 |
| | | *Carthamus* | 1 | | | 1 |
| | | *Centaurea* | 37 | 6 | 3 | 9 |
| | | *Chondrilla* | 22 | 7 | 2 | 5 |
| | | *Chrysanthemum* | 2 | 1 | | |
| | | *Cichorium* | 42 | 7 | 1 | 7 |
| | | *Cirsium* | 121 | 15 | 5 | 23 |
| | | *Cosmos* | 11 | 1 | 1 | 5 |
| | | *Crepis* | 82 | 11 | 12 | 26 |
| | | *Crocidium* | 8 | | 2 | 2 |
| | | *Cyclachaena* | 3 | 1 | | 2 |
| | | *Dittrichia* | 1 | | | 1 |
| | | *Doellingeria* | 1 | | | 1 |
| | | *Echinacea* | 32 | 3 | 4 | 14 |
| | | *Echinops* | 40 | 8 | 2 | 16 |
| | | *Eclipta* | 10 | 2 | 1 | 4 |
| | | *Erigeron* | 70 | 10 | 4 | 20 |
| | | *Eutrochium* | 111 | 14 | 11 | 30 |
| | | *Gaillardia* | 45 | 7 | 4 | 17 |
| | | *Galinsoga* | 5 | 1 | 1 | 1 |
| | | *Guizotia* | 9 | | 1 | 6 |
| | | *Helenium* | 3 | | | 3 |
| | | *Helianthus* | 24 | 3 | 2 | 9 |
| | | *Heliopsis* | 23 | 1 | 2 | 11 |
| | | *Hulteniella* | 9 | 2 | 1 | 6 |
| | | *Hypochaeris* | 26 | 5 | 3 | 12 |
| | | *Inula* | 4 | | | 2 |
| | | *Iva* | 1 | | | 1 |
| | | *Jacobaea* | 2 | | | 2 |
| **Asterales** | Asteraceae | *Lactuca* | 65 | 6 | 6 | 12 |
| | | *Lasthenia* | 2 | | | 2 |

| Plant Order | Plant Family | Plant Genus | Detections | Crop Sites | Forest Sites | Urban Sites |
|---|---|---|---|---|---|---|
| | | *Leontodon* | 41 | 5 | 3 | 14 |
| | | *Leucanthemum* | 83 | 9 | 5 | 23 |
| | | *Liatris* | 37 | 9 | 2 | 10 |
| | | *Logfia* | 6 | | | 3 |
| | | *Lygodesmia* | 28 | 8 | | 8 |
| | | *Madia* | 1 | | | 1 |
| | | *Microseris* | 89 | 10 | 7 | 26 |
| | | *Packera* | 1 | | | 1 |
| | | *Picradeniopsis* | 1 | | | 1 |
| | | *Pseudognaphalium* | 26 | 2 | 4 | 5 |
| | | *Pyrrocoma* | 135 | 15 | 6 | 29 |
| | | *Ratibida* | 123 | 16 | 8 | 28 |
| | | *Rhaponticum* | 4 | 1 | | 2 |
| | | *Rudbeckia* | 90 | 12 | 4 | 26 |
| | | *Senecio* | 9 | | 2 | 2 |
| | | *Shinnersoseris* | 3 | | | 1 |
| | | *Silphium* | 3 | | | 2 |
| | | *Solidago* | 4 | 2 | | 1 |
| | | *Sonchus* | 340 | 22 | 17 | 47 |
| | | *Stephanomeria* | 6 | 5 | | |
| | | *Symphyotrichum* | 143 | 15 | 8 | 36 |
| | | *Tagetes* | 28 | 4 | 3 | 9 |
| | | *Tanacetum* | 48 | 6 | 2 | 11 |
| | | *Taraxacum* | 1 | | 1 | |
| | | *Tragopogon* | 39 | 7 | 1 | 16 |
| | | *Tripleurospermum* | 22 | 2 | 1 | 9 |
| | | *Tussilago* | 10 | 3 | 2 | 3 |
| | Campanulaceae | *Campanula* | 178 | 18 | 16 | 41 |
| | | *Githopsis* | 5 | 1 | | 2 |
| | Menyanthaceae | *Menyanthes* | 2 | 1 | | 1 |
| | | *Nephrophyllidium* | 6 | 2 | | 3 |
| **Boraginales** | Boraginaceae | *Anchusa* | 5 | | | 1 |
| | | *Asperugo* | 1 | | | 1 |
| **Boraginales** | Boraginaceae | *Buglossoides* | 9 | 2 | | |
| | | *Hydrophyllum* | 1 | | | 1 |

| Plant Order | Plant Family | Plant Genus | Detections | Crop Sites | Forest Sites | Urban Sites |
|---|---|---|---|---|---|---|
| | | *Lappula* | 1 | 1 | | |
| | | *Lithospermum* | 10 | 4 | 2 | 4 |
| | | *Mertensia* | 1 | | 1 | |
| | | *Myosotis* | 8 | | 1 | 5 |
| **Brassicales** | Brassicaceae | *Arabidopsis* | 1 | | | 1 |
| | | *Berteroa* | 12 | | | 6 |
| | | *Brassica* | 35 | 7 | 2 | 16 |
| | | *Bunias* | 12 | 3 | 1 | 4 |
| | | *Cakile* | 11 | 2 | 1 | 4 |
| | | *Camelina* | 2 | | | 2 |
| | | *Cardamine* | 5 | | | 4 |
| | | *Draba* | 1 | | | 1 |
| | | *Erysimum* | 12 | 3 | | 7 |
| | | *Hesperis* | 7 | | | 5 |
| | | *Lepidium* | 9 | 3 | | 3 |
| | | *Lunaria* | 7 | 1 | 1 | 3 |
| | | *Physaria* | 1 | 1 | | |
| | | *Rorippa* | 5 | | | 4 |
| | | *Sinapis* | 1 | | | 1 |
| | | *Sisymbrium* | 58 | 9 | 10 | 15 |
| | | *Thlaspi* | 70 | 12 | 8 | 21 |
| **Caryophyllales** | Amaranthaceae | *Kochia* | 6 | 3 | | 3 |
| | | *Polycnemum* | 2 | | 1 | 1 |
| | | *Salsola* | 15 | 1 | | |
| | | *Spinacia* | 2 | 1 | 1 | |
| | Caryophyllaceae | *Cerastium* | 4 | 2 | | 2 |
| | | *Moehringia* | 1 | | 1 | |
| | | *Silene* | 1 | | 1 | |
| | | *Spergularia* | 1 | | | 1 |
| | | *Stellaria* | 1 | | | 1 |
| | Chenopodiaceae | *Atriplex* | 4 | 1 | | 2 |
| | | *Chenopodium* | 38 | 9 | 4 | 15 |
| | Nyctaginaceae | *Mirabilis* | 1 | | | 1 |
| **Caryophyllales** | Polygonaceae | *Fagopyrum* | 2 | | 1 | 1 |
| | | *Fallopia* | 4 | 1 | | |

| Plant Order | Plant Family | Plant Genus | Detections | Crop Sites | Forest Sites | Urban Sites |
|---|---|---|---|---|---|---|
| | | *Persicaria* | 19 | 3 | 3 | 8 |
| | | *Polygonum* | 60 | 7 | 3 | 15 |
| | | *Rheum* | 1 | | | 1 |
| | | *Rumex* | 13 | 2 | 1 | 7 |
| | Portulacaceae | *Portulaca* | 4 | 3 | | 1 |
| | Tamaricaceae | *Tamarix* | 15 | 1 | 2 | 5 |
| **Celastrales** | Celastraceae | *Euonymus* | 1 | | | 1 |
| **Cornales** | Cornaceae | *Cornus* | 190 | 20 | 16 | 40 |
| | | *Nyssa* | 1 | | | 1 |
| | Hydrangeaceae | *Hydrangea* | 11 | | | 4 |
| | | *Philadelphus* | 28 | 5 | 2 | 11 |
| **Cucurbitales** | Cucurbitaceae | *Citrullus* | 51 | 8 | 6 | 23 |
| | | *Cucumis* | 31 | 5 | 2 | 15 |
| | | *Cucurbita* | 51 | 8 | 6 | 18 |
| | | *Echinocystis* | 28 | 8 | 1 | 9 |
| | | *Marah* | 2 | | | 2 |
| **Dipsacales** | Adoxaceae | *Sambucus* | 43 | 7 | 3 | 15 |
| | | *Viburnum* | 43 | 3 | 8 | 15 |
| | Caprifoliaceae | *Diervilla* | 57 | 9 | 7 | 22 |
| | | *Dipsacus* | 2 | | | 2 |
| | | *Kolkwitzia* | 8 | 1 | 1 | 4 |
| | | *Linnaea* | 3 | 1 | | 2 |
| | | *Lonicera* | 85 | 14 | 9 | 24 |
| | | *Succisella* | 1 | | | 1 |
| | | *Triosteum* | 3 | 1 | 1 | 1 |
| | | *Valeriana* | 21 | 3 | | 7 |
| | Valerianaceae | *Plectritis* | 1 | | 1 | |
| **Equisetales** | Equisetaceae | *Equisetum* | 10 | 2 | 2 | 6 |
| **Ericales** | Balsaminaceae | *Impatiens* | 14 | 4 | 1 | 8 |
| | Ericaceae | *Arbutus* | 5 | 1 | 2 | 2 |
| | | *Arctostaphylos* | 1 | | | 1 |
| | | *Calluna* | 6 | | 2 | 1 |
| | | *Chamaedaphne* | 2 | | 2 | |
| **Ericales** | Ericaceae | *Gaultheria* | 1 | | | 1 |
| | | *Kalmia* | 13 | 1 | 2 | 2 |

| Plant Order | Plant Family | Plant Genus | Detections | Crop Sites | Forest Sites | Urban Sites |
|---|---|---|---|---|---|---|
| | | *Orthilia* | 1 | | | 1 |
| | | *Pyrola* | 3 | | 2 | |
| | | *Rhododendron* | 14 | | 3 | 3 |
| | | *Vaccinium* | 10 | | 3 | 3 |
| | Polemoniaceae | *Microsteris* | 3 | 1 | | |
| | | *Phlox* | 7 | | 2 | 3 |
| | Primulaceae | *Lysimachia* | 12 | 4 | 1 | 3 |
| | Sarraceniaceae | *Darlingtonia* | 1 | | | 1 |
| **Fabales** | Fabaceae | *Acmispon* | 5 | 1 | | 2 |
| | | *Astragalus* | 42 | 6 | 5 | 17 |
| | | *Baptisia* | 1 | | | 1 |
| | | *Caragana* | 92 | 10 | 11 | 26 |
| | | *Cladrastis* | 4 | 2 | 1 | 1 |
| | | *Coronilla* | 97 | 15 | 11 | 28 |
| | | *Cytisus* | 8 | | 3 | 5 |
| | | *Dalea* | 1 | 1 | | |
| | | *Desmodium* | 1 | 1 | | |
| | | *Genista* | 1 | | 1 | |
| | | *Gleditsia* | 24 | 3 | 5 | 9 |
| | | *Glycine* | 90 | 14 | 7 | 27 |
| | | *Glycyrrhiza* | 2 | 2 | | |
| | | *Hedysarum* | 1 | | | 1 |
| | | *Lathyrus* | 108 | 11 | 12 | 28 |
| | | *Lotus* | 196 | 21 | 14 | 40 |
| | | *Lupinus* | 2 | 1 | | 1 |
| | | *Medicago* | 200 | 19 | 11 | 38 |
| | | *Melilotus* | 279 | 20 | 15 | 45 |
| | | *Oxytropis* | 1 | | | 1 |
| | | *Phaseolus* | 3 | 1 | 1 | 1 |
| | | *Pisum* | 24 | 4 | 3 | 10 |
| | | *Robinia* | 76 | 14 | 9 | 26 |
| | | *Strophostyles* | 3 | 1 | | 1 |
| | | *Trifolium* | 295 | 21 | 17 | 42 |
| **Fabales** | Fabaceae | *Trigonella* | 1 | 1 | | |
| | | *Vicia* | 129 | 16 | 10 | 30 |

| Plant Order | Plant Family | Plant Genus | Detections | Crop Sites | Forest Sites | Urban Sites |
|---|---|---|---|---|---|---|
| Fagales | Betulaceae | *Alnus* | 24 | 3 | 7 | 7 |
| | | *Betula* | 1 | | | 1 |
| | | *Carpinus* | 2 | | | 2 |
| | | *Corylus* | 130 | 18 | 13 | 35 |
| | | *Ostrya* | 24 | 6 | 4 | 9 |
| | Fagaceae | *Fagus* | 64 | 9 | 4 | 21 |
| | | *Quercus* | 168 | 16 | 14 | 34 |
| | Juglandaceae | *Carya* | 61 | 11 | 6 | 24 |
| | Myricaceae | *Myrica* | 6 | | 2 | 1 |
| Gentianales | Apocynaceae | *Apocynum* | 1 | | 1 | |
| | | *Asclepias* | 4 | 2 | | 1 |
| | | *Vinca* | 3 | | 1 | 2 |
| | Rubiaceae | *Galium* | 4 | 3 | 1 | |
| Geraniales | Geraniaceae | *Geranium* | 5 | | | 4 |
| Lamiales | Bignoniaceae | *Campsis* | 77 | 11 | 8 | 28 |
| | | *Catalpa* | 2 | | | 2 |
| | Lamiaceae | *Agastache* | 30 | 5 | 3 | 14 |
| | | *Ajuga* | 8 | 1 | 1 | 4 |
| | | *Blephilia* | 4 | 1 | | 2 |
| | | *Collinsonia* | 3 | 1 | | 1 |
| | | *Glechoma* | 31 | 7 | 3 | 13 |
| | | *Hedeoma* | 9 | | 2 | 7 |
| | | *Hyssopus* | 1 | | | 1 |
| | | *Lamiastrum* | 1 | | | 1 |
| | | *Lamium* | 18 | 4 | 2 | 11 |
| | | *Leonurus* | 11 | 2 | | 3 |
| | | *Lycopus* | 8 | 3 | | 5 |
| | | *Marrubium* | 1 | | | 1 |
| | | *Monarda* | 7 | 1 | | 3 |
| | | *Nepeta* | 8 | 3 | | 2 |
| | | *Physostegia* | 5 | 2 | | 2 |
| | | *Prunella* | 34 | 8 | 3 | 14 |
| | | *Scutellaria* | 1 | | | 1 |
| Lamiales | Lamiaceae | *Teucrium* | 2 | | | 1 |
| | | *Thymus* | 61 | 12 | 5 | 25 |

| Plant Order | Plant Family | Plant Genus | Detections | Crop Sites | Forest Sites | Urban Sites |
|---|---|---|---|---|---|---|
| | Martyniaceae | *Proboscidea* | 1 | | | 1 |
| | Oleaceae | *Fraxinus* | 123 | 16 | 11 | 31 |
| | | *Ligustrum* | 37 | 4 | 3 | 18 |
| | | *Syringa* | 40 | 3 | 7 | 18 |
| | Plantaginaceae | *Antirrhinum* | 1 | | | 1 |
| | | *Chaenorhinum* | 6 | | | 3 |
| | | *Chelone* | 3 | | 1 | 2 |
| | | *Cymbalaria* | 6 | | | 1 |
| | | *Digitalis* | 1 | | 1 | |
| | | *Linaria* | 2 | | | 2 |
| | | *Plantago* | 66 | 12 | 6 | 20 |
| | | *Veronica* | 68 | 13 | 6 | 20 |
| | | *Veronicastrum* | 7 | 1 | 2 | 2 |
| | Scrophulariaceae | *Buddleja* | 4 | | | 4 |
| | Verbenaceae | *Verbena* | 2 | 2 | | |
| **Laurales** | Lauraceae | *Lindera* | 1 | 1 | | |
| **Liliales** | Melanthiaceae | *Trillium* | 4 | | 1 | 1 |
| **Magnoliales** | Magnoliaceae | *Magnolia* | 2 | 1 | | 1 |
| **Malpighiales** | Euphorbiaceae | *Acalypha* | 8 | 2 | 1 | 4 |
| | | *Euphorbia* | 32 | 4 | 4 | 16 |
| | Hypericaceae | *Hypericum* | 87 | 8 | 6 | 23 |
| | Salicaceae | *Populus* | 196 | 19 | 13 | 44 |
| | | *Salix* | 335 | 21 | 17 | 48 |
| | Violaceae | *Viola* | 40 | 7 | 6 | 16 |
| **Malvales** | Malvaceae | *Abutilon* | 51 | 10 | 6 | 19 |
| | | *Althaea* | 23 | 3 | 2 | 9 |
| | | *Lavatera* | 7 | 1 | 1 | 3 |
| | | *Malva* | 213 | 18 | 15 | 40 |
| | | *Sida* | 2 | 1 | | 1 |
| | | *Sidalcea* | 2 | 1 | | 1 |
| | | *Tilia* | 152 | 13 | 13 | 34 |
| **Myrtales** | Lythraceae | *Decodon* | 2 | 1 | | 1 |
| | | *Lythrum* | 36 | 7 | 2 | 7 |
| **Myrtales** | Lythraceae | *Rotala* | 1 | | | 1 |
| | Onagraceae | *Chamerion* | 189 | 18 | 14 | 39 |

| Plant Order | Plant Family | Plant Genus | Detections | Crop Sites | Forest Sites | Urban Sites |
|---|---|---|---|---|---|---|
| | | *Circaea* | 5 | 2 | | 3 |
| | | *Epilobium* | 36 | 5 | 6 | 17 |
| | | *Ludwigia* | 14 | 3 | 1 | 3 |
| | | *Neoholmgrenia* | 2 | | | 2 |
| | | *Oenothera* | 3 | | 2 | 1 |
| **Nymphaeales** | Nymphaeaceae | *Nymphaea* | 2 | | 1 | |
| **Oxalidales** | Oxalidaceae | *Oxalis* | 21 | 3 | | 10 |
| **Pinales** | Cupressaceae | *Juniperus* | 26 | 2 | 4 | 7 |
| | | *Thuja* | 2 | | 1 | 1 |
| | Pinaceae | *Abies* | 14 | | 2 | 3 |
| | | *Larix* | 51 | 8 | 6 | 19 |
| | | *Picea* | 90 | 10 | 9 | 29 |
| | | *Pinus* | 238 | 22 | 15 | 45 |
| | | *Tsuga* | 21 | 1 | 3 | 5 |
| **Poales** | Cyperaceae | *Carex* | 11 | 2 | 2 | 4 |
| | | *Cyperus* | 1 | | | 1 |
| | | *Scirpus* | 2 | | 1 | 1 |
| | Juncaceae | *Juncus* | 3 | 2 | 1 | |
| | Poaceae | *Agrostis* | 22 | 4 | 3 | 8 |
| | | *Andropogon* | 26 | 5 | 2 | 13 |
| | | *Arrhenatherum* | 16 | 1 | 3 | 7 |
| | | *Briza* | 4 | | 2 | 2 |
| | | *Bromus* | 97 | 11 | 10 | 28 |
| | | *Deschampsia* | 6 | | 1 | 4 |
| | | *Digitaria* | 6 | 2 | | 4 |
| | | *Echinochloa* | 1 | | | 1 |
| | | *Elymus* | 27 | 5 | 4 | 10 |
| | | *Glyceria* | 2 | 1 | 1 | |
| | | *Hordeum* | 4 | 1 | 1 | 2 |
| | | *Lolium* | 47 | 12 | 5 | 18 |
| | | *Panicum* | 2 | | | 2 |
| | | *Phalaris* | 14 | 3 | 2 | 2 |
| | | *Phragmites* | 3 | 1 | | 1 |
| **Poales** | Poaceae | *Poa* | 100 | 14 | 8 | 28 |
| | | *Polypogon* | 2 | | 1 | |

| Plant Order | Plant Family | Plant Genus | Detections | Crop Sites | Forest Sites | Urban Sites |
|---|---|---|---|---|---|---|
| | | *Pseudoroegneria* | 1 | 1 | | |
| | | *Secale* | 25 | 4 | 4 | 10 |
| | | *Setaria* | 4 | 1 | 1 | 2 |
| | | *Sorghastrum* | 15 | 2 | 2 | 7 |
| | | *Trisetum* | 9 | 2 | 2 | 2 |
| | | *Vahlodea* | 9 | 2 | | 4 |
| | | *Vulpia* | 3 | 1 | 1 | 1 |
| | Typhaceae | *Typha* | 10 | 1 | 1 | 7 |
| **Proteales** | Platanaceae | *Platanus* | 5 | | 1 | 3 |
| **Ranunculales** | Berberidaceae | *Berberis* | 2 | | | 1 |
| | | *Mahonia* | 7 | 1 | 1 | 1 |
| | Papaveraceae | *Chelidonium* | 4 | | | 4 |
| | | *Eschscholzia* | 4 | | | 2 |
| | | *Papaver* | 1 | | | 1 |
| | Ranunculaceae | *Aconitum* | 1 | | | 1 |
| | | *Actaea* | 3 | | 1 | 1 |
| | | *Anemone* | 11 | 4 | 2 | 1 |
| | | *Aquilegia* | 2 | | | 2 |
| | | *Caltha* | 22 | 3 | 4 | 12 |
| | | *Clematis* | 72 | 11 | 10 | 26 |
| | | *Ranunculus* | 4 | 1 | | 2 |
| | | *Thalictrum* | 4 | 2 | 1 | 1 |
| | | *Trollius* | 2 | 1 | | 1 |
| **Rosales** | Cannabaceae | *Celtis* | 22 | 5 | 3 | 11 |
| | | *Humulus* | 16 | 2 | 2 | 10 |
| | Elaeagnaceae | *Elaeagnus* | 9 | 2 | 2 | 4 |
| | Moraceae | *Morus* | 24 | 6 | 2 | 7 |
| | Rhamnaceae | *Ceanothus* | 7 | 1 | 3 | 3 |
| | | *Rhamnus* | 108 | 14 | 15 | 28 |
| | Rosaceae | *Aruncus* | 26 | 4 | 2 | 14 |
| | | *Comarum* | 20 | 4 | 3 | 11 |
| | | *Crataegus* | 249 | 21 | 18 | 44 |
| | | *Dasiphora* | 62 | 14 | 5 | 15 |
| **Rosales** | Rosaceae | *Dryas* | 3 | | 1 | 2 |
| | | *Drymocallis* | 12 | 3 | 2 | 3 |

| Plant Order | Plant Family | Plant Genus | Detections | Crop Sites | Forest Sites | Urban Sites |
|---|---|---|---|---|---|---|
| | | *Fragaria* | 226 | 20 | 15 | 43 |
| | | *Geum* | 288 | 19 | 17 | 43 |
| | | *Holodiscus* | 4 | | | 2 |
| | | *Physocarpus* | 10 | 2 | | 5 |
| | | *Potentilla* | 180 | 16 | 13 | 37 |
| | | *Prunus* | 157 | 17 | 13 | 35 |
| | | *Pyrus* | 13 | 4 | 3 | 5 |
| | | *Rosa* | 207 | 17 | 15 | 36 |
| | | *Rubus* | 129 | 16 | 16 | 30 |
| | | *Sanguisorba* | 1 | | | 1 |
| | | *Sorbaria* | 9 | | 1 | 5 |
| | | *Spiraea* | 72 | 8 | 5 | 28 |
| | Ulmaceae | *Ulmus* | 104 | 13 | 11 | 27 |
| | Urticaceae | *Boehmeria* | 3 | 1 | | 1 |
| | | *Laportea* | 5 | 3 | | |
| | | *Urtica* | 12 | 6 | 2 | 1 |
| **Santalales** | Santalaceae | *Comandra* | 2 | | 1 | |
| **Sapindales** | Anacardiaceae | *Rhus* | 84 | 10 | 11 | 29 |
| | Rutaceae | *Zanthoxylum* | 32 | 4 | 4 | 18 |
| | Sapindaceae | *Acer* | 232 | 18 | 15 | 44 |
| | | *Aesculus* | 12 | | 2 | 4 |
| | Simaroubaceae | *Ailanthus* | 14 | 2 | 2 | 7 |
| **Saxifragales** | Crassulaceae | *Hylotelephium* | 2 | | | 2 |
| | | *Phedimus* | 24 | 6 | 4 | 12 |
| | | *Sedum* | 1 | | | 1 |
| | | *Sempervivum* | 1 | | | 1 |
| | Grossulariaceae | *Ribes* | 14 | 2 | 2 | 8 |
| | Paeoniaceae | *Paeonia* | 1 | 1 | | |
| | Saxifragaceae | *Heuchera* | 2 | | 1 | 1 |
| **Solanales** | Convolvulaceae | *Convolvulus* | 115 | 14 | 11 | 33 |
| | | *Cuscuta* | 8 | 1 | 1 | 5 |
| | | *Ipomoea* | 1 | | | 1 |
| | Solanaceae | *Lycium* | 2 | | | 2 |
| **Solanales** | Solanaceae | *Nicotiana* | 1 | | | 1 |
| | | *Physalis* | 85 | 10 | 8 | 31 |

| Plant Order | Plant Family | Plant Genus | Detections | Crop Sites | Forest Sites | Urban Sites |
|---|---|---|---|---|---|---|
| | | *Scopolia* | 10 | 2 | | 5 |
| **Vitales** | Vitaceae | *Parthenocissus* | 290 | 22 | 16 | 48 |
| | | *Vitis* | 131 | 16 | 10 | 29 |

**Supplementary Table 5:** Cavity-nesting bees and wasps co-habiting trap nest tubes in 2019 and 2020. A slash between species shows that they were additionally found in the same tube; numbers in brackets denote multiple occurrences of co-habitation.

| Species | Co-habiting with |
|---|---|
| ***Ancistrocerus adiabatus*** | *Ancistrocerus antilope / Megachile rotundata* |
| | *Chelostoma rapunculi / Symmorphus canadensis* |
| | *Isodontia mexicana* |
| | *Megachile rotundata* |
| | *Osmia lignaria* |
| ***Ancistrocerus albophaleratus*** | *Chelostoma rapunculi / Megachile centuncularis* |
| | *Isodontia mexicana* |
| | *Megachile centuncularis / Osmia lignaria* |
| | *Megachile mendica / Megachile relativa* |
| | *Megachile relativa* |
| | *Osmia lignaria / Osmia taurus / Symmorphus bifasciatus* |
| | *Osmia lignaria* |
| | *Symmorphus bifasciatus* |
| | *Trypoxylon* cf. *nitidum* |
| ***Ancistrocerus antilope*** | *Ancistrocerus adiabatus / Megachile rotundata* |
| | *Euodynerus* sp. |
| | *Isodontia mexicana* (2) |
| | *Megachile campanulae / Symmorphus bifasciatus* |
| | *Megachile centuncularis* |
| | *Osmia lignaria / Symmorphus cristatus* |
| | *Osmia lignaria* |
| | *Osmia pumila* |
| | *Symmorphus bifasciatus* (2) |
| | *Trypoxylon* cf. *nitidum* |
| ***Chelostoma rapunculi*** | *Ancistrocerus adiabatus / Symmorphus canadensis* |
| | *Ancistrocerus albophaleratus / Megachile centuncularis* |
| | *Megachile rotundata* |
| ***Euodynerus foraminatus*** | *Trypoxylon lactitarse* |
| ***Euodynerus sp.*** | *Osmia taurus* |
| | *Ancistrocerus antilope* |
| ***Heriades carinatus*** | *Isodontia mexicana* (3) |
| | *Megachile centuncularis / Osmia pumila* |
| | *Megachile centuncularis* |
| | *Megachile mendica / Megachile rotundata / Trypoxylon* cf. *nitidum* |
| | *Megachile mendica / Osmia lignaria* |
| | *Megachile mendica* |
| | *Megachile rotundata* (2) |
| | *Osmia caerulescens* |
| | *Osmia lignaria* (2) |
| | *Psenulus pallipes* |
| | *Trypoxylon* cf. *nitidum* |
| ***Hylaeus annulatus*** | *Megachile campanulae* |
| ***Isodontia mexicana*** | *Ancistrocerus adiabatus* |
| ***Isodontia mexicana*** | *Ancistrocerus albophaleratus* |
| | *Ancistrocerus antilope* (2) |
| | *Heriades carinatus* (3) |

| Species | Co-habiting with |
|---|---|
| | *Megachile mendica* (3) |
| | *Megachile pugnata / Megachile rotundata* |
| | *Megachile relativa* |
| | *Osmia caerulescens / Osmia lignaria* |
| | *Osmia lignaria* (2) |
| | *Osmia pumila / Symmorphus bifasciatus* |
| | *Osmia pumila* |
| | *Osmia taurus / Symmorphus canadensis* |
| | *Trypoxylon* cf. *nitidum / Trypoxylon frigidum* |
| | *Trypoxylon* cf. *nitidum* |
| ***Megachile angelarum*** | *Megachile campanulae / Symmorphus cristatus* |
| | *Megachile campanulae* (4) |
| | *Osmia lignaria / Trypoxylon* cf. *nitidum* |
| ***Megachile campanulae*** | *Ancistrocerus antilope / Symmorphus bifasciatus* |
| | *Hylaeus annulatus* |
| | *Megachile angelarum / Symmorphus cristatus* |
| | *Megachile angelarum* (4) |
| | *Megachile mendica* |
| | *Megachile relativa* (2) |
| | *Megachile rotundata* |
| | *Osmia lignaria* |
| | *Osmia taurus / Trypoxylon carinatum / Trypoxylon* cf. *nitidum* |
| ***Megachile centuncularis*** | *Ancistrocerus albophaleratus / Chelostoma rapunculi* |
| | *Ancistrocerus albophaleratus / Osmia lignaria* |
| | *Ancistrocerus antilope* |
| | *Heriades carinatus / Osmia pumila* |
| | *Heriades carinatus* |
| | *Megachile mendica / Osmia taurus* |
| | *Megachile rotundata* (5) |
| | *Osmia caerulescens / Trypoxylon* cf. *nitidum* |
| | *Osmia lignaria* |
| | *Osmia pumila* |
| | *Osmia taurus / Trypoxylon figulus* |
| ***Megachile lapponica*** | *Megachile mendica* |
| | *Megachile relativa* |
| ***Megachile mendica*** | *Ancistrocerus albophaleratus / Megachile relativa* |
| | *Heriades carinatus / Megachile rotundata / Trypoxylon* cf. *nitidum* |
| | *Heriades carinatus / Osmia lignaria* |
| | *Heriades carinatus* |
| | *Isodontia mexicana* (3) |
| | *Megachile campanulae* |
| | *Megachile centuncularis / Osmia taurus* |
| | *Megachile lapponica* |
| | *Megachile rotundata / Osmia taurus* |
| | *Megachile rotundata* |
| | *Osmia caerulescens* |
| | *Osmia dolerosa / Osmia pumila / Osmia taurus* |
| ***Megachile mendica*** | *Osmia lignaria* (2) |
| | *Osmia pumila* |
| | *Symmorphus bifasciatus / Symmorphus cristatus* |

| Species | Co-habiting with |
|---|---|
| | *Symmorphus bifasciatus* (3) |
| | *Symmorphus canadensis* |
| | *Trypoxylon frigidum* |
| ***Megachile pugnata*** | *Isodontia mexicana / Megachile rotundata* |
| | *Megachile relativa* |
| | *Osmia coloradensis* |
| | *Osmia lignaria* (2) |
| ***Megachile relativa*** | *Ancistrocerus albophaleratus / Megachile mendica* |
| | *Ancistrocerus albophaleratus* |
| | *Isodontia mexicana* |
| | *Megachile campanulae* (2) |
| | *Megachile lapponica* |
| | *Megachile pugnata* |
| | *Megachile rotundata / Osmia taurus* |
| | *Megachile rotundata / Symmorphus bifasciatus* |
| | *Megachile rotundata* (2) |
| | *Osmia caerulescens* |
| | *Osmia lignaria* (3) |
| | *Osmia taurus* |
| | *Symmorphus bifasciatus* |
| | *Trypoxylon* cf. *nitidum* |
| ***Megachile rotundata*** | *Ancistrocerus adiabatus / Ancistrocerus antilope* |
| | *Ancistrocerus adiabatus* |
| | *Chelostoma rapunculi* |
| | *Heriades carinatus / Megachile mendica / Trypoxylon* cf. *nitidum* |
| | *Heriades carinatus* (2) |
| | *Isodontia mexicana / Megachile pugnata* |
| | *Megachile campanulae* |
| | *Megachile centuncularis* (5) |
| | *Megachile mendica / Osmia taurus* |
| | *Megachile mendica* |
| | *Megachile relativa / Osmia taurus* |
| | *Megachile relativa / Symmorphus bifasciatus* |
| | *Megachile relativa* (2) |
| | *Osmia caerulescens* (2) |
| | *Osmia coloradensis* |
| | *Osmia lignaria / Osmia taurus* |
| | *Osmia lignaria / Trypoxylon* cf. *nitidum* |
| | *Osmia lignaria* |
| | *Osmia pumila* (2) |
| | *Osmia taurus* |
| | *Osmia tersula* |
| | *Symmorphus bifasciatus / Trypoxylon* cf. *nitidum* |
| | *Symmorphus bifasciatus* (5) |
| | *Trypoxylon carinatum* |
| | *Trypoxylon* cf. *nitidum* (3) |
| ***Osmia caerulescens*** | *Heriades carinatus* |
| ***Osmia caerulescens*** | *Isodontia mexicana / Osmia lignaria* |
| | *Megachile centuncularis / Trypoxylon* cf. *nitidum* |
| | *Megachile mendica* |

| Species | Co-habiting with |
|---|---|
| | *Megachile relativa* |
| | *Megachile rotundata* (2) |
| | *Osmia coloradensis* |
| | *Osmia lignaria / Osmia taurus* |
| | *Osmia lignaria* (2) |
| | *Osmia tersula* |
| | *Symmorphus bifasciatus* |
| ***Osmia coloradensis*** | *Megachile pugnata* |
| | *Megachile rotundata* |
| | *Osmia caerulescens* |
| | *Osmia taurus / Trypoxylon frigidum* |
| ***Osmia dolerosa*** | *Megachile mendica / Osmia pumila / Osmia taurus* |
| | *Osmia taurus* |
| ***Osmia lignaria*** | *Ancistrocerus adiabatus* |
| | *Ancistrocerus albophaleratus / Megachile centuncularis* |
| | *Ancistrocerus albophaleratus / Osmia taurus / Symmorphus bifasciatus* |
| | *Ancistrocerus albophaleratus* |
| | *Ancistrocerus antilope / Symmorphus cristatus* |
| | *Ancistrocerus antilope* |
| | *Heriades carinatus / Megachile mendica* |
| | *Heriades carinatus* (2) |
| | *Isodontia mexicana / Osmia caerulescens* |
| | *Isodontia mexicana* (2) |
| | *Megachile angelarum / Trypoxylon* cf. *nitidum* |
| | *Megachile campanulae* |
| | *Megachile centuncularis* |
| | *Megachile mendica* (2) |
| | *Megachile pugnata* (2) |
| | *Megachile relativa* (3) |
| | *Megachile rotundata / Osmia taurus* |
| | *Megachile rotundata / Trypoxylon* cf. *nitidum* |
| | *Megachile rotundata* |
| | *Osmia caerulescens / Osmia taurus* |
| | *Osmia caerulescens* |
| | *Osmia taurus* |
| | *Osmia tersula* |
| | *Symmorphus bifasciatus / Trypoxylon* cf. *nitidum* |
| | *Symmorphus bifasciatus* |
| | *Trypoxylon frigidum* |
| ***Osmia pumila*** | *Ancistrocerus antilope* |
| | *Ancistrocerus antilope / Symmorphus bifasciatus* |
| | *Heriades carinatus / Megachile centuncularis* |
| | *Isodontia mexicana* |
| | *Megachile centuncularis* |
| | *Megachile mendica / Osmia dolerosa / Osmia taurus* |
| | *Megachile mendica* |
| ***Osmia pumila*** | *Megachile rotundata* (2) |
| | *Osmia taurus / Trypoxylon* cf. *nitidum* |
| | *Osmia taurus* |

| Species | Co-habiting with |
|---|---|
| | *Passaloecus gracilis* |
| ***Osmia taurus*** | *Ancistrocerus albophaleratus / Osmia lignaria / Symmorphus bifasciatus* |
| | *Euodynerus* sp. |
| | *Isodontia mexicana / Symmorphus canadensis* |
| | *Megachile campanulae / Trypoxylon carinatum / Trypoxylon* cf. *nitidum* |
| | *Megachile centuncularis / Megachile mendica* |
| | *Megachile centuncularis / Trypoxylon figulus* |
| | *Megachile mendica / Megachile rotundata* |
| | *Megachile mendica / Osmia dolerosa / Osmia pumila* |
| | *Megachile relativa / Megachile rotundata* |
| | *Megachile relativa* |
| | *Megachile rotundata / Osmia lignaria* |
| | *Megachile rotundata* |
| | *Osmia caerulescens / Osmia lignaria* |
| | *Osmia coloradensis / Trypoxylon frigidum* |
| | *Osmia dolerosa* |
| | *Osmia lignaria* |
| | *Osmia pumila / Trypoxylon* cf. *nitidum* |
| | *Osmia pumila* |
| | *Osmia tersula* |
| | *Symmorphus bifasciatus* |
| | *Trypoxylon* cf. *nitidum* |
| ***Osmia tersula*** | *Megachile rotundata* |
| | *Osmia caerulescens* |
| | *Osmia lignaria* |
| | *Osmia taurus* |
| ***Passaloecus gracilis*** | *Osmia pumila* |
| ***Psenulus pallipes*** | *Heriades carinatus* |
| ***Symmorphus bifasciatus*** | *Ancistrocerus albophaleratus / Osmia lignaria / Osmia taurus* |
| | *Ancistrocerus albophaleratus* |
| | *Ancistrocerus antilope / Megachile campanulae* |
| | *Ancistrocerus antilope / Osmia pumila* |
| | *Ancistrocerus antilope* (2) |
| | *Megachile mendica / Symmorphus cristatus* |
| | *Megachile mendica* (3) |
| | *Megachile relativa / Megachile rotundata* |
| | *Megachile relativa* |
| | *Megachile rotundata / Trypoxylon* cf. *nitidum* |
| | *Megachile rotundata* (5) |
| | *Osmia caerulescens* |
| | *Osmia lignaria / Trypoxylon* cf. *nitidum* |
| | *Osmia lignaria* |
| | *Osmia taurus* |
| | *Symmorphus canadensis* |
| | *Symmorphus cristatus / Trypoxylon* cf. *nitidum* |
| ***Symmorphus bifasciatus*** | *Symmorphus cristatus* (2) |
| ***Symmorphus canadensis*** | *Ancistrocerus adiabatus / Chelostoma rapunculi* |
| | *Isodontia mexicana / Osmia taurus* |

| Species | Co-habiting with |
|---|---|
| | *Megachile mendica* |
| | *Symmorphus bifasciatus* |
| ***Symmorphus cristatus*** | *Ancistrocerus antilope / Osmia lignaria* |
| | *Megachile angelarum / Megachile campanulae* |
| | *Megachile mendica / Symmorphus bifasciatus* |
| | *Symmorphus bifasciatus / Trypoxylon* cf. *nitidum* |
| | *Symmorphus bifasciatus* (2) |
| ***Trypoxylon carinatum*** | *Megachile campanulae / Osmia taurus / Trypoxylon* cf. *nitidum* |
| | *Megachile rotundata* |
| ***Trypoxylon* cf. *nitidum*** | *Ancistrocerus albophaleratus* |
| | *Ancistrocerus antilope* |
| | *Heriades carinatus / Megachile mendica / Megachile rotundata* |
| | *Heriades carinatus* |
| | *Isodontia mexicana / Trypoxylon frigidum* |
| | *Isodontia mexicana* |
| | *Megachile angelarum / Osmia lignaria* |
| | *Megachile campanulae / Osmia taurus / Trypoxylon carinatum* |
| | *Megachile centuncularis / Osmia caerulescens* |
| | *Megachile relativa* |
| | *Megachile rotundata / Osmia lignaria* |
| | *Megachile rotundata / Symmorphus bifasciatus* |
| | *Megachile rotundata* (3) |
| | *Osmia lignaria / Symmorphus bifasciatus* |
| | *Osmia pumila / Osmia taurus* |
| | *Osmia tersula* |
| | *Symmorphus bifasciatus / Symmorphus cristatus* |
| ***Trypoxylon figulus*** | *Megachile centuncularis / Osmia taurus* |
| | *Trypoxylon frigidum* (3) |
| ***Trypoxylon frigidum*** | *Isodontia mexicana / Trypoxylon* cf. *nitidum* |
| | *Megachile mendica* |
| | *Osmia coloradensis / Osmia taurus* |
| | *Osmia lignaria* |
| | *Trypoxylon figulus* (3) |
| ***Trypoxylon lactitarse*** | *Euodynerus foraminatus* |

**Supplementary Table 6:** Nesting material identified through visual observation in nests with predatory wasps and pollinating bees. Partial numbers denote the presence of a nesting material in a tube with multiple nesting materials. Bolded numbers represent nesting material reported in the literature. Numbers in brackets are nesting tubes which were singly occupied. The category of 'None' nesting material represents a tube with no visible contents other than larvae.

| Species | Cellophane | Grass | Leaves | Masticated leaves | Mud | Rocks/ sand | Straw | Tree Resin | None |
|---|---|---|---|---|---|---|---|---|---|
| ***Ancistrocerus adiabatus*** | | 1 | | 1 | **1 (1)** | | 1 | 1 | 1 |
| ***Ancistrocerus albophaleratus*** | | | 5 (1) | 1 | **14 (11)** | | 1 | | |
| ***Ancistrocerus antilope*** | | | 4.5 (4) | | **15.5 (11)** | | 2 | 5 (2) | 2 |
| ***Ancistrocerus catskill*** | | | | | **1 (1)** | | | | |
| ***Ancistrocerus gazella*** | | | | | **1 (1)** | | | | |
| ***Chelostoma rapunculi*** | | 1 | | | **5 (1)** | | | | |
| ***Euodynerus foraminatus*** | | | | | **5 (4)** | | | | |
| ***Euodynerus hidalgo*** | | | | | **1 (1)** | | | | |
| ***Euodynerus* sp.** | | 1 | 0.5 | | **0.5** | | | | |
| ***Heriades carinatus*** | 1 (1) | 4.5 (3) | 4 | 1.5 (1) | 12 (8) | | 1 | **22 (19)** | 2 (1) |
| ***Hoplitis albifrons*** | | | 1 (1) | | | | | | |
| ***Hylaeus annulatus*** | | | 1 (1) | | | | 1 (1) | 1 (1) | **3 (2)** |
| ***Isodontia mexicana*** | | 4.5 (2) | 6 (1) | 0.5 | 2 (1) | | **13 (5)** | 9 (8) | 3 (1) |
| ***Megachile angelarum*** | | | | | **1.5** | | 2 | **2.5** | |
| ***Megachile campanulae*** | | 1 (1) | 1 | | **5.5 (1)** | | 2 | **12.5 (8)** | 1 |
| ***Megachile centuncularis*** | | 1 | **21 (11)** | 1 (1) | 7 (5) | | 2 (2) | 4 (2) | 2 (2) |
| ***Megachile lapponica*** | | | **4 (2)** | | | | | | |
| ***Megachile mendica*** | | 1.5 | **21 (11)** | **0.5** | 16 (7) | | 1 (1) | 2 (1) | 1 |
| ***Megachile pugnata*** | | 1 | **2 (1)** | **0.5** | **5 (4)** | | | 0.5 | |
| ***Megachile relativa*** | | 1.5 (1.5) | **20.5 (9)** | | 4.5 (1.5) | | 1 | 3.5 (2) | |
| ***Megachile rotundata*** | | 3 (1) | **36 (20)** | 2 | 13 (4) | | 2 (1) | 8 (4) | 14 (8) |
| ***Monobia quadridens*** | | | | | **1 (1)** | | | | |
| ***Osmia caerulescens*** | | 5 (2.5) | 5 | **0.5 (0.5)** | 11 (6.5) | 0.5 (0.5) | 1 (1) | 2 (2) | 1 |
| ***Osmia coloradensis*** | | **5 (3)** | 1 | **2 (2)** | 1 (1) | | | 1 | 1 (1) |
| ***Osmia dolerosa*** | | 2 (1) | 1 | 1 (1) | 2 (2) | | | | |

| Species | Cellophane | Grass | Leaves | Masticated leaves | Mud | Rocks/ sand | Straw | Tree Resin | None |
|---|---|---|---|---|---|---|---|---|---|
| ***Osmia lignaria*** | | | 4.5 (1) | 2 | **32 (12)** | | | 11.5 (7) | 3 (1) |
| ***Osmia pumila*** | | **8 (5)** | 5.5 (2.5) | **0.5 (0.5)** | 5 (3) | | 2 | | 3 (1) |
| ***Osmia taurus*** | | 3 | 7.5 (2.5) | | **19.5 (8.5)** | | 1 | 2 (2) | |
| ***Osmia tersula*** | 1 (1) | **7 (6.5)** | 4 (4) | **0.5 (0.5)** | 10 (7.5) | 0.5 (0.5) | | 2 (2) | 1 |
| ***Passaloecus cuspidatus*** | | | 1 (1) | | 1 (1) | | | **3 (3)** | |
| ***Passaloecus eremita*** | | | | | 3 (3) | | | **4 (4)** | |
| ***Passaloecus gracilis*** | | | 1 | | | | | **1 (1)** | 1 (1) |
| ***Psenulus pallipes*** | 1 (1) | | | | 5 (5) | | | 1 | 4 (4) |
| ***Symmorphus albomarginatus*** | | | | | **1 (1)** | | | | |
| ***Symmorphus bifasciatus*** | 0.5 (0.5) | 5 (3) | 8.5 (1.5) | | **35 (23)** | | 1 (1) | 3 (2) | 7 (3) |
| ***Symmorphus canadensis*** | | 1 | 1 (1) | | **8 (6)** | | 1 | 1 (1) | |
| ***Symmorphus cristatus*** | | 1 (1) | 2 | | **7 (4)** | | 1 | | |
| ***Symmorphus* sp.** | | | 1 (1) | | | | | | |
| ***Trypoxylon carinatum*** | | | | 0.5 | **2.5 (1)** | | | 2 (2) | |
| ***Trypoxylon* cf. *nitidum*** | | 1 (1) | 12 (3) | 0.5 | **14 (7)** | 1 (1) | 2 | 2.5 (2) | 2 (1) |
| ***Trypoxylon figulus*** | | | 1 | | **3** | | | | |
| ***Trypoxylon frigidum*** | | 2.5 (1) | 1 | 0.5 | **7 (3)** | | | 2 (2) | |
| ***Trypoxylon lactitarse*** | | | | | **1** | | | | |

**Supplementary Table 7:** Number of unique plant families and genera foraged by each pollinator species at crop, forest, and urban sites.

| **Pollinator Species** | **Crop** | | **Forest** | | **Urban** | | **Total** | |
|---|---|---|---|---|---|---|---|---|
| | Plant Family | Plant Genus | Plant Family | Plant Genus | Plant Family | Plant Genus | Plant Family | Plant Genus |
| ***Chelostoma rapunculi*** | 0 | 0 | 18 | 36 | 43 | 102 | 61 | 138 |
| ***Heriades carinatus*** | 35 | 88 | 33 | 78 | 51 | 163 | 119 | 329 |
| ***Hylaeus annulatus*** | 0 | 0 | 5 | 8 | 34 | 83 | 39 | 91 |
| ***Megachile angelarum*** | 25 | 60 | 0 | 0 | 32 | 69 | 57 | 129 |
| ***Megachile campanulae*** | 26 | 62 | 8 | 29 | 45 | 134 | 79 | 225 |
| ***Megachile centuncularis*** | 47 | 124 | 25 | 55 | 49 | 155 | 121 | 334 |
| ***Megachile lapponica*** | 0 | 0 | 6 | 11 | 15 | 35 | 21 | 46 |
| ***Megachile mendica*** | 42 | 113 | 27 | 50 | 48 | 148 | 117 | 311 |
| ***Megachile pugnata*** | 21 | 52 | 0 | 0 | 38 | 109 | 59 | 161 |
| ***Megachile relativa*** | 39 | 104 | 31 | 76 | 45 | 129 | 115 | 309 |
| ***Megachile rotundata*** | 49 | 140 | 17 | 44 | 55 | 186 | 121 | 370 |
| ***Osmia caerulescens*** | 30 | 79 | 30 | 74 | 52 | 146 | 112 | 299 |
| ***Osmia coloradensis*** | 2 | 2 | 24 | 57 | 48 | 133 | 74 | 192 |
| ***Osmia dolerosa*** | 13 | 32 | 30 | 60 | 30 | 60 | 73 | 152 |
| ***Osmia lignaria*** | 37 | 102 | 31 | 77 | 58 | 161 | 126 | 340 |
| ***Osmia pumila*** | 24 | 49 | 33 | 87 | 56 | 165 | 113 | 301 |
| ***Osmia taurus*** | 40 | 107 | 22 | 43 | 52 | 168 | 114 | 318 |
| ***Osmia tersula*** | 31 | 60 | 21 | 41 | 39 | 87 | 91 | 188 |

**Supplementary Table 8:** References used to determine natural history of cavity-nesting bees including guild (pollinator/parasite), range, and foraging preferences.

| Family | Species | Natural History Reference |
|---|---|---|
| **Colletidae** | *Hylaeus annulatus* | (Romankova 2007) |
| **Colletidae** | *Hylaeus pictipes* | (Gibbs and Dathe 2017) |
| **Megachilidae** | *Chelostoma rapunculi* | (Buck et al. 2005) |
| **Megachilidae** | *Coelioxys funeraria* | (Scott et al. 2000) |
| **Megachilidae** | *Coelioxys modesta* | (Krombein 1967, O'Neill and O'Neill 2018) |
| **Megachilidae** | *Coelioxys moesta* | (Scott et al. 2000) |
| **Megachilidae** | *Coelioxys sayi* | (Krombein 1967) |
| **Megachilidae** | *Heriades carinatus* | (O'Neill and O'Neill 2010) |
| **Megachilidae** | *Hoplitis albifrons* | (Michener 1947) |
| **Megachilidae** | *Megachile angelarum* | (Sheffield et al. 2011) |
| **Megachilidae** | *Megachile campanulae* | (Sheffield et al. 2011) |
| **Megachilidae** | *Megachile centuncularis* | (Sheffield et al. 2011) |
| **Megachilidae** | *Megachile lapponica* | (Sheffield et al. 2011) |
| **Megachilidae** | *Megachile mendica* | (Sheffield et al. 2011) |
| **Megachilidae** | *Megachile pugnata* | (Sheffield et al. 2011) |
| **Megachilidae** | *Megachile relativa* | (Sheffield et al. 2011) |
| **Megachilidae** | *Megachile rotundata* | (Sheffield et al. 2011) |
| **Megachilidae** | *Megachile snowi* | (Bzdyk 2012) |
| **Megachilidae** | *Osmia albiventris* | (Cane et al. 2007) |
| **Megachilidae** | *Osmia bicornis bicornis* | (Everaars et al. 2011) |
| **Megachilidae** | *Osmia caerulescens* | (Gibbs et al. 2017) |
| **Megachilidae** | *Osmia coloradensis* | (Cane et al. 2007) |
| **Megachilidae** | *Osmia dolerosa* | (Cane et al. 2007) |
| **Megachilidae** | *Osmia lignaria* | (Cane et al. 2007) |
| **Megachilidae** | *Osmia pumila* | (Cane et al. 2007) |
| **Megachilidae** | *Osmia taurus* | (Gibbs et al. 2017) |
| **Megachilidae** | *Osmia tersula* | (Cane et al. 2007) |
| **Megachilidae** | *Stelis coarctatus* | (Gibbs et al. 2017) |

**Supplementary Table 9:** Plant-pollinator network metrices examined with all pollinators at a minimum read detection of 1000 for overall and landscape-specific networks. Crop, forest, and urban network metrices are at the plant family level.

| Network measure | Overall (plant family) | Overall (plant genus) | Crop Landscape | Forest Landscape | Urban Landscape |
|---|---|---|---|---|---|
| **Number of pollinators** | 18 | 18 | 15 | 16 | 18 |
| **Number of plants** | 82 | 341 | 63 | 59 | 77 |
| **Links per species** | 8.720 | 7.237 | 5.910 | 4.813 | 8.316 |
| **Connectance** | 0.591 | 0.423 | 0.488 | 0.382 | 0.570 |
| **Nestedness** | 19.714 | 18.908 | 11.731 | 17.364 | 19.057 |
| **Web asymmetry** | -0.640 | -0.900 | -0.615 | -0.573 | -0.621 |
| **Cluster coefficient** | 0.640 | 0.458 | 0.492 | 0.415 | 0.6034 |
| **Fisher alpha** | 228.862 | 1131.788 | 166.512 | 181.369 | 236.528 |

**Supplementary Figure 2:** Maps of bee species with expanded ranges in comparison to discoverlife.org and GBIF. a-c) non-native species and d-j) native species. Black shading shows previous ranges from GBIF, and green circles indicate sites where each species was found in this study. *Chelostoma rapunculi* is native to the Palearctic range, but has previously been found in Ontario, Canada (Buck et al. 2005). In addition to four Ontario detections, we found *C. rapunculi* at one site in British Columbia (Duncan, BC: Figure 2a). *Hylaeus pictipes* is native to Europe and has previously been found in Ontario (Gibbs and Dathe 2017), but we found one detection in southern Manitoba (Elie, MB: Figure 2b). *Osmia taurus* is native to Asia but has been accidentally shipped to the United States for agricultural purposes in place of *O. cornifrons* (Gibbs et al. 2017). While this species has previously been detected in Ontario, we found it in British Columbia, Alberta, Manitoba, and Prince Edward Island as well as in Ontario (43 sites Canada-wide: Figure 2c). *Coelioxys modesta* is native to eastern Canada and the U.S. (O'Neill and O'Neill 2018), but we detected it in British Columbia (Kamloops, BC, Kelowna, BC, Nanoose Bay, BC) in addition to Ontario (Figure 2d). *Megachile campanulae* is native to eastern Canada and the U.S. (Sheffield et al. 2011), but we detected it in British Columbia, Alberta, and Manitoba (eight sites), in addition to Ontario (Figure 2e). *Megachile angelarum* is native to western Canada and the U.S. (Sheffield et al. 2011), but we detected it in Ontario (eight sites), in addition to British Columbia (Figure 2f). *Megachile snowi* is native to western Canada and the U.S. (Bzdyk 2012), but we detected it in Manitoba and Ontario, in addition to British Columbia (Figure 2g). *Osmia dolerosa* is native to western Canada and the U.S. (Cane et al. 2007), but we found it in Ontario (Milton, ON, Phelpston, ON, Renfrew, ON), in addition to British Columbia (Figure 2h). *Megachile mendica* was found in northern Alberta and Manitoba when it had previously only been reported in the south of these provinces (High Level, AB, Churchill, MB: Figure 2i; Sheffield et al. 2011). *Osmia caerulescens* is native to Canada (Gibbs et al. 2017) but was found in northern Manitoba for the first time (Churchill, MB: Figure 2j).

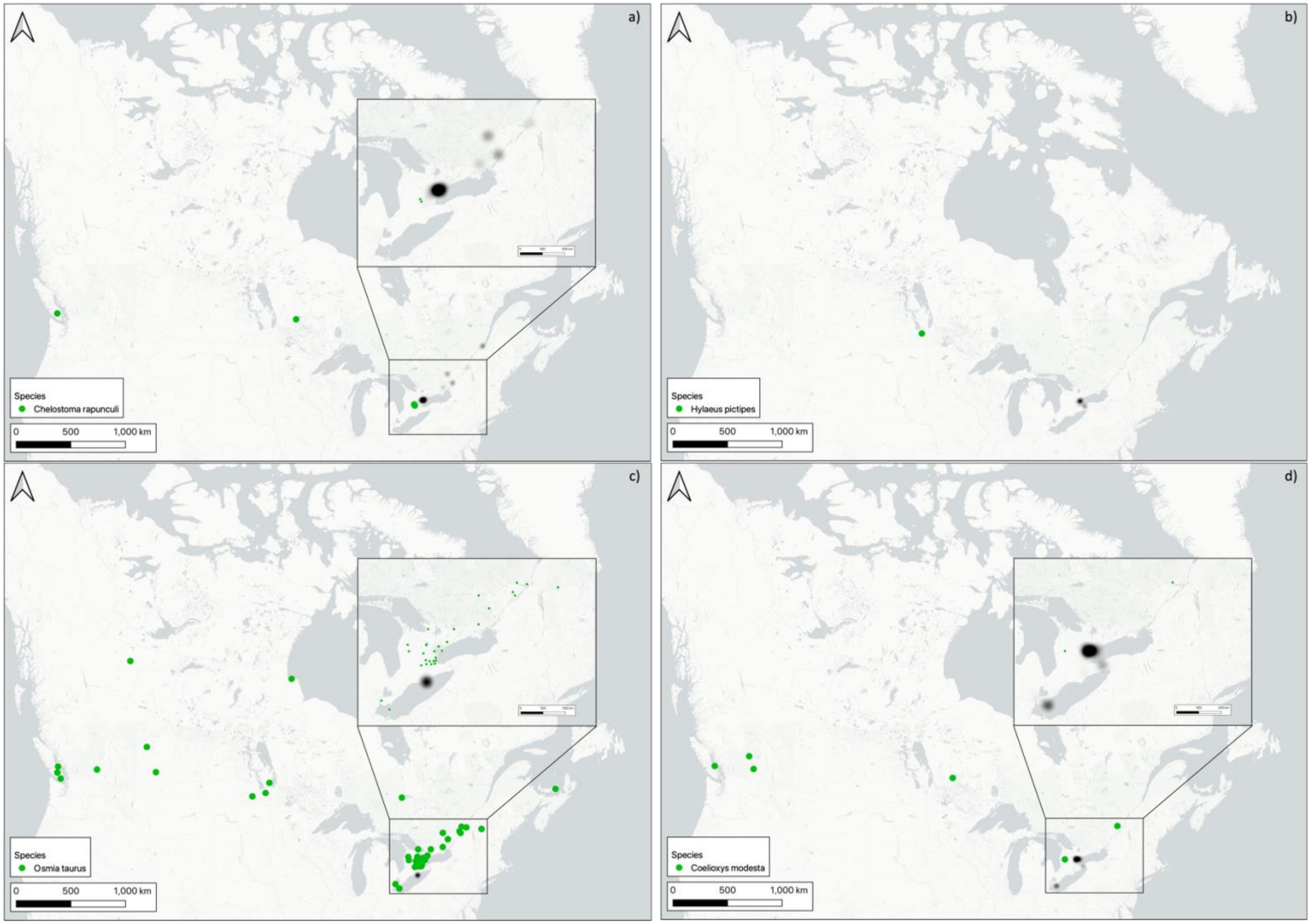
a)
Species
Chelostoma rapunculi
0 500 1,000 km
b)
Species
Hylaeus pictipes
0 500 1,000 km
c)
Species
Osmia taurus
0 500 1,000 km
d)
Species
Coelioxys modesta
0 500 1,000 km

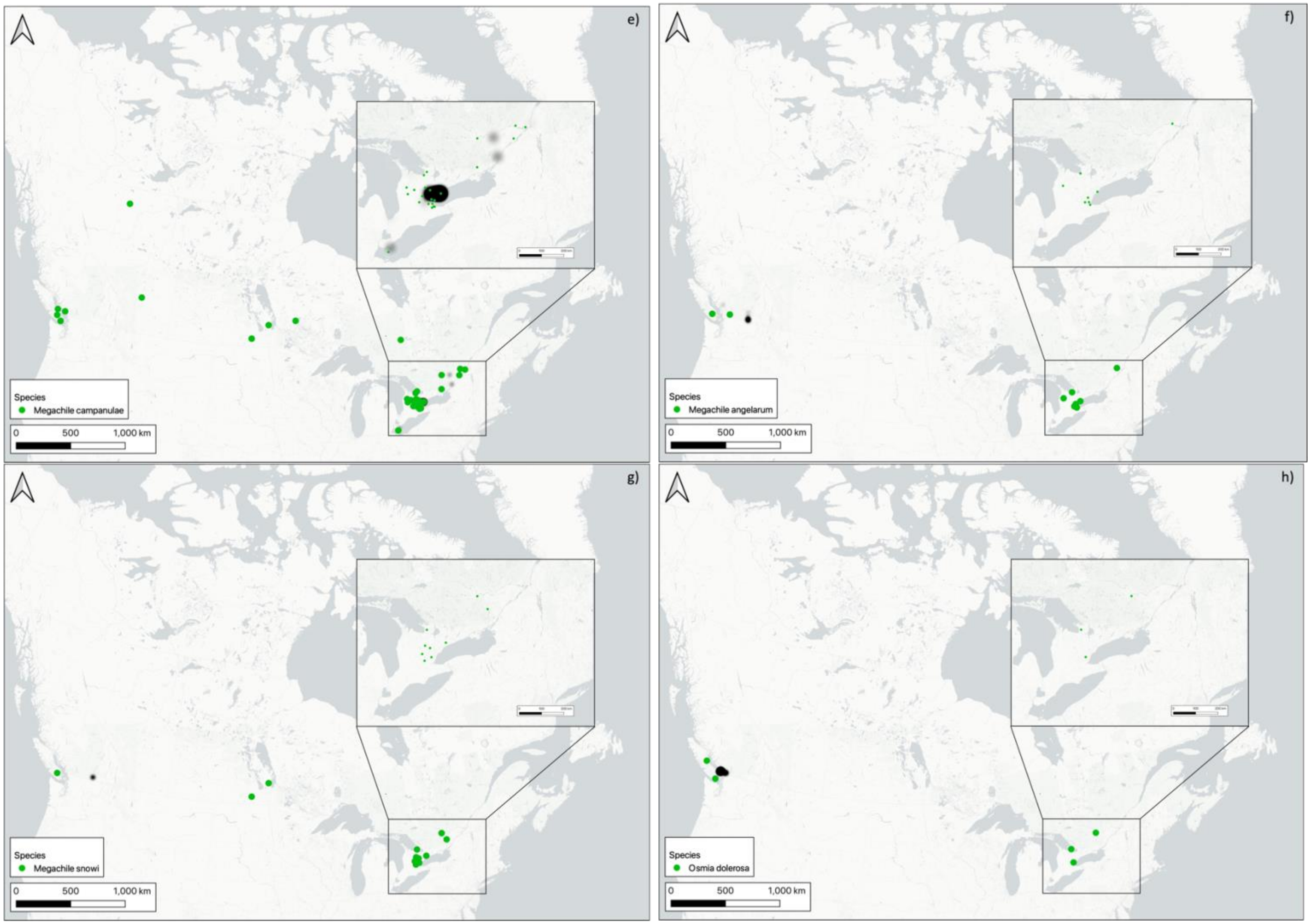
e)
Species
Megachile campanulae
0 500 1,000 km
f)
Species
Megachile angelarum
0 500 1,000 km
g)
Species
Megachile snowi
0 500 1,000 km
h)
Species
Osmia dolerosa
0 500 1,000 km

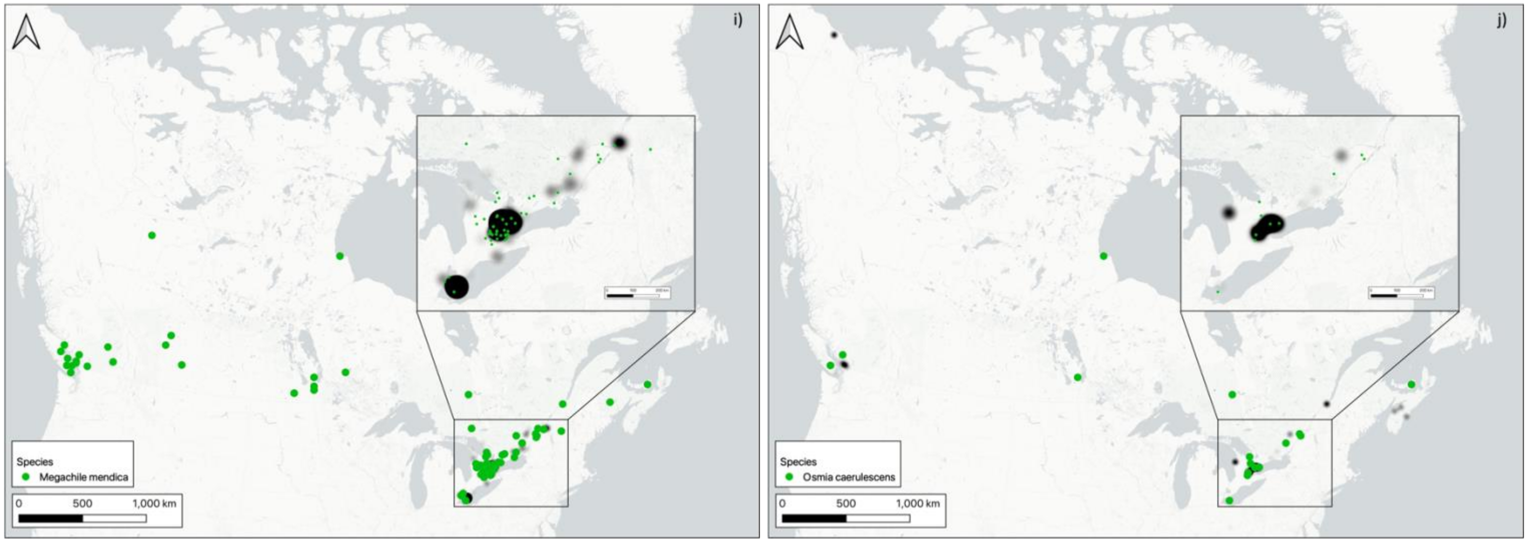

i)
Species
Megachile mendica
0 500 1,000 km
j)
Species
Osmia caerulescens
0 500 1,000 km

**Supplementary Figure 3:** Maps of bee species with ranges comparable to existing records in comparison to discoverlife.org and GBIF. Black shading shows previous ranges from GBIF, and green circles indicate sites where each species was found in this study.

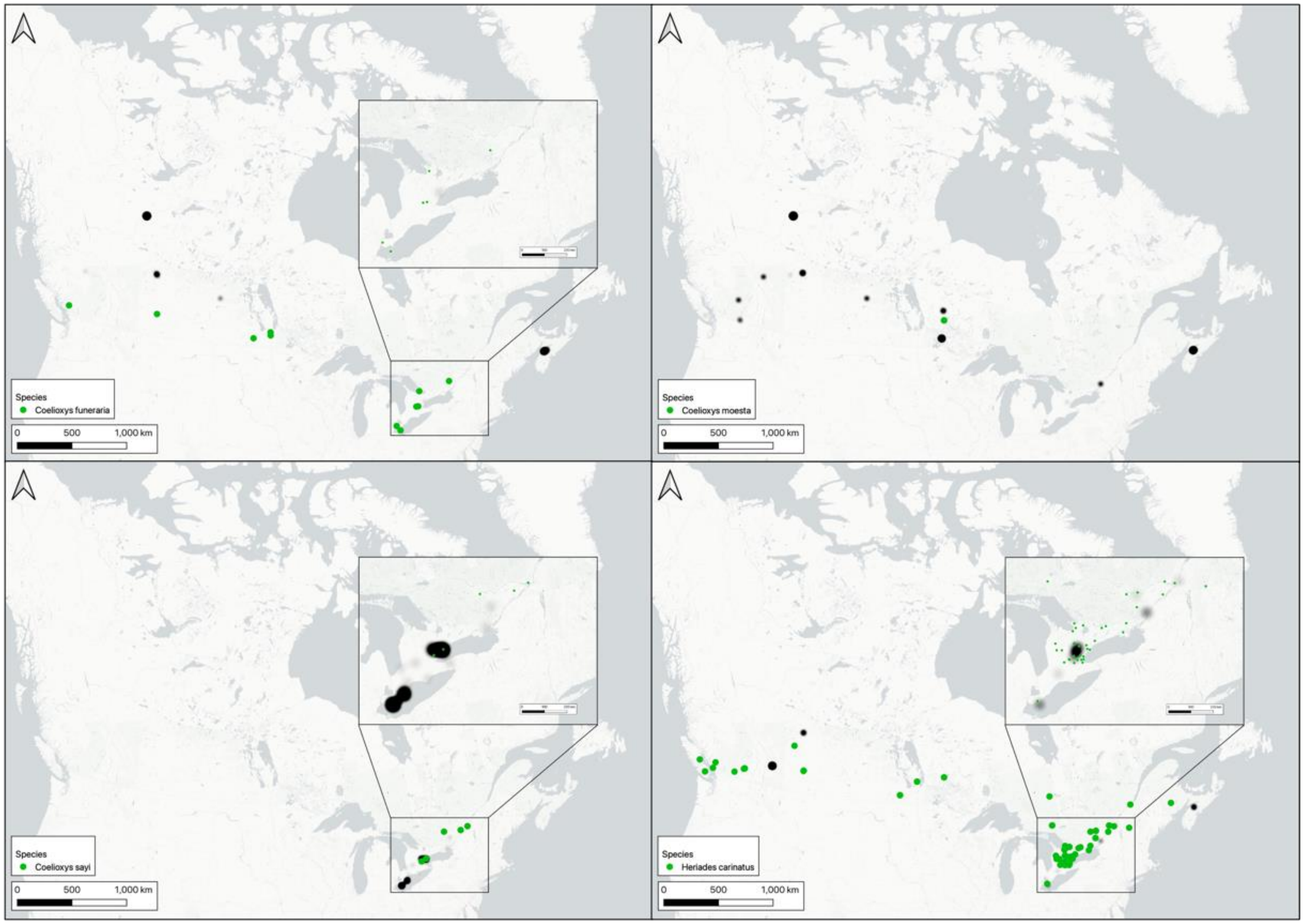

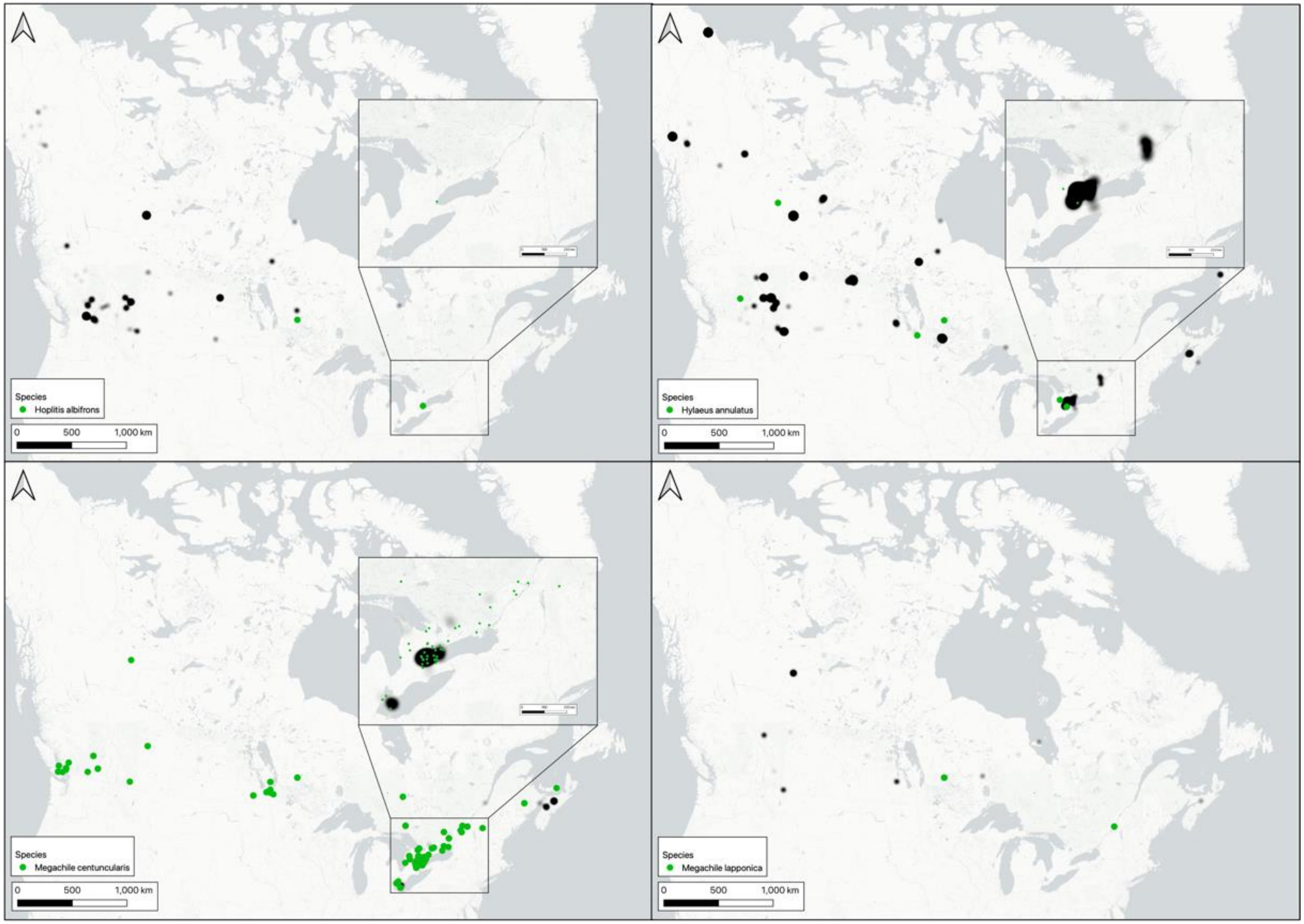
Species
Hoplitis albifrons
0
500
1,000 km
Species
Hylaeus annulatus
0
500
1,000 km
Species
Megachile centuncularis
0
500
1,000 km
Species
Megachile lapponica
0
500
1,000 km

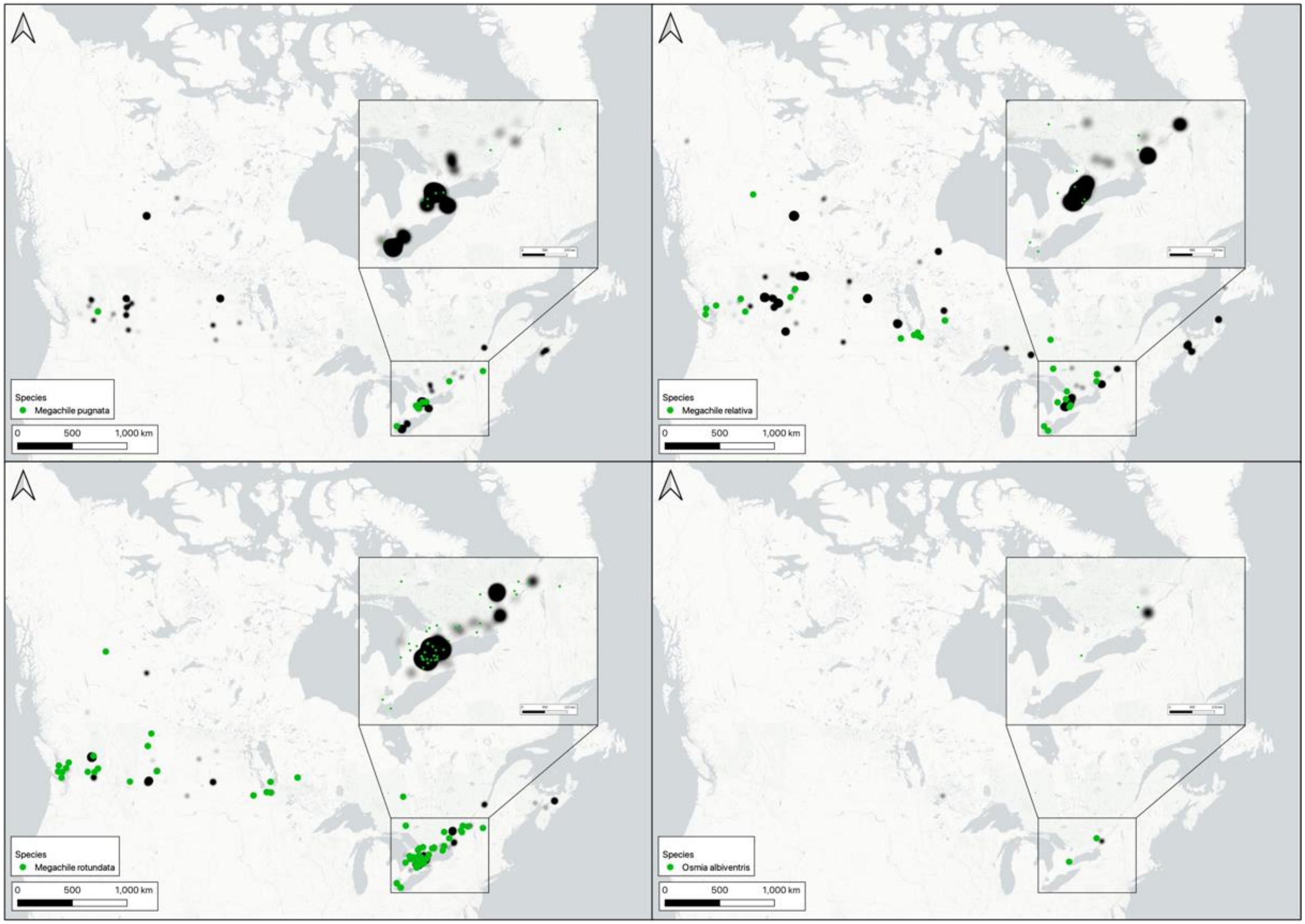
Species
Megachile pugnata
0 500 1,000 km
Species
Megachile relativa
0 500 1,000 km
Species
Megachile rotundata
0 500 1,000 km
Species
Osmia albiventris
0 500 1,000 km

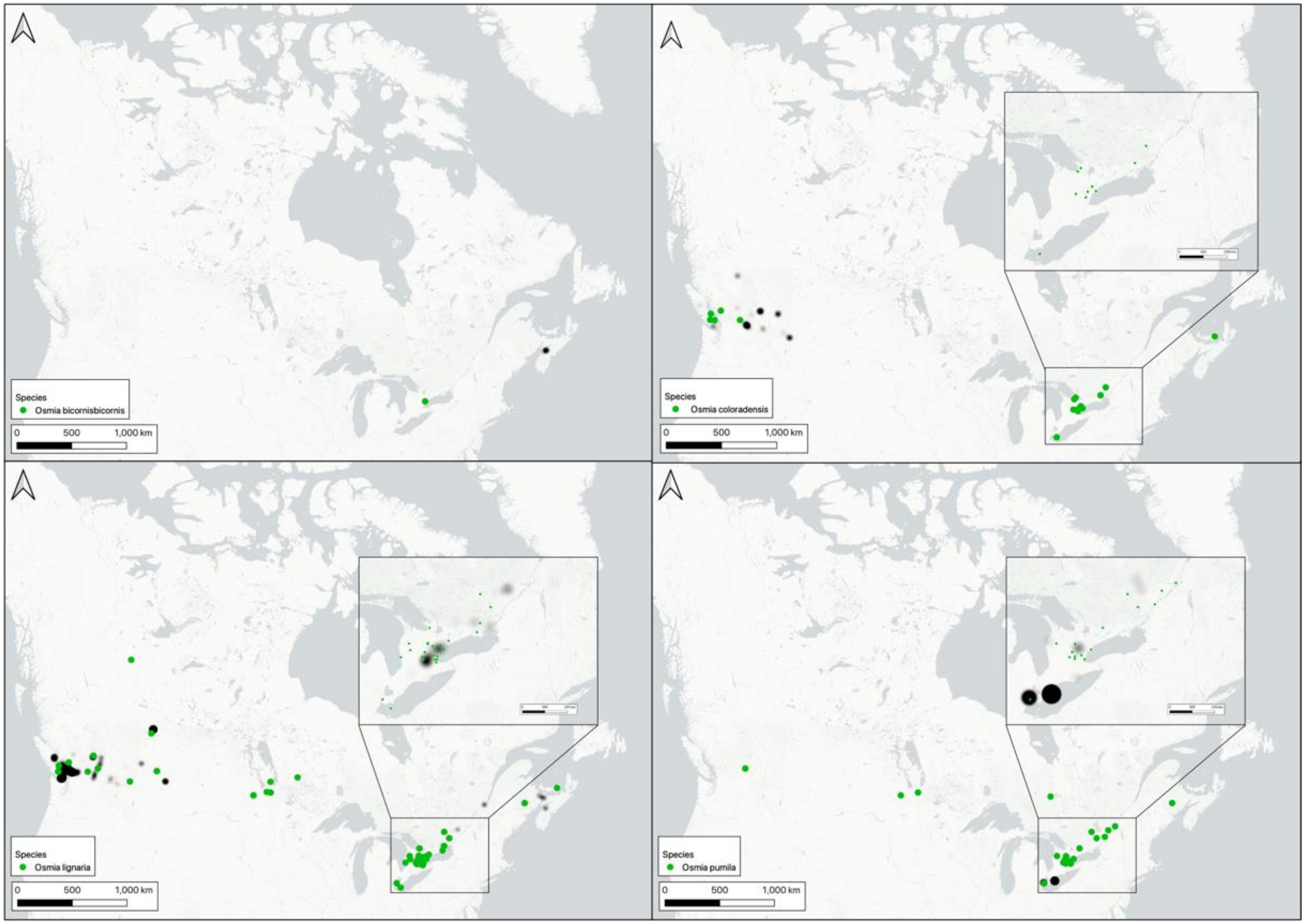
Species
Osmia bicornisbicornis
0 500 1,000 km
Species
Osmia coloradensis
0 500 1,000 km
Species
Osmia lignaria
0 500 1,000 km
Species
Osmia pumila
0 500 1,000 km

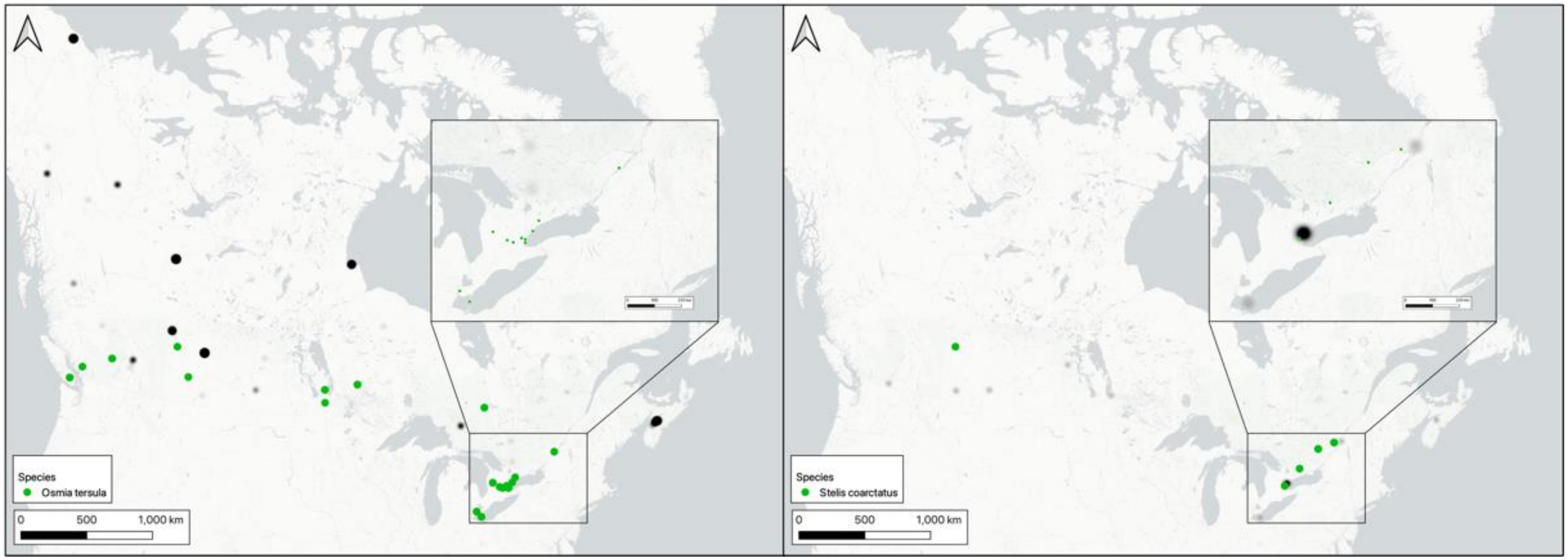
Species
Osmia tersula
0 500 1,000 km
Species
Stelis coarctatus
0 500 1,000 km

**Supplementary Figure 4**: Maps of wasp species with expanded ranges in comparison to discoverlife.org and GBIF. Black shading shows previous ranges from GBIF, and green circles indicate sites where each species was found in this study. *Passaloecus eremita* (Figure 4a), *Passaloecus gracilis* (Figure 4b) are both non-native species that have previously been found in Ontario. In addition to Ontario, we found one detection for each species (Kelowna, BC and Richmond, BC respectively) in British Columbia. *Psenulus pallipes* is non-native but has previously been found in Ontario (Figure 4c), however we found additional records in British Columbia (four sites) (Schmid-Egger 2016). Coville (1982) treats *Trypoxylon nitidum* as a species complex found in southeastern Canada. We found *Trypoxylon* cf. *nitidum* (Barcode Index Number: BOLD:ACF3990) in eastern Canada as well as British Columbia and Alberta (12 sites: Figure 4d). Voucher specimens with this Barcode Index Number (BIN) have been photographed and sent for verification by a global expert wasp taxonomist who specializes in this taxon.

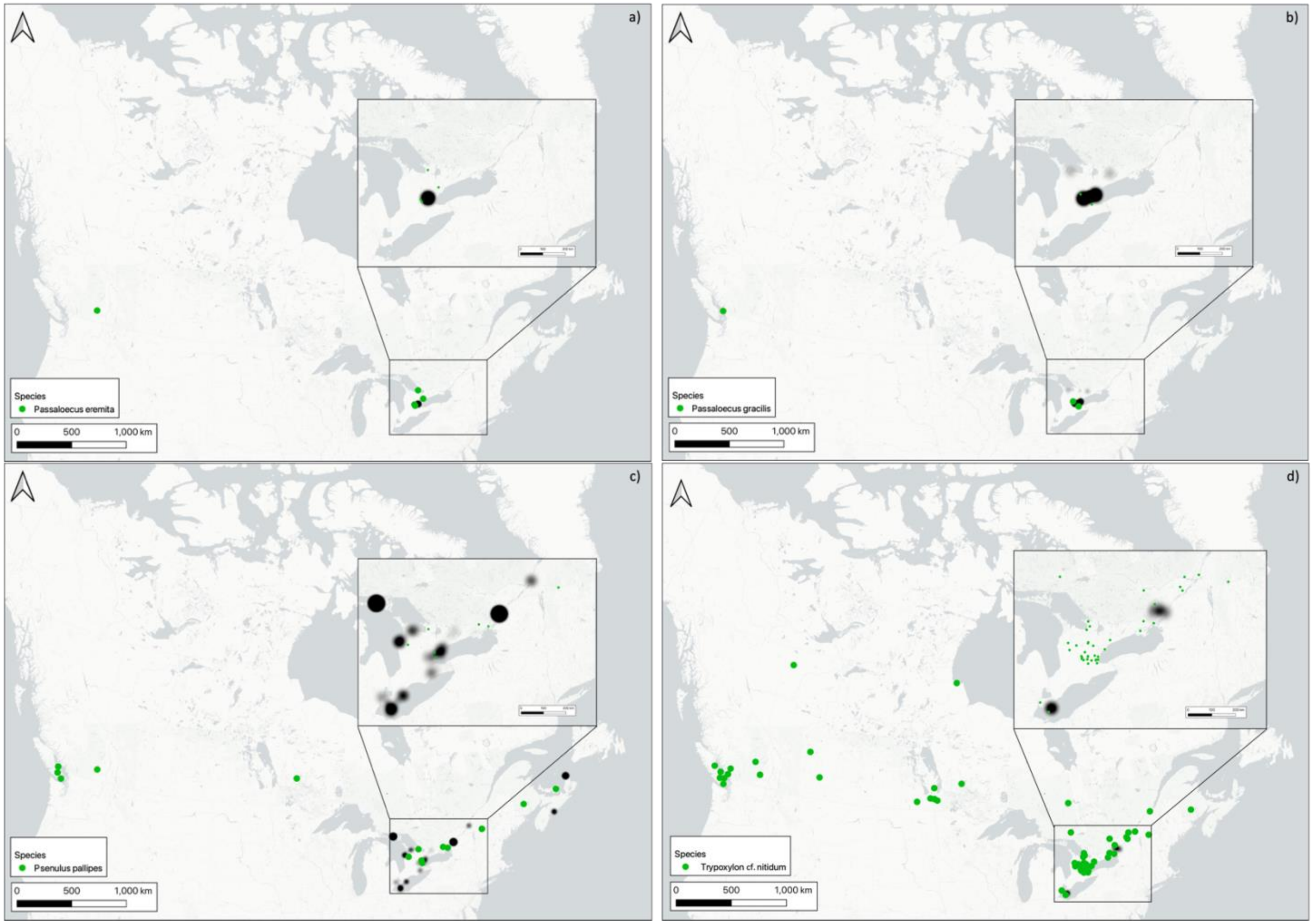
a)
Species
Passaloecus eremita
0 500 1,000 km
b)
Species
Passaloecus gracilis
0 500 1,000 km
c)
Species
Psenulus pallipes
0 500 1,000 km
d)
Species
Trypoxylon cf. nitidum
0 500 1,000 km

**Supplementary Table 10:** Predatory wasps found in trap nests along with prey. Number of observations is shown in brackets. Bold observations are those not found in historical records.

| Predator Family/Order | Predator Species | Prey Family | Prey Taxa |
|---|---|---|---|
| Crabronidae (Hymenoptera) | *Passaloecus cuspidatus*[1] | Aphididae | *Uroleucon sonchi* (1) |
| | *Passaloecus eremita*[2] | Lachnidae | *Schizolachnus obscurus* (1) |
| | *Psenulus pallipes*[3] | Aphididae | *Acyrthosiphon pisum* (2) |
| | | | *Acyrthosiphon* sp. (1) |
| | | | *Macrosiphum euphorbiae* (1) |
| | | | *Myzus persicae* (2) |
| | *Trypoxylon* cf. *nitidum*[4] | Araneidae | *Araneus diadematus* (3) |
| | | | *Eustala anastera* (1) |
| | | | *Larinioides patagiatus* (1) |
| | | Chrysomelidae | ***Chrysomela aeneicollis*** (1) |
| | | | ***Phratora purpurea*** (1) |
| | | | ***Plagiodera versicolora*** (2) |
| | | Gryllidae | ***Oecanthus nigricornis*** (1) |
| | | | ***Oecanthus niveus*** (1) |
| | | Tettigoniidae | ***Orchelimum gladiator*** (1) |
| | | Theridiidae | *Parasteatoda tepidariorum* (5) |
| | | | *Theridion murarium* (6) |
| | | | *Wamba crispulus* (1) |
| | | Tortricidae | ***Choristoneura rosaceana*** (1) |
| | *Trypoxylon figulus*[5] | Araneidae | *Larinioides patagiatus* (2) |
| | *Trypoxylon frigidum*[1] | Araneidae | *Larinioides patagiatus* (2) |
| | | | *Zygiella atrica* (1) |
| | | Pholcidae | *Pholcus manueli* (1) |
| | | Theridiidae | *Parasteatoda tabulata* (3) |
| | | | *Parasteatoda tepidariorum* (1) |
| Ichneumonidae (Hymenoptera) | *Zatypota percontatoria*[6] | Araneidae | *Eustala anastera* (1) |
| Pompilidae (Hymenoptera) | *Auplopus nigrellus*[7,8] | Crysomelidae | ***Plagiodera versicolora*** (1) |
| Sphecidae (Hymenoptera) | *Isodontia mexicana*[1,9] | Baetidae | ***Baetis intercalaris*** (1) |
| | | Chrysomelidae | ***Plagiodera versicolora*** (1) |
| | | Cicadellidae | **Deltocephalinae** (1) |
| | | Gryllidae | *Oecanthus celerinictus* (1) |
| | | | *Oecanthus nigricornis* (6) |
| | | | *Oecanthus niveus* (18) |
| | | Tettigoniidae | *Conocephalus brevipennis* (1) |
| | | | *Meconema* sp. (2) |
| | | | *Meconema thalassinum* (2) |
| | | | *Orchelimum gladiator* (3) |
| | | Theridiidae | ***Parasteatoda tepidariorum*** (1) |
| | | | ***Theridion murarium*** (1) |
| | | Tortricidae | ***Epinotia radicana*** (1) |
| Vespidae (Hymenoptera) | *Ancistrocerus adiabatus*[10,11] | Gryllidae | ***Oecanthus niveus*** (1) |
| | | Tortricidae | *Ancylis spiraeifoliana* (1) |

| Predator Family/Order | Predator Species | Prey Family | Prey Taxa |
|---|---|---|---|
| | *Ancistrocerus albophaleratus*[1,10,12] | Blastobasidae | *Holcocera chalcofrontella* (1) |
| | | Chrysomelidae | ***Plagiodera versicolora*** (2) |
| | | Gelechiidae | *Scrobipalpa* sp. (1) |
| | | Tortricidae | *Acleris albicomana* (1) |
| | | | *Acleris* sp. (1) |
| | | | *Epinotia radicana* (1) |
| | *Ancistrocerus antilope*[1,10] | Amphisbatidae | *Psilocorsis reflexella* (1) |
| | | Chrysomelidae | ***Phratora purpurea*** (1) |
| | | | ***Plagiodera versicolora*** (5) |
| | | Gryllidae | *Oecanthus niveus* (1) |
| | | Pyralidae | *Sciota subcaesiella* (1) |
| | | Tortricidae | *Choristoneura rosaceana* (2) |
| | *Ancistrocerus catskill*[1,10] | Tortricidae | *Pseudexentera cressoniana* (1) |
| | *Ancistrocerus gazella*[11] | Cosmopterigidae | *Limnaecia phragmitella* (1) |
| | *Euodynerus foraminatus*[11] | Tortricidae | *Acleris forbesana* (1) |
| | | | *Olethreutes nigranum* (1) |
| | | | *Olethreutes permundana* (1) |
| | | | *Olethreutes sp.* (1) |
| | *Euodynerus* sp.[11] | Tortricidae | *Choristoneura fumiferana* (1) |
| | *Symmorphus bifasciatus*[13] | Araenidae | ***Eustala anastera*** (1) |
| | | Baetidae | ***Baetis intercalaris*** (1) |
| | | Bucculatricidae | **Bucculatricidae** (2) |
| | | Chrysomelidae | *Chrysomela aeneicollis* (1) |
| | | | *Gastrophysa polygoni* (6) |
| | | | *Phratora purpurea* (12) |
| | | | *Plagiodera versicolora* (34) |
| | | Depressariidae | ***Machimia tentoriferella*** (2) |
| | | Gryllidae | ***Oecanthus nigricornis*** (1) |
| | | | ***Oecanthus niveus*** (1) |
| | | Theridiidae | ***Theridion murarium*** (1) |
| | | Thripidae | ***Thrips tabaci*** (1) |
| | *Symmorphus canadensis*[1,10,11] | Bucculatricidae | Bucculatricidae (2) |
| | | Chrysomelidae | *Plagiodera versicolora* (1) |
| | | Curculionidae | *Isochnus sequensi* (4) |
| | | Gracillariidae | *Caloptilia burgessiella* (1) |
| | | Gryllidae | ***Oecanthus niveus*** (1) |
| | | Heliozelidae | *Antispila cornifoliella* (1) |
| | | | *Antispila freemani* (1) |
| | *Symmorphus cristatus*[1,10,11] | Chrysomelidae | *Chrysomela aeneicollis* (1) |
| | | | *Chrysomela scripta* (1) |
| | | | *Chrysomela* sp. (1) |
| | | | *Phratora purpurea* (1) |
| | | | *Plagiodera versicolora* (7) |
| | | Theridiidae | ***Theridion murarium*** (1) |
| | *Symmorphus* sp.[1,10] | Bucculatricidae | *Bucculatrix ainsliella* (1) |

1. Krombein, K. V. 1967. Trap-nesting wasps and bees life histories nests and associates. Smithsonian Press.
2. Lomholdt, O. 1984. The Sphecidae (Hymenoptera) of Fennoscandia and Denmark.

3. Schmid-Egger, C. 2016. The *Psenulus pallipes* species group in Central Europe (Hymenoptera, Crabronidae). Ampulex 8:40–44.
4. Scher, R., and S. das Graças Pompolo. 2003. Evolutionary dynamics of the karyotype of the wasp *Trypoxylon* (*Trypargilum*) *nitidum* (Hymenoptera, Sphecidae) from the Rio Doce State Park, Minas Gerais, Brazil. Genetics and Molecular Biology 26:307–311.
5. Coudrain, V., F. Herzog, and M. H. Entling. 2013. Effects of habitat fragmentation on abundance, larval food and parasitism of a spider-hunting wasp. PLoS ONE 8.
6. Korenko, S., V. Michalková, K. Zwakhals, and S. Pekr. 2011. Host specificity and temporal and seasonal shifts in host preference of a web-spider parasitoid Zatypota percontatoria. Journal of Insect Science 11:1–12.
7. Conrow, R. T., K. M. Zivicki, and G. P. Setliff. 2016. Cuckoo wasps of Pennsylvania (Hymenoptera: Chrysididae). Transactions of the American Entomological Society 142:113–129.
8. Townes, H. 1957. Nearctic wasps of the subfamilies Pepsinae and Ceropalinae. Bulletin of the United States National Museum:1–286.
9. O'Hara, J. E. 2005. A review of the tachinid parasitoids (Diptera: Tachinidae) of Nearctic *Choristoneura* species (Lepidoptera: Tortricidae), with keys to adults and puparia. Zootaxa 938:1–46.
10. Krombein, K. V., and P. D. Hurd. 1979. Catalog of Hymenoptera in America North of Mexico.
11. Buck, M., S. A. Marshall, and D. K. B. Cheung. 2008. Identification atlas of the Vespidae (Hymenoptera, Aculeata) of the northeastern Nearctic region. Canadian Journal of Arthropod Identification 5.
12. Fye, R. E. 1965. Biology of Apoidea taken in trap nests in northwestern Ontario (Hymenoptera). The Canadian Entomologist 97:863–877.
13. Budrienė, A. 2003. Prey of *Symmorphus* wasps (Hymenoptera: Eumeninae) in Lithuania. Acta Zoologica Lituanica 13:306–310.

**Supplementary Table 11:** Parasitic wasps, flies, and beetles found in cavity-nest boxes along with hosts. Number of observations is shown in brackets. Bold observations are those not found in historical records.

| Natural Enemy Family/Order | Natural Enemy Species | Host Family | Host Species |
|---|---|---|---|
| Bombyliidae (Diptera) | *Anthrax irroratus*[1] | Colletidae | *Hylaeus annulatus* (1) |
| | | Megachilidae | *Megachile campanulae* (2) |
| | | | *Megachile lapponica* (2) |
| | | | *Megachile relativa* (4) |
| | | | *Megachile rotundata* (1) |
| | | Vespidae | *Ancistrocerus albophaleratus* (1) |
| | *Anthrax* sp.[1] | Megachilidae | *Heriades carinatus* (1) |
| Braconidae (Hymenoptera) | *Meteorus* sp.[2] | Vespidae | *Symmorphus bifasciatus* (1) |
| Chrysididae (Hymenoptera) | *Caenochrysis tridens* [3,4] | Crabronidae | ***Trypoxylon* cf. *nitidum*** (4) |
| | | Sphecidae | ***Isodontia mexicana*** (1) |
| | *Chrysis cembricola*[5] | Crabronidae | ***Trypoxylon* cf. *nitidum*** (1) |
| | | Megachilidae | ***Megachile mentica*** (1) |
| | | | ***Megachile rotundata*** (2) |
| | | Vespidae | ***Ancistrocerus gazella*** (1) |
| | | | ***Symmorphus bifasciatus*** (3) |
| | *Chrysis sp.*[5] | Crabronidae | *Trypoxylon* cf. *nitidum* (2) |
| | | Megachilidae | *Megachile relativa* (1) |
| | | Sphecidae | *Isodontia mexicana* (1) |
| | *Omalus aeneus* [5,6] | Crabronidae | *Passaloecus eremita* (1) |
| | *Pseudomalus* sp.[7] | Crabronidae | *Passaloecus eremita* (1) |
| Conopidae (Diptera) | *Zodion fulvifrons*[8] | Megachilidae | *Megachile centuncularis* (1) |
| | *Zodion intermedium*[8] | Megachilidae | *Megachile centuncularis* (1) |
| | *Zodion* sp. [8] | Megachilidae | *Megachile centuncularis* (1) |
| Eulophidae (Hymenoptera) | Eulophidae[9] | Megachilidae | *Megachile mendica* (1) |
| | *Melittobia acasta*[10] | Megachilidae | *Megachile rotundata* (1) |
| Ichneumonidae (Hymenoptera) | *Perithous* sp.[11] | Crabronidae | *Psenulus pallipes* (1) |
| Megachilidae (Hymenoptera) | *Coelioxys funeraria*[12] | Megachilidae | ***Megachile mendica*** (1) |
| | | | *Megachile relativa* (6) |
| | | | ***Osmia lignaria*** (3) |
| | *Coelioxys modesta*[5,13] | Megachilidae | ***Megachile angelarum*** (1) |
| | | | *Megachile campanulae* (2) |
| | | | *Megachile relativa* (3) |
| | | Vespidae | ***Ancistrocerus antilope*** (1) |
| | | | ***Euodynerus* sp.** (1) |
| | *Coelioxys moesta*[4,12] | Megachilidae | *Megachile relativa* (1) |
| | | | ***Megachile rotundata*** (2) |
| | *Coelioxys sayi*[5,14] | Megachilidae | ***Megachile lapponica*** (1) |
| | | | *Megachile mendica* (5) |
| | | | ***Osmia pumila*** (1) |
| | | Sphecidae | ***Isodontia mexicana*** (1) |
| Megachilidae (Hymenoptera) | *Stelis coarctatus*[15] | Crabronidae | ***Trypoxylon* cf. *nitidum*** (1) |
| | | Megachilidae | *Heriades carinatus* (6) |
| | | | ***Megachile campanluae*** (1) |

| Natural Enemy Family/Order | Natural Enemy Species | Host Family | Host Species |
|---|---|---|---|
| | | Sphecidae | ***Isodontia mexicana*** (1) |
| Meloidae (Coleoptera) | *Nemognatha* sp.[16] | Megachilidae | *Megachile rotundata* (1) |
| | | Sphecidae | *Isodontia mexicana* (1) |
| | | Vespidae | *Symmorphus canadensis* (1) |
| Phoridae (Diptera) | *Apocephalus borealis*[17] | Megachilidae | *Megachile rotundata* (1) |
| Sapygidae (Hymenoptera) | *Sapyga louisi*[5] | Megachilidae | *Heriades carinatus* (5) |
| | | | ***Megachile angelarum*** (1) |
| | | | ***Megachile campanulae*** (2) |
| | | | ***Megachile mendica*** (1) |
| | | | ***Osmia lignaria*** (1) |
| | *Sapyga similis*[18,] 19 | Crabronidae | ***Trypoxylon* cf. *nitidum*** (1) |
| | | Megachilidae | *Osmia tersula* (3) |
| | *Sapyga* sp.[5] | Megachilidae | *Heriades carinatus* (1) |
| | | | *Megachile rotundata* (1) |
| | | Vespidae | *Symmorphus bifasciatus* (1) |
| Sarcophagidae (Diptera) | *Amobia* sp.[20] | Megachilidae | ***Megachile mendica*** (1) |
| | | | ***Megachile rotundata*** (1) |
| | | | ***Osmia lignaria*** (1) |
| | | Vespidae | *Ancistrocerus adiabatus* (1) |
| | | | *Symmorphus bifasciatus* (2) |
| | | | *Symmorphus canadensis* (1) |
| Tachinidae (Diptera) | *Actia diffidens*[21] | Tortricidae | *Acleris albicomana* (1) |
| | *Hemisturmia* sp.[21] | Depressariidae | *Machimia tentoriferella* (1) |
| | | Chrysomelidae | ***Plagiodera versicolora*** (1) |
| | *Lypha fumipennis*[21] | Tortricidae | *Acleris forbesana* (1) |
| | | | *Olethreutes nigranum* (1) |
| | | | *Olethreutes permundana* (1) |
| | | | *Olethreutes* sp. (1) |
| Torymidae (Hymenoptera) | *Monodontomerus montivagus*[22,23] | Crabronidae | *Trypoxylon* cf. *nitidum* (3) |
| | | | *Trypoxylon figulus* (1) |
| | | Megachilidae | *Heriades carinatus* (1) |
| | | | *Megachile centuncularis* (2) |
| | | | *Megachile relativa* (1) |
| | | | *Megachile rotundata* (16) |
| | | | *Osmia dolerosa* (1) |
| | | | *Osmia lignaria* (3) |
| | | | *Osmia pumila* (3) |
| | | | *Osmia taurus* (9) |
| | | | *Osmia tersula* (1) |
| | | Vespidae | *Euodynerus* sp. (1) |
| | | | *Symmorphus bifasciatus* (1) |
| | *Monodontomerus* sp.[22,23] | Crabronidae | *Trypoxylon* cf. *nitidum* (2) |
| | | | *Trypoxylon figulus* (1) |
| | | | *Trypoxylon frigidum* (1) |
| | | Megachilidae | *Heriades carinatus* (1) |
| | | | *Megachile angelarum* (1) |
| Torymidae (Hymenoptera) | *Monodontomerus* sp.[22,23] | Megachilidae | *Megachile campanulae* (3) |
| | | | *Megachile centuncularis* (3) |
| | | | *Megachile mendica* (3) |

| Natural Enemy Family/Order | Natural Enemy Species | Host Family | Host Species |
|---|---|---|---|
| | | | *Megachile relativa* (1) |
| | | | *Megachile rotundata* (14) |
| | | | *Osmia caerulescens* (2) |
| | | | *Osmia coloradensis* (3) |
| | | | *Osmia dolerosa* (2) |
| | | | *Osmia lignaria* (18) |
| | | | *Osmia pumila* (3) |
| | | | *Osmia taurus* (20) |
| | | Sphecidae | *Isodontia mexicana* (1) |
| | | Vespidae | *Ancistrocerus adiabatus* (1) |
| | | | *Ancistrocerus albophaleratus* (2) |
| | | | *Euodynerus sp.* (1) |
| | | Vespidae | *Symmorphus bifasciatus* (3) |
| Xenidae (Hymenoptera) | Xenidae[24] | Megachilidae | *Megachile relativa* (1) |
| | | Vespidae | *Ancistrocerus albophaleratus* (3) |


1. Scott, V. L., and K. Strickler. 1992. New host records for two species of *Anthrax* (Diptera: Bombyliidae). Journal of the Kansas Entomological Society 65:393–402.
2. Muesebeck, C. F. W. 1923. A revision of the North American species of Ichenumon-flies belonging to the genus *Meteorus* Haliday. Proceedings of the United States National Museum 63:1–44.
3. Krombein, K. V., and P. D. Hurd. 1979. Catalog of Hymenoptera in America North of Mexico.
4. Conrow, R. T., K. M. Zivicki, and G. P. Setliff. 2016. Cuckoo wasps of Pennsylvania (Hymenoptera: Chrysididae). Transactions of the American Entomological Society 142:113–129.
5. Krombein, K. V. 1967. Trap-nesting wasps and bees life histories nests and associates. Smithsonian Press.
6. Bohart, R. M. 1960. Addendum to a review of the Genus *Omalus* Panzer in North America (Hymenoptera, Chrysididae). Annals of the Entomological Society of America 53:435–435.
7. Bohart, R. M., and L. S. Kimsey. 1982. A synopsis of the Chrysididae in America North of Mexico. Memoirs of the American Entomological Institute 33:1–266.
8. Freeman, B. A. 1966. Notes on Conopid flies, including insect host, plant and phoretic relationships (Diptera: Conopidae). Journal of the Kansas Entomological Society 39:123–131.
9. Gauthier, N., J. Lasalle, D. L. J. Quicke, and H. C. J. Godfray. 2000. Phylogeny of Eulophidae (Hymenoptera: Chalcidoidea), with a reclassification of Eulophinae and the recognition that Elasmidae are derived eulophids. Systematic Entomology 25:521–539.
10. González, J. M., J. B. Terán, and R. W. Matthews. 2004. Review of the biology of *Melittobia acasta* (Walker) (Hymenoptera: Eulophidae) and additions on development and sex ratio of the species. Caribbean Journal of Science 40:52–61.
11. Tormos, J., J. D. Asís, S. F. Gayubo, and J. Selfa. 2004. Descriptions of the final instar larvae of *Perithous septemcinctorius*, *Zatypota bohemani* and *Z. gracilis* (Hymenoptera: Ichneumonidae: Pimplinae). Journal of Entomological Science 39:475–482.
12. Scott, V. L., S. T. Kelley, and K. Strickler. 2000. Reproductive biology of two *Coelioxys* cleptoparasites in relation to their *Megachile* hosts (Hymenoptera: Megachilidae). Annals of the Entomological Society of America 93:941–948.
13. O'Neill, K. M., and J. F. O'Neill. 2018. Cavity-nesting wasps and bees (Hymenoptera) of Central New York State: Finger Lakes National Forest. Proceedings of the Entomological Society of Washington 120:260–271.
14. Baker, J. R., E. D. Kuhn, and S. B. Bambara. 1985. Nests and immature stages of leafcutter bees (Hymenoptera: Megachilidae). Journal of the Kansas Entomological Society 58:290–313.
15. Gibbs, J., J. S. Ascher, M. G. Rightmyer, and R. Isaacs. 2017. The bees of Michigan (Hymenoptera: Apoidea: Anthophila), with notes on distribution, taxonomy, pollination, and natural history. Zootaxa 4352:1–160.
16. Blochtein, B., and D. Wittmann. 1988. Mating site specificity, reproduction and vector selection in *Nemognatha nigrotarsata* (Col., Meloidae), a nest parasite of leaf-cutter bees and other pollinators of crops in Rio Grande do Sul. Journal of Applied Entomology 105:414–419.
17. Khattab, M. M., and E. Nowar. 2014. The first records of the parasite Zombie Fly (*Apocephalus borealis* Brues) on honeybee, *Apis mellifera* in Egypt. International Journal of Agricultural Science and Research 4:37–42.
18. Malyshev, S. I. 1968. Genesis of the Hymenoptera and the phases of their evolution.
19. Müller, A., R. Prosi, C. Praz, and H. Richter. 2019. Nesting in bark – the peculiar life history of the rare boreoalpine osmiine bee *Osmia* (*Melanosmia*) *nigriventris* (Hymenoptera, Megachilidae). Alpine Entomology 3:105–119.
20. Spofford, M. G., F. E. Kurczewski, and W. L. Downes. 1989. Nearctic species of Miltogrammini (Diptera: Sarcophagidae) associated with species of Aculeata (Hymenoptera: Vespoidea, Pompiloidea, Sphecoidea, Apoidea). Kansas (Central States) Entomological Society 62:254–267.
21. O'Neill, K. M., and J. F. O'Neill. 2009. Prey, nest associates, and sex ratios of *Isodontia mexicana* (Saussure) (Hymenoptera: Sphecidae) from two sites in New York state. Entomologica Americana 115:90–94.
22. Farzan, S., J. A. Whitney, and L. H. Yang. 2017. The phenology and spatial distribution of cavity-nesting Hymenoptera and their parasitoids in a California oak-chaparral landscape mosaic. The American Midland Naturalist 177:84–99.
23. Huber, J. T., A. M. R. Bennett, G. A. P. Gibson, Y. M. Zhang, and D. C. Darling. 2021. Checklist of Chalcidoidea and Mymarommatoidea (Hymenoptera) of Canada, Alaska and Greenland. Journal of Hymenoptera Research 82:68–138.
24. Benda, D., K. Votýpková, Y. Nakase, and J. Straka. 2021. Unexpected cryptic species diversity of parasites of the family Xenidae (Strepsiptera) with a constant diversification rate over time. Systematic Entomology 46:252–265.
